# Ancestral Musaceae karyotype reconstruction provides insights into chromosome evolution and bract coloration

**DOI:** 10.64898/2026.05.04.721291

**Authors:** Ning Fu, Pengchuan Sun, Xin Liu, Tong-Jian Liu, Yi-Bing Wang, Wei-Ming Li, Tian-Wen Xiao, Xiao-Nan Li, Yong-Yuan Mi, Zheng-Feng Wang, Mathieu Rouard, Xue-Jun Ge, Hui-Run Huang, Xin-Feng Wang

**Affiliations:** Key Laboratory of National Forestry and Grassland Administration on Plant Conservation and Utilization in Southern China, South China Botanical Garden, Chinese Academy of Sciences, Guangzhou 510650, China; Key Laboratory for Bio-Resources and Eco-Environment, Sichuan Zoige Alpine Wetland Ecosystem National Observation and Research Station, College of Life Sciences, Sichuan University, Chengdu 610065, China; School of Marine Sciences and Biotechnology, Guangxi Minzu University, Nanning 530008, China; Bioversity International, Parc Scientifique Agropolis II, 34397 Montpellier, France; South China National Botanical Garden, Guangzhou 510650, China; The Seventh Affiliated Hospital, Southern Medical University, Foshan 528244, China; State Key Laboratory of Biocontrol, Guangdong Provincial Key Laboratory of Plant Stress Biology, School of Life Sciences, Sun Yat-sen University, Guangzhou 510275, China

**Keywords:** Musaceae, *Musa exotica*, T2T genome, Karyotype evolution, Chromosomal rearrangement, Anthocyanin biosynthesis, Transcriptional regulation

## Abstract

The banana family (Musaceae) exhibits remarkable diversity in both karyotype structure and bract coloration, yet the evolutionary dynamics of chromosomes and the genomic and regulatory basis underlying color diversification remain poorly understood. Here, we present a telomere-to-telomere (T2T), gap-free genome assembly of *Musa exotica*, an ornamental species with brightly colored bracts occupying an early-branching lineage within sect. *Callimusa* (*Musa* L.). By integrating this high-quality genome with other available Musaceae genomes, we reconstruct the ancestral Musaceae karyotype (AMK) for the first time, inferring a haploid chromosome number of *n* = 17. Comparative genomic analyses reveal recurrent, highly complex lineage-specific inter-chromosome rearrangements across extant Musaceae lineages, leading to inferred stepwise reductions in chromosome number to *n* = 11, 10, and 9. Notably, closely related species share similar rearrangement patterns, suggesting conserved evolutionary trajectories shaped by lineage-specific structural remodeling. Strikingly, rearrangement-associated regions are enriched for functionally important genes, particularly structural genes (*CHS* and *F3H*) and regulatory transcription factors (MYB and bHLH) involved in the anthocyanin biosynthesis pathway. Integrative transcriptomic and regulatory analyses further demonstrate coordinated activation of anthocyanin biosynthetic genes (*CHS*, *CHI*, *F3′5′H*, and *ANS*) in bracts, with expression divergence largely decoupled from gene dosage and predominantly driven by transcriptional regulation. Co-expression analyses reveal extensive MYB– and bHLH–enzyme interactions, underscoring their central role in modulating pathway activity and bract coloration diversity. Collectively, our findings suggest a link between genome structural evolution to trait diversification, offering a refined framework for understanding genome evolution and phenotypic diversification in Musaceae and other monocots.

**Significance:** We reconstructed the ancestral Musaceae karyotype and revealed extensive lineage-specific chromosome rearrangements underlying karyotype evolution. Rearrangement-associated regions are enriched for anthocyanin biosynthetic and regulatory genes, suggesting that genome structural evolution may have contributed to bract coloration diversification in Musaceae. Integrative transcriptomic analyses further indicate that variation in anthocyanin-mediated bract coloration is more closely associated with transcriptional regulation than with gene dosage alone.

## Introduction

The banana family (Musaceae) represents an important lineage within the order Zingiberales, encompassing approximately 80 species across three genera, *Musa* L., *Ensete* Horan., and *Musella* (Fr.) C.Y. Wu [1–3]. Among them, *Musa* includes ∼70 species and has traditionally been divided into two major sections, sect. *Musa* and sect. *Callimusa*, partly based on differences in chromosome number (*n* = 11 versus *n* = 10/9/7, respectively) [4–6]. In contrast, both *Ensete* and *Musella* exhibit a chromosome number of *n* = 9 [7], overlapping with that observed in sect. *Callimusa*. This convergence highlights a fundamental limitation of chromosome number alone in resolving evolutionary relationships, as species with identical chromosome counts may harbor substantially different chromosomal architectures. Understanding how such structural variation arises and contributes to lineage divergence remains a central challenge in plant evolutionary genomics.

Chromosomal rearrangements, including translocations, fusions, and fissions, are major drivers of genome evolution and karyotype diversification [8–10]. Reconstruction of ancestral karyotypes provides a powerful framework for tracing these structural changes and inferring the evolutionary trajectories of extant genomes [8, 9]. Because identical chromosomal rearrangements are unlikely to arise independently in separate lineages, shared structural features can provide robust phylogenetic signals and insights into genome evolution [11–16]. In plants, ancient whole-genome duplications (WGDs) are often followed by extensive chromosomal reshuffling, leading to substantial variation in chromosome number and organization [17, 18]. In Musaceae, three ancient WGD events (α, β, and γ) have been identified [19, 20]; however, how subsequent chromosomal rearrangements shaped the divergence of karyotypes across lineages remains largely unresolved. In particular, the lack of a well-defined ancestral Musaceae karyotype has hindered efforts to reconstruct the stepwise processes underlying chromosome evolution in this family.

Beyond structural evolution, Musaceae species also exhibit striking phenotypic diversity, particularly in bract coloration, which ranges from dull green to vivid red, orange, and pink [3, 21, 22]. These color variations are largely determined by anthocyanins, a class of flavonoid pigments synthesized through a well-characterized metabolic pathway [23]. While the core biosynthetic genes are broadly conserved, the extent to which gene copy number variation, transcriptional regulation, and genome structural variation contribute to pigmentation diversity remains unclear. In particular, whether large-scale chromosomal rearrangements influence the organization and regulation of anthocyanin biosynthesis genes has not been systematically investigated.

Addressing these questions requires high-quality genome assemblies within a robust comparative framework. Although several Musaceae genomes are available [24], only a few have reached T2T or near-T2T quality [25–29], and high-quality assemblies from sect. *Callimusa* remain scarce, limiting comprehensive reconstruction of ancestral karyotypes and genome evolution. Here, we present a telomere-to-telomere (T2T), gap-free genome assembly of *Musa exotica* (*n* = *x* = 10), a basal species of sect. *Callimusa* with distinctive brightly colored bracts [5, 21, 30, 31]. Leveraging this high-quality genome together with other available Musaceae genomes, we reconstruct the ancestral Musaceae karyotype (AMK) for the first time and delineate the stepwise trajectory of chromosome evolution across the family. We show that lineage-specific inter-chromosomal rearrangements have driven progressive reductions in chromosome number and generated distinct karyotype architectures. Importantly, we demonstrate that rearrangement-associated regions are enriched for genes involved in anthocyanin biosynthesis and transcriptional regulation, linking genome structural evolution to phenotypic diversification. Integrating comparative genomics, gene duplication analysis, and transcriptomic profiling, we further reveal that variation in bract coloration is primarily governed by transcriptional regulation rather than gene dosage. Together, our study establishes a mechanistic framework connecting chromosomal evolution, genome organization, and trait diversification in Musaceae.

## Results

### A telomere-to-telomere genome assembly of *M. exotica* provides a high-quality Musaceae reference

The estimated genome size is 509.72 Mb with a heterozygous ratio of 0.67% based on 21-mer analysis (**Table 1 and Table S1**). The primary assembly is 595.00 Mb in length, with a contig N50 of 47.41 Mb (**Table S2–S3**), representing the highest contiguity among the 11 published *Musa* genomes (**Table S4**). After removing redundant contigs, the genome size was 567.95 Mb (**Table S3**). Using Hi-C data, 542.68 Mb of sequences were anchored onto ten pseudo-chromosomes, containing 13 telomeres and four gaps (**Tables S3 and S5**). These telomeres and gaps were subsequently patched (**Table S6**), followed by polishing with Illumina reads, resulting in a telomere-to-telomere (T2T) gap-free genome. The final chromosome lengths range from 45.51 Mb to 69.60 Mb (**Fig. 1D; Table S3**). The genome shows high completeness and accuracy, with a 98.70% BUSCO completeness, 99.62% and 95.75% mapping rates for HiFi and Illumina reads, respectively, a GC content of 39.32%, an LTR assembly index (LAI) score of 13.56 and consensus quality value (QV) of 36.83, respectively (**Table 1; Tables S4 and S7**). A total of 38,669 gene models (43,757 transcripts) were predicted, with 96.22% of embryophyta BUSCO genes were successfully annotated (**Table 1; Table S8**). Among them, 86.73% (37,076) of proteins were functionally annotated. Genes are preferentially distributed in distal chromosomal regions, which are characterized by lower transposable element (TE) density (**Fig. 1D**).

**Table 1.**
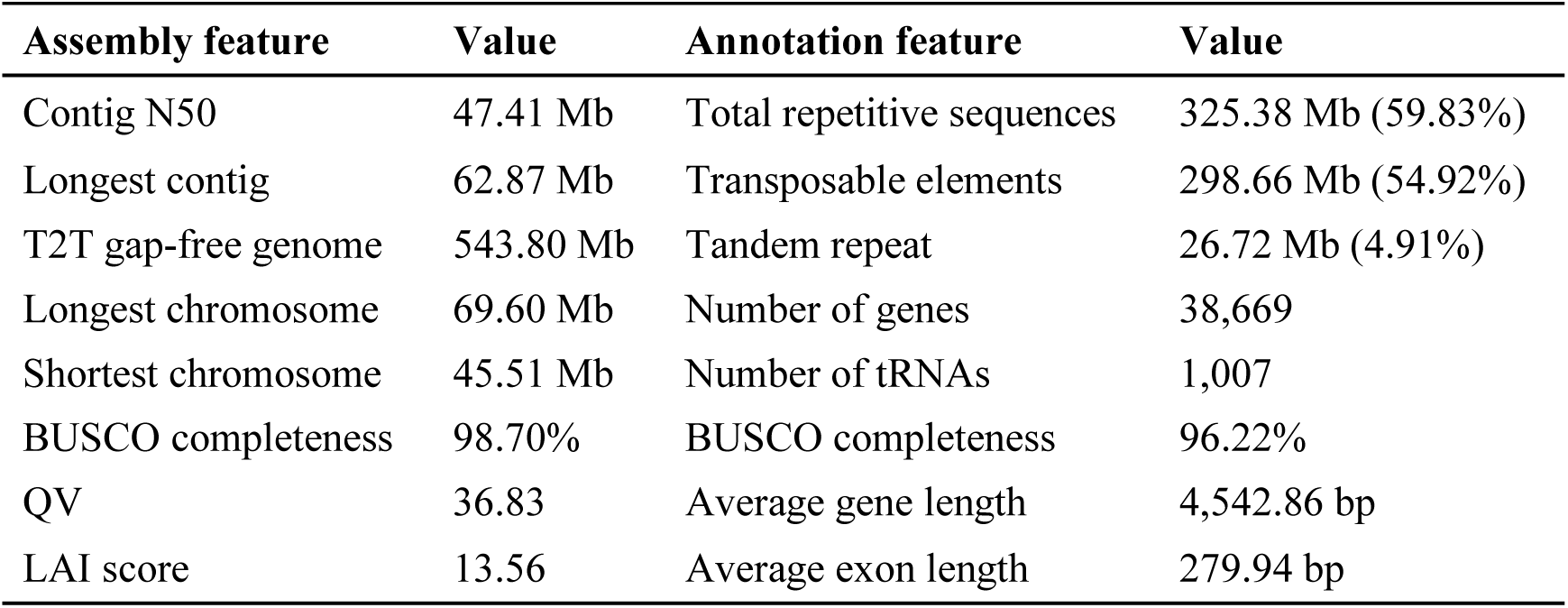
Statistics of genomic assembly and annotation feature of *M*. *exotica*.

| Assembly feature | Value | Annotation feature | Value |
| --- | --- | --- | --- |
| Contig N50 | 47.41 Mb | Total repetitive sequences | 325.38 Mb (59.83%) |
| Longest contig | 62.87 Mb | Transposable elements | 298.66 Mb (54.92%) |
| T2T gap-free genome | 543.80 Mb | Tandem repeat | 26.72 Mb (4.91%) |
| Longest chromosome | 69.60 Mb | Number of genes | 38,669 |
| Shortest chromosome | 45.51 Mb | Number of tRNAs | 1,007 |
| BUSCO completeness | 98.70% | BUSCO completeness | 96.22% |
| QV | 36.83 | Average gene length | 4,542.86 bp |
| LAI score | 13.56 | Average exon length | 279.94 bp |

**Figure 1.**
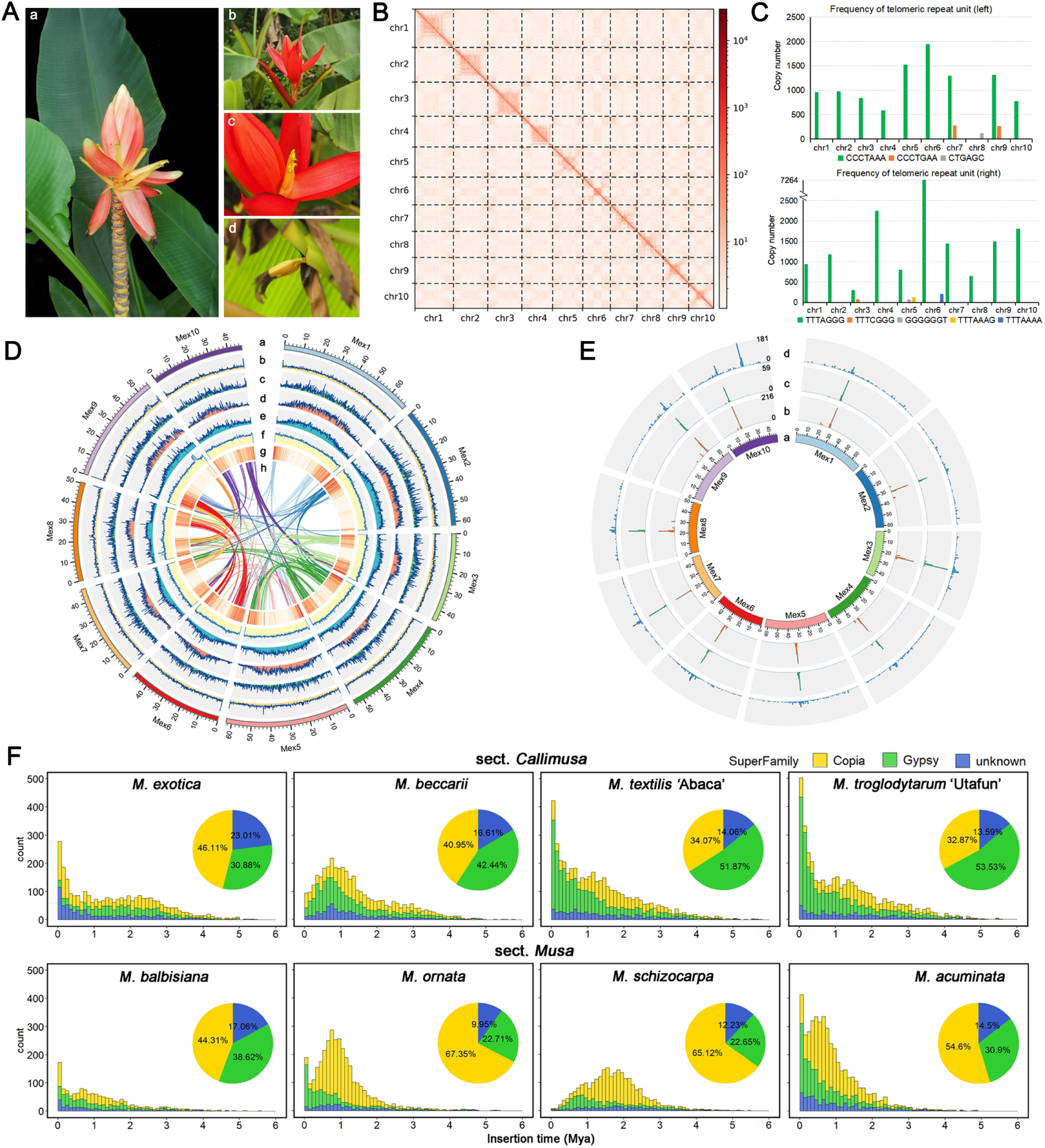
Morphological characteristics and genomic features of *M*. *exotica*. **(A)** Morphology of *M*. *exotica*. a-c: inflorescences; d: fruit. **(B)** Hi-C contact map showing interactions among the 10 pseudo-chromosomes. **(C)** Frequency of telomeric repeat units in the genome. **(D)** Genomic features of the T2T genome assembly. Circles from outside to inside represent: 10 pseudo-chromosomes, tandem repeat distribution, *Gypsy* element distribution, *Copia* element distribution, overall TE distribution, GC content, gene distribution, and syntenic blocks. Window size: 100 kb. **(E)** Centromere organization. Circles from outside to inside represent: 10 pseudo-chromosomes, *Nanica* distribution, *Mex-Cen186* distribution, and *Gypsy*/CRM distribution. Window size: 500 kb. (F) Insertion time distribution of intact LTRs.

### Repetitive sequences shape centromere organization in *M. exotica*

Approximately 59.83% of the *M. exotica* genome consists of repetitive sequences, predominantly long terminal repeat retrotransposons (LTRs) (**Table 1; Table S9**). Insertion-time analyses of 2,960 intact LTRs revealed a recent amplification burst beginning around 0.3 million years ago (Mya), suggesting ongoing transpositional activity. Different *Musa* species exhibit varying preferences for LTR superfamilies, indicating lineage-specific TE dynamics (**Fig. 1F**). Telomeric regions are dominant by the canonical plant telomeric repeat motif C_3_TA_3_/T_3_AG_3_) (**Fig. 1C**).

Centromeric regions of *M*. *exotica* are characterized primarily by enrichment of *Nanica* elements, a long interspersed nuclear element (LINE) proposed as a centromeric indicator in *M*. *acuminata* based on cytogenetic evidence showing strong enrichment in centromeric regions despite the absence of typical high-copy tandem repeats [19]. In total, 2,985 copies are concentrated near the middle of chromosomes and largely overlapping with a 186-bp tandem repeat unit (*Mex-Cen186*) (**Fig. 1E**). In contrast, previously reported centromeric repeats from *M. acuminata* (*Mac-Cen183*) [26], and *Ensete glaucum* (*Egcen*)[32] are largely absent, highlighting divergence in centromere composition across Musaceae species (**Figs. S1–S4**). *Nanica*-enriched regions also consistently coincide with *Gypsy*-enriched but *Copia*-depleted regions (**Fig. S5 and Tables S10–S11**), suggesting a distinct TE landscape associated with putative centromeres. Together these findings indicate that repetitive sequences have played important roles in shaping chromosome architecture and centromere evolution in Musaceae.

### Comparative genomics reveals extensive lineage-specific divergence in Musaceae

Phylogenetic relationships were reconstructed based on 472 single-copy orthologs from Musaceae and outgroup monocot species (**Fig. 2A; Table S12**). Both concatenation-based (IQ-tree) and coalescent-based (ASTRAL) approaches produced highly congruent topologies, with 100% support across all nodes (**Fig. S6**). The three genera within Musaceae (*Musa*, *Ensete*, and *Musella*) were clearly resolved, with *Ensete* and *Musella* forming a well-supported clade (hereafter referred to as Clade EM). Within *Musa*, the two major sections, sect. *Musa* and sect. *Callimusa*, were distinctly separated (**Fig. 2A**). Divergence time estimation indicated that the crown age of Musaceae is approximately 51.89 million years ago (Mya; 95% HPD : 44.05–61.17 Mya) (**Fig. 2A; Fig. S7**). Within *Musa,* sect. *Musa* and sect. *Callimusa* diverged around 39.56 Mya. *M. exotica* diverged from the most recent common ancestor (MRCA) of sect. *Callimusa* approximately 23.77 Mya and is supported as a basal lineage within this section, based on plastome and nuclear ribosomal DNA (nrDNA) evidence (**Figs. S6, S8 and Table S13**). These results provide a robust phylogenetic framework for interpreting genome evolution and trait diversification within Musaceae.

**Figure 2.**
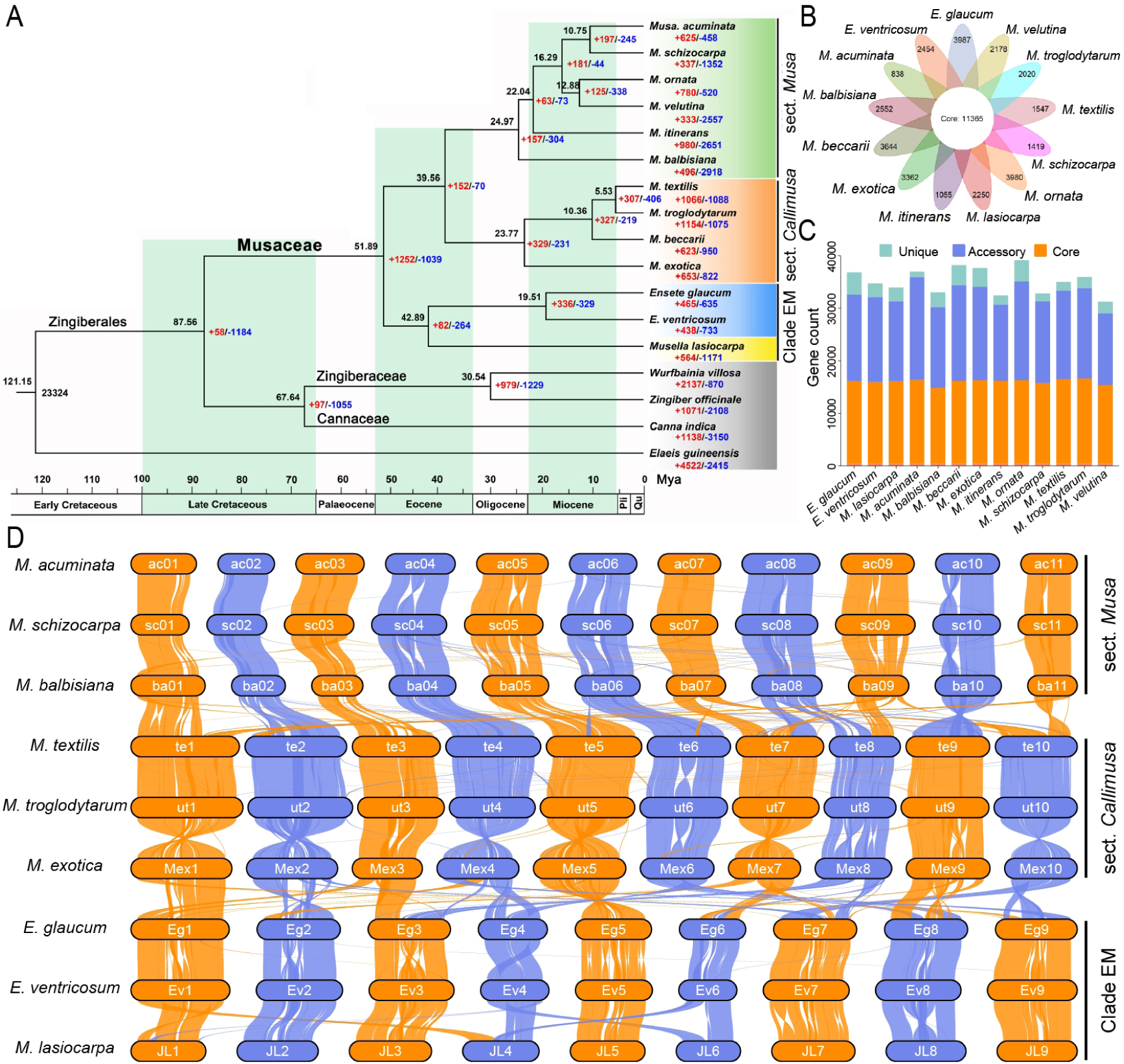
Phylogeny, gene family evolution, and genome synteny across Musaceae species. **(A)** Phylogenetic relationships and divergence time estimation based on 472 single-copy orthologs. Black numbers near each node indicate divergence times, while red and blue numbers represent the numbers of expanded and contracted gene families, respectively. **(B)** Petal diagram of gene families. The central circle shows the number of core orthogroups shared by all species, whereas the petals indicate unique (species-specific) orthogroups. **(C)** Distributions of gene counts across different orthogroup categories. **(D)** Genome-wide synteny among Musaceae species.

To further investigate gene content variation across Musaceae, we conducted a pangenome analysis and identified 70,194 gene families. Of these, 16.19% (11,365) were core (shared by all species), 44.57% (31,286) were unique (species-specific), and 39.24% (27,543) were accessory (**Fig. 2B–C; Tables S14–S15**). The number of unique gene families varied from 838 in *M. acuminata* to 3,987 in *E. glaucum*, with *M. exotica* possessing 3,362 unique families. Functional enrichment analysis showed that species-specific gene families were associated with a broad range of Gene Ontology (GO) terms, spanning 1,076 to 3,116 categories across species, indicating substantial functional diversity (**Table S16 and Fig. S9**). Notably, the significantly enriched GO categories exhibited strong species-specific patterns, with ∼70.51%–100.00% of enriched terms being unique to individual species, collectively indicating pronounced functional divergence among lineages (**Table S16 and Fig. S10**). In contrast, *M. balbisiana* deviated from this trend, showing a higher proportion of enriched terms shared with other species.

This lineage-specific functional divergence was particularly evident in pathways related to pigment metabolism, photosynthesis, and environmental adaptation (**Table S17**). Genes associated with pigment biosynthesis and modification showed clear lineage-specific enrichment. For example, *M. textilis* was enriched in apocarotenoid and abscisic acid biosynthetic processes, whereas *M. itinerans* exhibited enrichment in carotenoid and xanthophyll metabolism, and *M. beccarii* showed strong enrichment in anthocyanin accumulation and UV-induced pigmentation processes, highlighting divergent evolutionary trajectories of pigment metabolism across species. In addition, photosynthesis- and light-related processes were unevenly distributed among species. For instance, *M. itinerans* displayed enrichment in chlorophyll metabolic processes, while *M. lasiocarpa* was enriched in responses to light and far-red light, indicating potential differences in light perception and photosynthetic adaptation.

Furthermore, lineage-specific divergence was also reflected in adaptive and stress-related pathways (**Table S17**). *M. acuminata* exhibited strong enrichment in terpenoid and sesquiterpenoid biosynthesis as well as responses to herbivory and wounding, suggesting a role in chemical defense. In contrast, *M. exotica* showed enrichment in stress-activated protein kinase signaling and cytoskeleton organization, while *E. glaucum* was characterized by nitrogen utilization and meristem development processes, indicating adaptation to nutrient availability and developmental regulation. Meanwhile, *M. beccarii* and *M. itinerans* displayed enrichment in transposition and DNA repair-related processes, implying genome plasticity-associated adaptive mechanisms. Taken together, these results indicate pronounced lineage-specific divergence in gene content across Musaceae, reflecting functional differentiation that may underlie their ecological adaptation and morphological diversification.

To further investigate the structural basis underlying gene content variation across Musaceae, we performed synteny and collinearity analyses. Intraspecific collinearity analysis of *M. exotica* revealed extensive duplicated chromosomal segments, with each chromosome exhibiting syntenic relationships with multiple others (**Fig. 1D**). Synonymous substitution rate (Ks)-based analyses supported the presence of three ancient whole-genome duplication (WGD) events (α, β, and γ) shared across Musaceae species, consistent with previous studies [19, 20], with no evidence for recent species- or lineage-specific WGDs (**Fig. S11**).

Despite this shared WGD history, the three major clades of Musaceae differ in chromosome base numbers, with sect. *Musa* predominantly exhibiting *n* = 11, sect. *Callimusa n* = 10, and Clade EM mainly *n* = 9 (**Fig. 2D**). Synteny analysis revealed largely conserved one-to-one chromosomal relationships within each clade, whereas extensive inter-chromosome rearrangements were observed among clades (**Fig. 2D; Figs. S12–S14**). In contrast, only limited rearrangements were detected within Clade EM, such as those between *E. glaucum* and *M. lasiocarpa* (**Fig. S13**). These results indicate that, although Musaceae species share common ancient WGD events, subsequent genome evolution has been shaped by extensive chromosomal rearrangements, which likely contribute to lineage-specific variation in chromosome number within the species analyzed here and provide a foundation for reconstructing the ancestral karyotype.

### Ancestral karyotype reconstruction reveals extensive chromosome rearrangements in Musaceae

Nine high-quality genomes from representative Musaceae species were used to investigate karyotype evolution in this family (**Fig. 3 and Table S12**). We first identified conserved one-to-one chromosomal relationships within each lineage, defining protochromosomes for ancestral karyotypes (**Fig. 3**). This resulted in 10 protochromosomes for sect. *Callimusa*, 11 for sect. *Musa*, and 9 for Clade EM (*Ensete* and *Musella*) (**Figs. S12–S14**). Notably, a reciprocal chromosome translocation (RCT) was identified between chromosomes 1 and 3 of *M*. *balbisiana* and their counterparts in *M. acuminata* (**Fig. S12A**). By comparing *M. acuminata* and *M. exotica*, we inferred that this translocation occurred independently in *M. balbisiana* (**Fig. S12**), in line with previous reports [33, 34].

**Figure 3.**
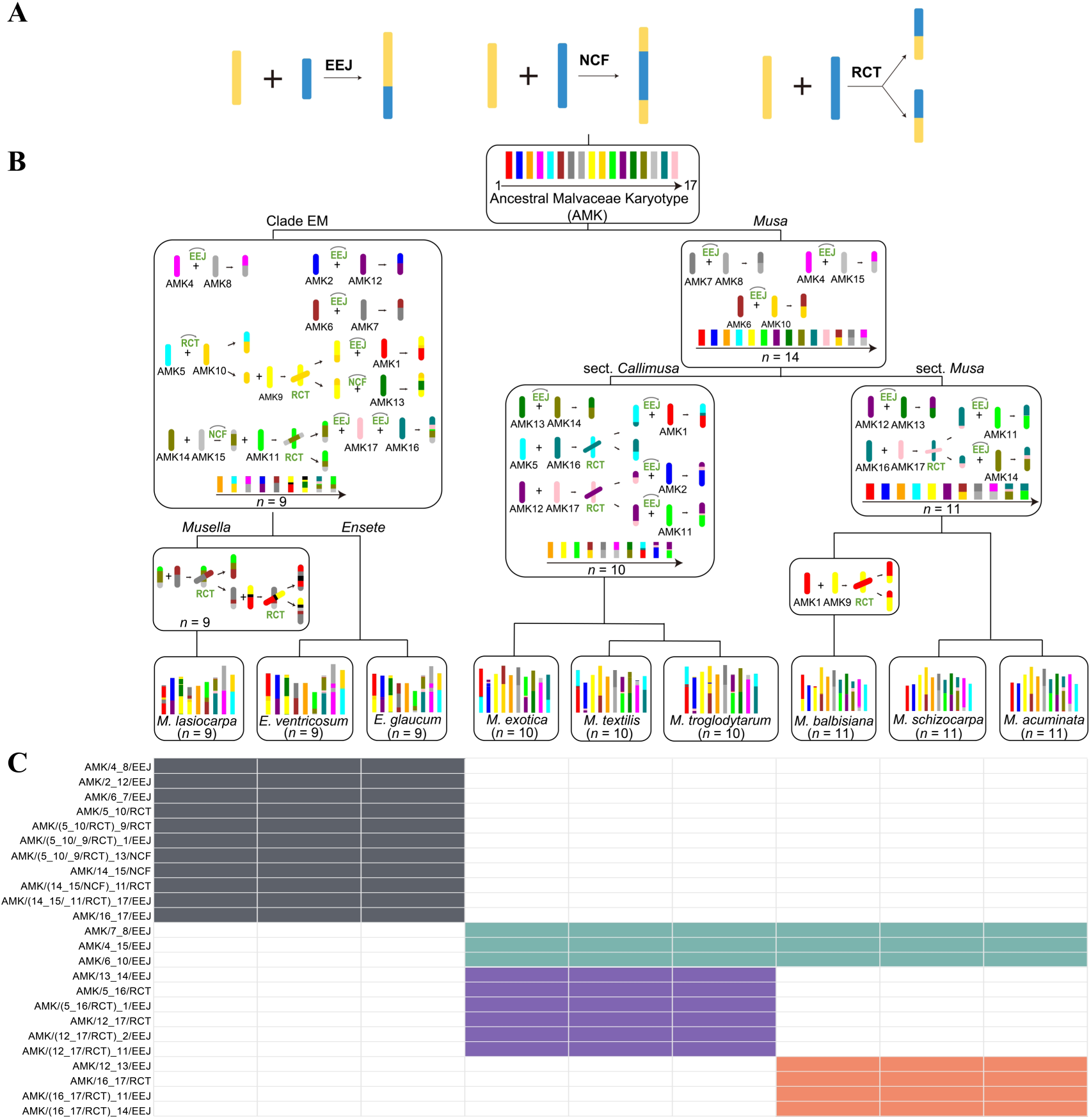
Reconstruction of the Ancestral Musaceae karyotype (AMK). **(A)** Schematic illustration of the three major types of inter-chromosome rearrangements: reciprocal chromosome translocation (RCT), end-to-end joining (EEJ), and nested chromosome fusion (NCF). **(B)** Evolutionary trajectory of karyotype changes across Musaceae lineages. **(C)** Summary of shared inter-chromosome rearrangements among different lineages.

We then compared the protochromosomes among the inferred ancestral karyotypes of sect. *Musa,* sect. *Callimusa* and Clade EM. While many protochromosomes were retained as conserved linkage chromosome-scale syntenic blocks across lineages, others underwent rearrangements consistent with three major mechanisms: reciprocal chromosome translocation (RCT), end-to-end joining (EEJ), and nested chromosome fusion (NCF) (**Fig. 3A**). By further integrating breakpoint comparisons with the outgroup, we inferred the directionality of these rearrangements and reconstructed an ancestral Musaceae karyotype (AMK) comprising 17 protochromosomes (**Fig. 3B; Fig. S15, Supplementary Data and Table S18**).

Based on these identified rearrangements, we reconstructed the evolutionary trajectory of chromosome number changes within Musaceae. Starting from AMK, the common ancestor of *Musa* (sect. *Musa* and sect. *Callimusa*) underwent three EEJs, reducing the chromosome number to 14 (**Fig. 3B**). Subsequent lineage-specific rearrangements led to divergence into sect. *Musa* and sect. *Callimusa*. The ancestral karyotype of sect. *Musa* experienced three additional EEJs and one RCT, resulting in a chromosome number of 11, whereas that of sect. *Callimusa* underwent four EEJs and two RCTs, leading to a *n* = 10 (**Fig. 3B**). Despite this small difference in chromosome number, their chromosomal gene compositions have diverged substantially. Similarly, the lineage leading to Clade EM (*Musella* and *Ensete*) underwent six EEJs, two NCFs, and three RCTs, reducing the chromosome number to *n* = 9. Following this, *Musella* likely experienced two additional RCTs, contributing to its divergence from *Ensete*. Breakpoint analyses further confirmed that these rearrangements are consistently inherited within each lineage, supporting their evolutionary stability (**Fig. 3C; Table S18**).

To explore the functional consequences of inter-chromosome rearrangements, we examined 2,011**–**3,418 genes located within and adjacent to breakpoint regions (±50 genes flanking each breakpoint) across Musaceae genomes (**Tables S18–S19**). Functional annotation and enrichment analyses revealed that these regions harbor extensive functional diversity, with 3,053–3,533 enriched GO terms and 104–115 enriched KEGG pathways (**Tables S19–S20**). Notably, significantly enriched functions exhibited strong lineage-specific divergence among sect. *Musa*, sect. *Callimusa*, and Clade EM (**Table S19 and Fig. S16**). In Clade EM, almost no significantly enriched GO terms were detected, with only a single term identified in *E. glaucum*. In sect. *Musa*, a limited number of significantly enriched GO terms (35) were detected exclusively in *M. acuminata*. In contrast, sect. *Callimusa* exhibited the highest number of significantly enriched GO terms (133–258), with the largest numbers observed in *M. exotica* (258) and *M. textilis* (214).

It is worth noting that Musaceae includes numerous ornamental species characterized by diverse and often vivid bract coloration, which represents a key horticultural trait [21, 22]. Consistent with this, pigmentation- and flavonoid biosynthesis-related pathways were significantly enriched in structurally dynamic regions (**Table S20**). Specifically, in *M. exotica* and *M. textilis*, several genes involved in pigmentation and flavonoid biosynthesis—including key structural enzymes such as *CHS* (chalcone synthase), and *F3H* (flavanone 3-hydroxylase)—were significantly enriched in these regions (**Tables S20–S21**), providing a rationale for further investigation into the genetic basis of bract color variation. *CHS* catalyzes the first committed step of flavonoid biosynthesis, while *F3H* functions downstream to generate dihydroflavonols, key intermediates for anthocyanin and flavonol production (**Fig. 4A**).

**Figure 4.**
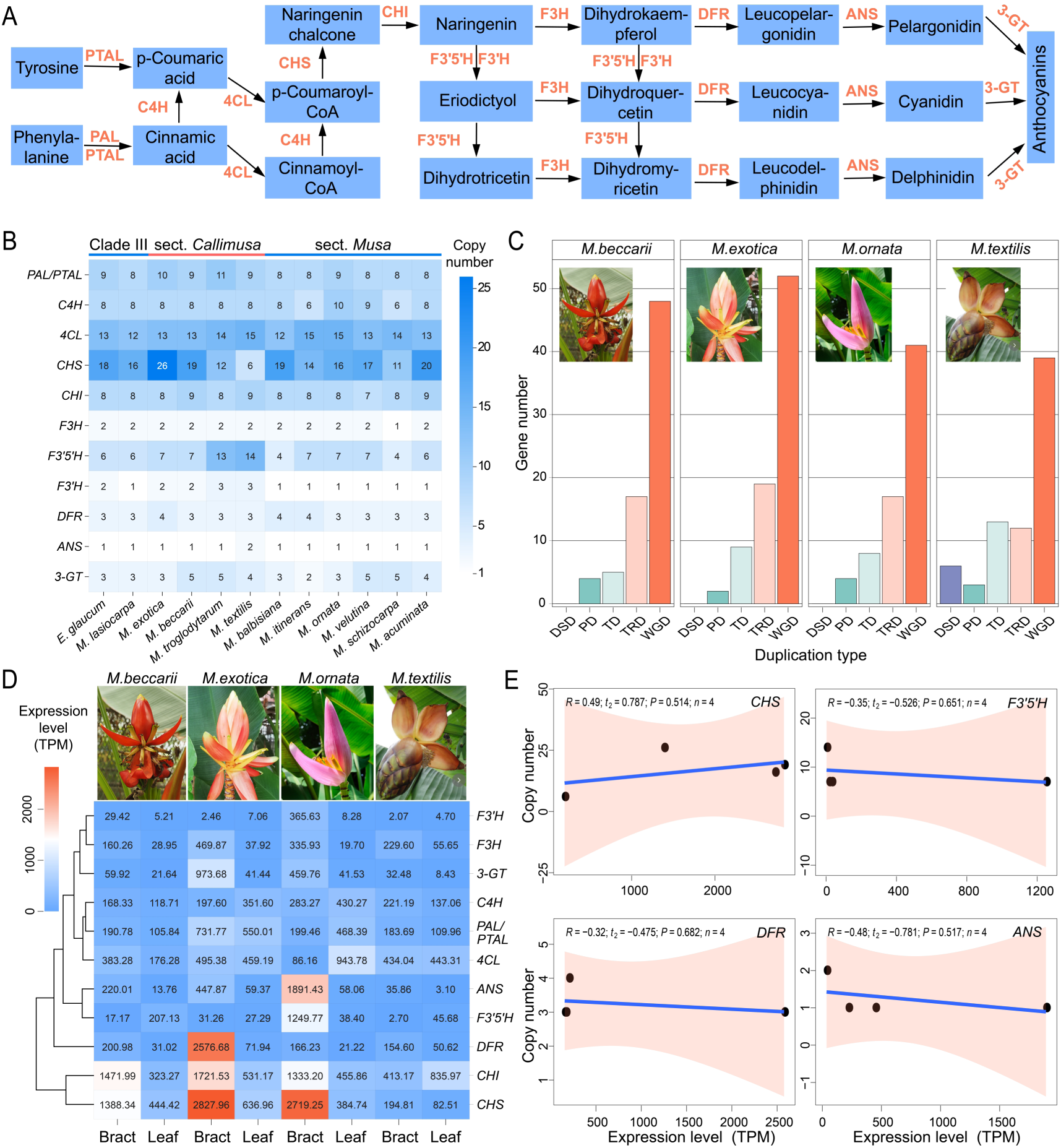
Integrated analyses of anthocyanin biosynthesis in Musaceae. **(A)** Schematic representation of the anthocyanin biosynthesis pathway. Enzymes are indicated above the arrows. *PAL*, phenylalanine ammonia-lyase; *PTAL*, phenylalanine/tyrosine ammonia-lyase; *C4H*, cinnamic 4-hydroxylase; *4CL*, 4-coumarate CoA ligase; *CHS*, chalcone synthase; *CHI*, chalcone isomerase; *F3H*, flavanone 3-hydroxylase; *F3’H*, flavonoid 3’-hydroxylase; *F3’5’H*, flavonoid 3’,5’-hydroxylase; *DFR*, dihydroflavonol 4-reductase; *ANS*, anthocyanidin synthase; *3*-*GT*, anthocyanidin 3-O-glucosyltransferase. **(B)** Copy number variation of structural genes involved in the anthocyanin biosynthesis pathway across Musaceae species. **(C)** Distribution of duplication modes for anthocyanin biosynthesis genes. Gene duplication types include dispersed duplication (DSD), proximal duplication (PD), tandem duplication (TD), transposed duplication (TRD), and whole-genome duplication (WGD). **(D)** Total expression levels of genes encoding different enzymes in bracts and leaves. Expression levels were normalized using TPM. **(E)** Correlation between gene copy number and expression levels of key anthocyanin biosynthesis genes. Scatter plots show the relationships for *CHS*, *F3′5′H*, *DFR*, and *ANS*. Linear regression lines were fitted using an ordinary least squares (OLS) model, with 95% confidence intervals indicated by shaded regions. Pearson’s correlation coefficients (*R*) were used to assess statistical significance. The *R*, *t*, *P* values, and sample sizes (*n*) are provided in each panel, indicating no significant correlation between gene copy number and expression level.

This non-random distribution is consistent with the possibility that chromosomal rearrangements have influenced the genomic organization of pigmentation-related genes through repositioning, clustering, and/or preferential retention. Such structural dynamics may, in turn, have contributed to the rewiring of metabolic pathways and the diversification of bract coloration during Musaceae evolution.

### Anthocyanin biosynthesis diversification is associated with transcriptional regulation rather than gene dosage

The enrichment of anthocyanin biosynthesis genes within structurally dynamic genomic regions, together with the pronounced diversity of bract coloration across Musaceae species (**Fig. 4**), suggests a potential link between genome structural variation and pigmentation traits. To further evaluate this relationship, we systematically investigated the copy number variation, duplication modes, and expression patterns of anthocyanin biosynthesis genes.

Across all examined species, a complete set of anthocyanin biosynthesis genes is retained, indicating overall conservation of the pathway (**Fig. 4A–B**). However, a clear positional trend in gene copy number is observed along the pathway: upstream genes (e.g., *PAL*/*PTAL*, *C4H*, and *4CL*) generally exhibit higher copy numbers, whereas downstream genes (e.g., *DFR*, *ANS*, and *3-GT*) are present in fewer copies (**Fig. 4B**). This pattern suggests stronger dosage requirements or functional constraints at early steps. Notably, substantial copy-number variation is detected in several genes, particularly for *4CL*, *CHS,* and *F3′5′H.* For instance, *CHS* copy number varies markedly across species, ranging from six copies (three within rearrangement breakpoint regions) in *M*. *textilis* to 26 copies (eight within breakpoint regions) in *M. exotica*, while *F3′5′H* shows moderate but lineage-specific expansion, reaching up to 14 copies in *M*. *textilis*. Phylogenetic analysis further reveals that *CHS* has undergone multiple species-specific duplication events in *M. exotica*, whereas *F3′5′H* exhibits similar lineage-specific duplications in *M. textilis* (**Fig. S17**). In contrast, genes such as *F3H*, *DFR*, and *ANS* are highly conserved in copy number and retain phylogenetic relationships consistent with the species tree (**Fig. 4B; Fig. S17**).

Focusing on four species with contrasting bract coloration—*M. exotica*, *M*. *beccarii*, and *M*. *ornata* (brightly colored) versus *M*. *textilis* (duller)—gene duplication analysis revealed that whole-genome duplication (WGD) is the predominant source of anthocyanin biosynthesis genes, with additional contributions from transposed duplication (TRD), and minor contributions from tandem (TD) and proximal duplication (PD), whereas dispersed duplication (DSD) is nearly absent (**Fig. 4C; Table S22**). In contrast, the genomic background shows a broader distribution of duplication modes, including substantial DSD (**Table S22**). These patterns suggest that anthocyanin pathway genes are likely subject to stronger selective constraints, favoring the retention of dosage-balanced WGD-derived duplicates while limiting random dispersal.

At the individual gene level, several enzymes—including *PAL*/*PTAL*, *C4H*, *4CL, CHI*, *DFR*, and *3-GT*—are predominantly WGD-derived across all four species, whereas *F3H* is exclusively TRD-derived (**Table S23**). *CHS* exhibits multiple duplication modes (WGD, TRD, TD, and PD) in the three pigmented species, but is represented by single DSD-derived copies in the duller species *M. textilis*. *F3′5′H* shows lineage-specific expansion: in pigmented species, it is mainly TRD-derived with seven conserved copies, whereas in *M. textilis* it is primarily TD-derived with the highest copy number of 14. *ANS* is represented by a single TRD-derived copy in pigmented species but is WGD-derived in *M. textilis*. Together, these results highlight a conserved pathway architecture accompanied by differential evolutionary flexibility among specific gene families.

Expression profiling revealed a strong association between transcriptional activation and bract pigmentation. Most structural genes show markedly higher expression in bracts than in leaves across all four species, indicating coordinated activation of the anthocyanin pathway (**Fig. 4D–E; Fig. S18 and Tables S24–S28**). In the three pigmented species, several genes—including *CHS*, *CHI*, *F3′5′H*, *DFR*, *ANS*, and *3-GT*—exhibit higher expression levels compared with *M. textilis*, with *CHS*, *CHI*, *F3’5’H*, and *ANS* showing more than twofold upregulation (**Fig. 4D; Table S28**). Among these, *CHS* displays exceptionally high expression in pigmented species, consistent with its expanded copy number and central role in pathway flux control. In contrast, downstream genes such as *DFR* and *ANS* show elevated expression despite relatively low copy numbers, suggesting that transcriptional regulation rather than gene dosage drives their functional contribution (**Fig. 4E; Fig. S19**). Hydroxylase genes exhibited species-specific expression patterns. Notably, *F3′5′H* exhibits high expression in *M. ornata* but relatively low expression in the duller species *M. textilis*, despite its expanded copy number in the latter, indicating a decoupling between gene dosage and transcriptional output (**Fig. 4E**). Together, these findings highlight that variation in anthocyanin biosynthesis is shaped not only by gene copy number but also by transcriptional regulation.

To further investigate the regulatory basis underlying these expression differences, we examined the potential involvement of transcription factors (TFs). Orthogroup inference was performed using MYB and bHLH protein sequences identified by iTAK v2.0.5 [35] across four species (**Tables S29-S36**), enabling cross-species comparison of TF expression patterns within a unified orthology framework. Based on integrated expression data from eight samples (four species × two tissues), we performed Pearson correlation analysis between TFs and anthocyanin biosynthetic genes. A total of 37 and 34 high-confidence MYB–enzyme and bHLH–enzyme associations were identified, respectively (|*r*| ≥ 0.8, *P* < 0.01; **Fig. 5; Tables S37–S40**). MYB transcription factors exhibited extensive correlations—both positive and negative—with all major structural genes, indicating broader co-expression relationships across the pathway (**Fig. 5A**). In contrast, bHLH transcription factors were strongly correlated with most structural genes but showed limited association with *F3′5′H*, suggesting potential divergence in regulatory connectivity for this branch of the pathway (**Fig. 5B**). Clustering of the correlation matrix further revealed distinct co-expression modules, reflecting coordinated and gene-specific expression relationships between TFs and anthocyanin biosynthetic genes across species. A co-expression network derived from this correlation matrix further illustrates these interactions, highlighting potential regulatory links and modular organization between MYB/bHLH transcription factors and structural genes (**Fig. 5C**). To further link these regulatory patterns to genome structural variation, we found that several co-expression modules include transcription factors located within chromosomal rearrangement breakpoint regions. Specifically, one MYB-associated module and five bHLH-associated modules that show significant correlations with enzyme genes contain TFs residing in breakpoint regions, suggesting that structural rearrangements may influence the organization of transcriptional regulatory networks underlying anthocyanin biosynthesis (**Fig. 5**). Together, these results identify candidate MYB and bHLH regulators associated with anthocyanin biosynthetic gene expression.

**Figure 5.**
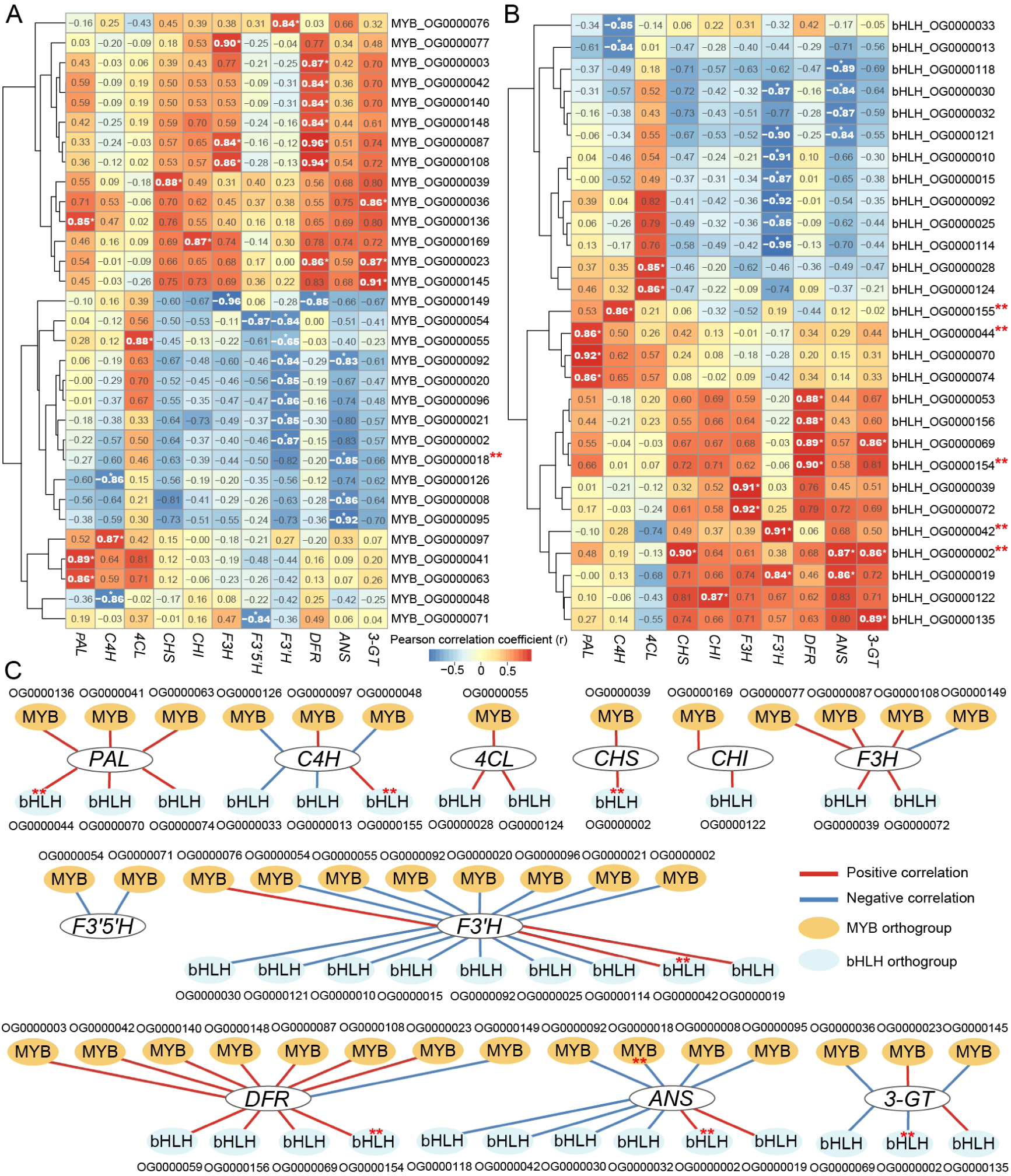
Co-expression relationships between transcription factors and anthocyanin biosynthetic genes across Musaceae species. **(A)** Heatmap showing significant Pearson correlations between MYB transcription factors and anthocyanin biosynthetic structural genes. **(B)** Heatmap showing significant Pearson correlations between bHLH transcription factors and anthocyanin biosynthetic structural genes. Correlation coefficients (r) were calculated based on gene expression levels across eight samples (bract and leaf tissues from four species). Only significant correlations are shown (|*r*| ≥ 0.8 and *P* < 0.01). Positive correlations are indicated in red and negative correlations in blue. Hierarchical clustering was applied to transcription factors (rows), while structural genes (columns) are ordered according to their positions in the anthocyanin biosynthesis pathway. Asterisks indicate statistically significant correlations. **(C)** Co-expression network illustrating the interactions between transcription factors (MYB and bHLH) and anthocyanin biosynthetic genes. Nodes represent genes, and edges represent significant correlations. Red edges indicate positive correlations, whereas blue edges indicate negative correlations. Notably, double red asterisks across all panels denote transcription factor orthogroups that contain genes located within chromosomal breakpoint regions.

## Discussion

### Chromosomal evolution and karyotype diversification in Musaceae

The availability of high-quality, chromosome-level genomes has enabled a refined reconstruction of the Musaceae evolutionary history, particularly regarding karyotype evolution and its functional consequences. In contrast to earlier studies based on fragmented assemblies [36], we reconstructed an ancestral Musaceae karyotype (AMK) with *n* = 17 protochromosomes, providing a robust framework for tracing chromosome evolution across the family. Despite shared ancient whole-genome duplication (WGD) events (α, β, and γ), analyzed Musaceae lineages exhibit considerable variation in chromosome numbers (*n* = 11 in sect. *Musa*, *n* = 10 in sect. *Callimusa*, and *n* = 9 in Clade EM). Synteny and breakpoint analyses demonstrate that this divergence is primarily driven not by recent WGD events, but by extensive inter-chromosomal rearrangements, including reciprocal chromosome translocation (RCT), end-to-end joining (EEJ), and nested chromosome fusion (NCF). These rearrangements display strong lineage-specific patterns and are stably inherited within clades, providing clear phylogenetic signals. For instance, the shared rearrangement patterns between *Musella* and *Ensete* in Clade EM, contrasted with their divergence from *Musa*, reinforce their close evolutionary relationship, while distinct rearrangement trajectories differentiate sect. *Musa* and sect. *Callimusa*.

Importantly, chromosome number alone is insufficient to capture evolutionary relationships, as species with identical chromosome numbers may possess markedly different chromosomal architectures. By reconstructing ancestral karyotypes and tracing breakpoint-defined rearrangements, we reveal the dynamic processes shaping genome evolution and offer a more nuanced framework for interpreting species divergence. Crucially, chromosomal rearrangements are not merely structural events—they carry functional consequences. Genes located within and adjacent to rearrangement breakpoint regions exhibit broad functional diversity and strong lineage-specific enrichment patterns. Notably, in sect. *Callimusa*, breakpoint-associated regions are significantly enriched for genes involved in pigmentation and flavonoid biosynthesis, including key structural enzymes such as *CHS* and *F3H*, as well as regulatory components like MYB and bHLH transcription factors. Integrative analysis combining co-expression networks and genome structural variation reveals that several TF modules significantly associated with anthocyanin biosynthetic genes are located within these breakpoint regions. This non-random co-localization of structural and regulatory elements suggests that chromosomal rearrangements may have facilitated coordinated repositioning, clustering, and/or preferential retention of functionally related genes, contributing to the reorganization of transcriptional and metabolic networks.

Notably, this structural organization likely provides the genomic foundation for phenotypic diversification. The clustering of pigment-related structural and regulatory genes within breakpoint regions implies that chromosomal rearrangements may have facilitated coordinated transcriptional regulation and contributed to the rewiring of metabolic networks. Collectively, the evolutionary history of Musaceae appears to reflect a combination of shared ancient polyploidy and lineage-specific chromosomal rearrangements, with the latter likely shaping patterns of genomic organization and contributing to phenotypic diversity.

### Regulatory and evolutionary mechanisms underlying anthocyanin biosynthesis in Musaceae

Building upon observed genomic and structural variations in Musaceae, the integration of gene duplication and expression analyses reveals a complex regulatory architecture underlying anthocyanin biosynthesis in this family. Although substantial variation in gene copy number exists among species, particularly for key genes such as *CHS* and *F3′5′H*, these differences do not consistently translate into proportional changes in transcript abundance. This decoupling of gene dosage and transcriptional output highlights that copy number alone is insufficient to explain phenotypic variation in pigmentation, emphasizing an important role of transcriptional regulation in modulating pathway activity.

Consistent with this observation, co-expression network analysis illustrates extensive and coordinated associations between transcription factors (TFs) and structural genes. MYB transcription factors show broad connectivity across the entire pathway, with *F3′H* and *DFR* serving as major hubs, each forming up to eight TF–enzyme associations, supporting their pivotal regulatory roles. This aligns with prior reports that MYB repressors (e.g., MaMYB4) can inhibit multiple structural genes (e.g., *F3′H*, *ANS*, *3-GT*), reducing anthocyanin accumulation in *Musa* tissues [37]. In contrast, bHLH transcription factors exhibit more selective correlations, particularly showing limited association with *F3′5′H* but strong connectivity with *F3′H*, suggesting branch-specific regulatory divergence within hydroxylation pathways.

Functionally, these network patterns are reinforced by expression profiles: brightly colored bracts show upregulation of key structural genes (e.g., *CHS*, *CHI*, *F3’5’H*, and *ANS*), whereas reduced expression in *M. textilis* may underlie its comparatively dull pigmentation. In *M. ornata*, which produces pink bracts, *F3′H*, *F3′5′H*, and *ANS* reach peak expression. As key branch-point enzymes controlling flux partitioning within the pathway, *F3′H* and *F3′5′H* direct metabolic flux toward different anthocyanin derivatives, while *ANS* catalyzes the conversion of leucoanthocyanidins to anthocyanidins, representing a key downstream step in pigment formation. These coordinated expression patterns parallel observations in other species, such as flux shifts caused by *F3′H* in *Euphorbia pulcherrima* [38], and elevated *F3′5′H* expression driving pink coloration in *Curcuma alismatifolia* [39].

Gene duplication patterns provide an evolutionary framework for the observed decoupling between gene dosage and transcriptional output. Preferential retention of WGD-derived duplicates, combined with limited contribution from dispersed duplication (DSD), suggests selection for dosage balance in tightly coordinated pathways. Lineage-specific expansions, such as increased *F3′5′H* copies in *M. textilis* without concomitant transcriptional activation, point to post-duplication regulatory divergence or functional attenuation. Structural domain analyses further indicate that some duplicates have undergone divergence, potentially enabling functional differentiation and regulatory innovation.

Taken together, these results demonstrate that the evolution of anthocyanin biosynthesis in Musaceae is likely shaped by the interplay of chromosomal rearrangements, gene duplication, regulatory divergence, and transcriptional control. This multilayered architecture not only explains the decoupling of gene dosage and expression but also provides a mechanistic basis linking genomic evolution to observable phenotypic variation, ultimately underpinning the remarkable diversity of bract coloration across Musaceae species.

## Materials and methods

### Sample collection and sequencing

The *Musa exotica* individual was obtained and cultivated in the greenhouse of South China Botanical Garden, Chinese Academy of Sciences (SCBG, CAS), Guangzhou, China. Fresh young leaves were collected and immediately frozen in liquid nitrogen. Total genomic DNA was extracted using the CTAB method [40]. The Illumina short paired-end (PE, 2×150 bp) reads were sequenced on a HiSeq 4000 platform after libraries were constructed with an insertion size of 350 bp. For ONT sequencing, large fragments (>20 kb) were selected with BluePippin, libraries were constructed using Ligation Sequencing Kit SQK-LSK109 and then sequenced on the Promethion platform. PacBio sequencing was performed on Sequel II platform under the circular consensus sequencing (CCS) mode to generate HiFi reads. For Hi-C library construction, young leaves were fixed, cut with the restriction enzyme DpnII, marked with biotin, and purified. Hi-C libraries were sequenced using the HiSeq 4000 platform. For RNA sequencing (RNA-seq), we collected leaves, flowers, and bracts. Total RNA was extracted using TRIzol reagent, and mRNAs enriched using oligodT beads were broken into short fragment for cDNA synthesis. Finally, PE sequencing of cDNA libraries was conducted using the HiSeq X Ten platform. For quality control, the Illumina, Hi-C, and RNA raw reads were filtered using Fastp v. 0.23.2 [41] with parameters set as “-q 20 -u 40 -n 1”. For ONT raw reads, we used Porechop v. 0.2.4 (https://github.com/rrwick/Porechop) to remove adaptors, and then used Fastp to discard low-quality reads (average quality score < 7) and reads that were too short (length < 1000) with parameters “-u 100 -e 7 -l 1000”. HiFi raw reads were filtered by Fastp with parameters “-q 20 -u 40”. Finally, a total of 174.54 Gb ONT, 17.73 Gb HiFi, 45.17 Gb Illumina, 133.67 Gb Hi-C, and 26.77 Gb RNA clean reads were obtained (**Table S2**).

### Genome survey and assembly

Using Illumina reads, we employed Jellyfish v2.3.0 [42] to calculate *K*-mer with 17-31, the outputs were sent to GenomeScope v1.0.0 [43] to estimate genome size and heterozygosity level. For genome assembling, ultralong-ONT (ul-ONT) reads were screened out using Seqkit v2.3.0 [44]. We used two strategies to assemble genome. In the first strategy, NextDenovo (https://github.com/Nextomics/NextDenovo) was used to correct the longest 60× ul-ONT reads with parameters “seed_depth = 60, genome_size = 510m, correction_options = -b”, and then to assemble corrected reads longer than 80 kb and 85 kb, respectively. The assembly results of these two datasets showed that longer reads produce better continuity of contigs (**Table S3A**). In the second strategy, a combination of 17.73 Gb (c. 34×) HiFi reads, 53.91 Gb (c. 105×) ul-ONT reads, and 133.67 Gb (over 200×) Hi-C reads was used, and hifiasm v0.19.3 [45] with parameter “--ul, --h1, --h2” was applied for genome assembly, resulting in a contig N50 of 47.41 Mb, which showed the best continuity (**Table S3A–B**). The completeness of primary assemblies was assessed based on the viridiplantae_odb10 database using BUSCO v5.4.3 [46] with default parameters. The assembly from hifiasm shows the best completeness compared to the assemblies of Nextdenovo (**Table S7A**). Thus, the heterozygous assembly (named p_ctg) from hifiasm was used to construct the final genome. This assembly was polished using NextPolish v1.4.1 [47] with Illumina reads and default parameters. Purge_Haplotigs v1.1.2 [48] was then applied to remove redundant sequences with default parameters and Illumina reads. Hi-C reads were aligned to the purged assembly using BWA v0.7.17 [49], and Juicer v1.6 [50] with default parameters was employed to identify valid interaction pairs. 3D-DNA v201008 [51] was then used to anchor the contigs into pseudo-chromosomes with parameter “-r 0”, and Juicebox v1.11.08 [50] was applied to manually improve the precision of the anchoring.

Gaps in pseudo-chromosomes were patched using TGS-GapCloser v1.2.1 [52] with the two primary assemblies generated from Nextdenovo and parameter “--min_match 500”. The terminal 100 kb sequence of the chromosome missing telomere was extracted as probe for searching its telomere. The polished heterozygous assembly and the two haplotype assemblies (named hap*.p_ctg) generated by hifiasm, along with the two primary assemblies from Nextdenovo, were combined to create a search database. This database was mapped against each probe using Minimap2 v2.24 [53] with parameters “-ax asm20 -m 20000 --sam-hit-only”, and the sequence most similar to each probe was selected for telomere repair. The telomere-repaired genome was polished again using NextPolish with Illumina reads, yielding the T2T gap-free genome. The integrity of this genome was assessed based on the embryophyta_odb10 database using BUSCO, and was also estimated by aligning HiFi and Illumina reads to the genome using BWA, and SAMtools v1.14 [54] was applied with “flagstat” option to calculate the percentage of mapped reads. The QV was calculated using Merqury [55] with Illumina reads, and the LAI score was calculated using LTR_retriever v2.9.1 [56], with default parameters.

### Identification and annotation of repeat elements

The Extensive De-Novo TE Annotator (EDTA) v2.1.0 [57] was used to identify TEs with parameters “--species others --step all --evaluate 1 --sensitive 1 --anno 1”. The resulting non-redundant TE libraries were sent to RepeatMasker v4.1.0 [58] to soft-mask repeats with parameter “-xsmall”. The insertion time of intact LTRs was also calculated by EDTA. We applied TEsorter [59] to classify TEs using the rexdb-plant database [60]. Tandem Repeats Finder v4.09 [61] was employed to identify TRs. Tidk v0.2.1 (https://github.com/tolkit/telomeric-identifier) was used to analyze telomeric repeats with parameters “--minimum 5 --maximum 12 -e tsv –log --threshold 50 --distance 1000000”. The *Nanica* and *Egcen* were downloaded as probes to locate the centromeres, and Blastn v2.11.0 [62] was used to perform searches with default parameters. Tandem Repeats Finder and followed the pipeline Telomeres_and_Centromeres (https://github.com/Immortal2333/Telomeres_and_Centromeres) were also used to find centromeric TR unit.

### Gene prediction and annotation

Using the soft-masked genome as input, we adopted a strategy combining RNA-seq evidence, *ab initio* prediction, and homology search approaches to predict and annotate genes using Funannotate v1.8.15 (https://github.com/nextgenusfs/funannotate). For the RNA-seq evidence-based method, the command “funannotate train” was used to perform genome-guided RNA-seq assembly, by running the embedded pipeline Hisat2 v2.2.1, Trinity v2.8.5, and PASA v2.5.2, with parameters “--stranded no --max_intronlen 100000”. The script “funannotate predict” was used with proteins of *M*. *beccarii*, *M*. *textilis*, and *M*. *troglodytarum* as homologous evidence, and parameters “busco_db embryophyta --max_intronlen 100000 --genemark_mode ET --busco_seed_species arabidopsis --organism other --repeats2evm”. This script performed a comprehensive *ab initio* gene prediction using AUGUSTUS v3.5.0, GeneMark v4.71, Snap v2006-07-28, GlimmerHMM, EVidence Modeler v1.1.1. The “funannotate update” script was then applied to update those predicted gene models using RNA-seq data. BUSCO was used to assess the completeness of gene annotation based on embryophyta_odb10 database. The functional annotation of protein was firstly conducted by EggNOG-mapper v 2.1.10 [63], with parameters “-m diamond --dbmem --tax_scope Viridiplantae”. We also used the command “funannotate iprscan”, by which InterProScan v5.62-94.0 [64] was employed, to annotate protein domains. Finally, using the results from EggNOG-mapper and InterProScan as input, the “funannotate annotate” script was run to complete the final functional annotation.

### Assembly and annotation of plastome and nrDNA

We retrieved next-generation sequencing (NGS) data from NCBI (https://www.ncbi.nlm.nih.gov/) and used GetOrganelle v1.7.7.0 [65] to assemble plastomes and nrDNAs. The NGS raw reads were filtered using Fastp with default parameters. Plastomes were aligned using MAFFT v7.388 [66] and annotated using the “Annotation Transfer” option in Geneious prime v2019.2.1 [67], with *M. paracoccinea* (GenBank accession No.: OK012343) as the reference. Similarly, nrDNA annotation was performed with *M. textilis* (GenBank accession No.: LC610767) as the reference. In total, we assembled and annotated five plastomes (five species) and 26 nrDNAs (25 species) (**Table S13**).

### Phylogeny analyses and divergence time estimate

We used MAFFT to align 27 plastomes with one inverted-repeat region removed and 27 nrDNAs (26 species), respectively (**Table S11**). The alignments were trimmed using Gblocks v0.91b [68] with parameters “-b4=5 -b5=h -t=d -e=.2”. The maximum likelihood (ML) analyses and Bayesian inference (BI) analyses were both performed for the plastome and nrDNA dataset. For BI analyses, the GTR+I+G4 was selected as the best-fit model for the two datasets using ModelTest-NG [69] under the corrected Akaike Information Criterion. Then, we applied Mrbayes v3.2.6 [70] to conduct two MCMC runs with one million generations, sampling every 100 generations and discarding the first 20% as burnin. Finally, Tracer v1.7.2 [71] was used to ensure that the effective sample size for each parameter was larger than 200. For the ML analyses, we applied IQ-tree v2.2.3 [72] with the automatically selected best-fit substitution model (-m MFP) and the 1000 ultrafast bootstrap replicates (-B 1,000).

The longest protein of each nuclear gene was used to perform the GF analysis. We identified the single-copy GFs by using OrthoFinder v2.5.4 [73], and then applied Muscle v3.8.31 [74] to align proteins for single gene. Gblocks with parameters “-b4=5 -b5=h -t=p -e=.2” was used to trim aligned sequences, which were then concatenated to build a ML tree employing IQ-tree with paramater “-m MFP -B 1,000”. Gene trees were also constructed applying IQ-tree with the same paramater and imported into ASTRAL v5.7.8 [75] to infer coalescent-based species tree. We estimated divergence times for the concatenated dataset applying MCMCTree in PAML package v.4.9j [76], with parameters: clock = 2, model = 0, burnin = 200,000, nsample = 1000,000, and sampfreq = 100. Tracer was used to test the convergence. Four calibration points were set: a 95% HPD of 110-130 Mya was applied to constrain the split of Zingiberales/Arecales according to TimeTree database [77] (http://www.timetree.org/, accessed on March 5, 2024); the oldest-known fossil of Zingiberaceae, *Spirematospermum chandlerae* [78], was used to calibrate the crown age of Zingiberales (83.5 Mya, 95% HPD 75-92 Mya); another Zingiberaceae fossil, *Zingiberopsis attenuate* [79], was applied to constrain the split of Cannaceae/Zingiberaceae (65 Mya, 95% HPD 60-75 Mya); the only known Musaceae fossil, *E. oregonense* [80], was used to calibrate the split of *Ensete*/*Musella* (43 Mya, 95% HPD 41-49 Mya).

### Gene family expansion/contraction and GO enrichment

We used CAFE5 [81] with default parameters to assess the expansion and contraction of GFs. The GF statistics from OrthoFinder and the phylogenetic tree from MCMCTree were used as input for CAFE5. The built-in script clade_and_size_filter.py of CAFE5 was employed to remove GFs with significant copy-number variations across different species. GO enrichment analysis of significantly (p < 0.05) expanded and contracted GFs was performed using the R package clusterProfiler v4.6.2 [82]. OrthoFinder was also used to analyze GFs across 13 Musaceae species (**Fig. 2A–C**), to categorize them into core, accessory, and species-specific GFs.

### Genomic collinearity and WGD analyses

The longest protein of each gene was used to perform the collinearity analysis. Nine representatives were chosen for the analysis of Musaceae (**Fig. 2D and Table S12**). Then, the recently published chromosome-level genome of *M*. *troglodytarum* ‘Karat’ [20] were added for a separate analysis of sect. *Callimusa* (**Fig. S12C–D**). *E*. *ventricosum* [83] was included to explore genomic collinearity in *Musella*-*Ensete* clade (**Fig. S13**). Blastp v2.14.1+ [62] was used to align proteins, with parameters: -evalue 1e-5 -outfmt 6 -num_alignments 5. MCScanX [84] was then applied to find collinear blocks (CLBs) based on the Blastp results. In addition, WGDI v0.6.5 [11] was employed with the “-d” option to generate the collinear dot plots for each species pair.

We used five species to perform the WGD analyses. WGDI with the “-icl” option was applied to identify CLBs and collinear genes (CLGs) within or between species. The “-ks” option was used to calculate *Ks* of CLGs, followed by employing “-bi” option to integrate CLBs and *Ks* information, with parameter “ks_col = ks_NG86”. Then, we used the “-c” option with parameter “tandem = false” to remove CLBs caused by tandem duplicates. For *M*. *exotica*, we utilized the csv file yielded from the “-c” option to count the number of CLGs for each gene, and the “-bk” option was used to plot *Ks* dots of CLGs. The “-kp” option was applied to draw the frequency distribution peaks of *Ks* values, and we fit each *Ks* peak using the “-pf” option to obtain the fitting parameters. Finally, we performed the “-kf” command to plot the fitted frequency distribution peaks.

### Ancestral genome reconstruction

We followed the workflow of WGDI toolkit (https://github.com/SunPengChuan/wgdi-example/blob/main/Karyotype_Evolution.md) to reconstruct ancestral karyotypes of Musaceae. Gene sequence at the rearrangement breakpoints was compared with those in the outgroup to distinguish protochromosomes and identify the directionality of chromosome rearrangements. Then, we used the ‘-km’ option of WGDI to map the obtained 17 protochromosomes of ancestral Musaceae karyotype (AMK) onto the chromosomes of the sampled species. In addition, the option ‘-sf’ was applied to quickly identify inter-chromosome rearrangements and document the shared rearrangements as well as relevant breakpoints. We also used the option ‘-fpd’ to obtain the corresponding rearrangement positions database and subsequently used the option ‘-fd’ to check if these rearrangements are shared among other genomes. Finally, phylogenetic relationships were inferred based on the sharing patterns of rearrangement breakpoints among species. To ensure transparency and reproducibility, a detailed, step-by-step description of the ancestral Musaceae karyotype reconstruction workflow—spanning ortholog identification, synteny-based ancestral contig assembly, chromosome reconstruction, and evolutionary breakpoint inference—is provided in Fig. S15, Supplementary Data, and the accompanying online documentation (https://github.com/SunPengChuan/Ancestral_Musaceae_Karyotype/blob/main/AMK_reconstruction_details/ancestor_reconstruction.md).

### Anthocyanin biosynthesis pathway

We search for the information about anthocyanin biosynthesis pathway, its upstream phenylpropanoid synthesis pathway, and flavonoid synthesis pathway on the KEGG PATHWAY Database [85]. The K numbers of all enzymes in the pathway were extracted. To identify structural genes encoding enzymes in the genomes, we used the software Eggnog-mapper for functional annotation of proteins. The results were processed by eggNOG-mapper Helper plugin in TBtools v2.042 [86] to obtain the K number library of structural genes, which was then used to calculate the copy numbers of structural gene. We sequenced the RNA from the bract and leaf of *M. exotica* and *M*. *beccarii*, and downloaded RNA data of bracts and leaves of *M*. *textilis* and *M*. *ornata* [20, 87]. The RNA data was aligned to each genome using Hisat2 v2.2.1 [88], followed by quantification of expression levels using featureCounts v2.0.6 [89] with parameters: -t exon -g gene_id -Q 10 -primary -s 0 -p. The expression level was then normalized in R v4.2.3 (https://www.r-project.org/) using the TPM method to eliminate the effects caused by differences in sequencing depth or gene lengths between samples. Finally, a heatmap of expression levels was generated using ChiPlot (https://www.chiplot.online/).

### Transcription factor–anthocyanin pathway gene correlation analysis

To investigate the regulatory relationships between transcription factors (TFs) and anthocyanin biosynthetic genes, we performed a genome-wide co-expression analysis based on normalized transcript abundance data. Expression matrices of TFs (MYB and bHLH families identified using iTAK v2.0.5 [35]) and anthocyanin pathway structural genes were obtained from RNA-seq datasets and quantified as TPM values.

Prior to analysis, genes with zero expression across all samples were removed to avoid spurious correlations. The expression matrices were then log₂-transformed [log₂(TPM + 1)] to stabilize variance and reduce the influence of extreme values. All matrices were transposed such that samples were treated as observations and genes as variables. Pairwise Pearson correlation coefficients (*r*) were calculated between each TF and each anthocyanin biosynthetic gene across all samples, and statistical significance was evaluated using corresponding *P*-values. The resulting correlation and *P*-value matrices were combined and filtered to identify high-confidence TF–target associations. Given the exploratory nature of this analysis, we applied thresholds of |*r*| ≥ 0.8 and *P* < 0.01 without multiple testing correction to capture robust co-expression relationships.

The correlation matrix was visualized as a heatmap using hierarchical clustering to reveal co-expression patterns among TFs and structural genes. In addition, significantly correlated TF–enzyme pairs were extracted to construct a co-expression network, where nodes represent genes and edges represent significant correlations, including both positive and negative associations. The resulting networks were visualized in Cytoscape v3.10.3 [90], enabling the identification of key regulatory modules and candidate TFs potentially involved in anthocyanin biosynthesis.

## Acknowledgements

This work was financially supported by the National Natural Science Foundation of China (No. 32570436, 32070237, 31261140366, 32500199) and the Guangdong Basic and Applied Basic Research Foundation (2026A1515010301). We thank Yu-Ying Zhou and Yu-Shi Ye for their help in sample collection.

## Author contributions

H.-R.H., and X.-J.G.conceived the project. X.-J.G. and N.F. collected the materials. N.F., P.-C.S., X.L., X.-F.W., T.J.L., Y.-B.W., W.-M.L., T.-W.X., Z.-F.W., and X.-N.L. performed the analyses. N.F., P.-C.S., and X.L. wrote the manuscript. X.-F.W., H.-R.H., X.-J.G., and M.R. revised the manuscript. All authors approved the final manuscript.

## Data availability

All the raw sequence data were deposited in the Genome Sequence Archive in the National Genomics Data Center (NGDC), China National Center for Bioinformation (CNCB) with the accession number of CRA030696 under BioProject PRJCA046700 (https://ngdc.cncb.ac.cn/). The final T2T genome and its annotation file were deposited in the Science Data Bank at https://doi.org/10.57760/sciencedb.28970.

