## Supplemental Figures for "Ancestral Musaceae karyotype reconstruction provides insights into chromosome evolution and bract coloration"

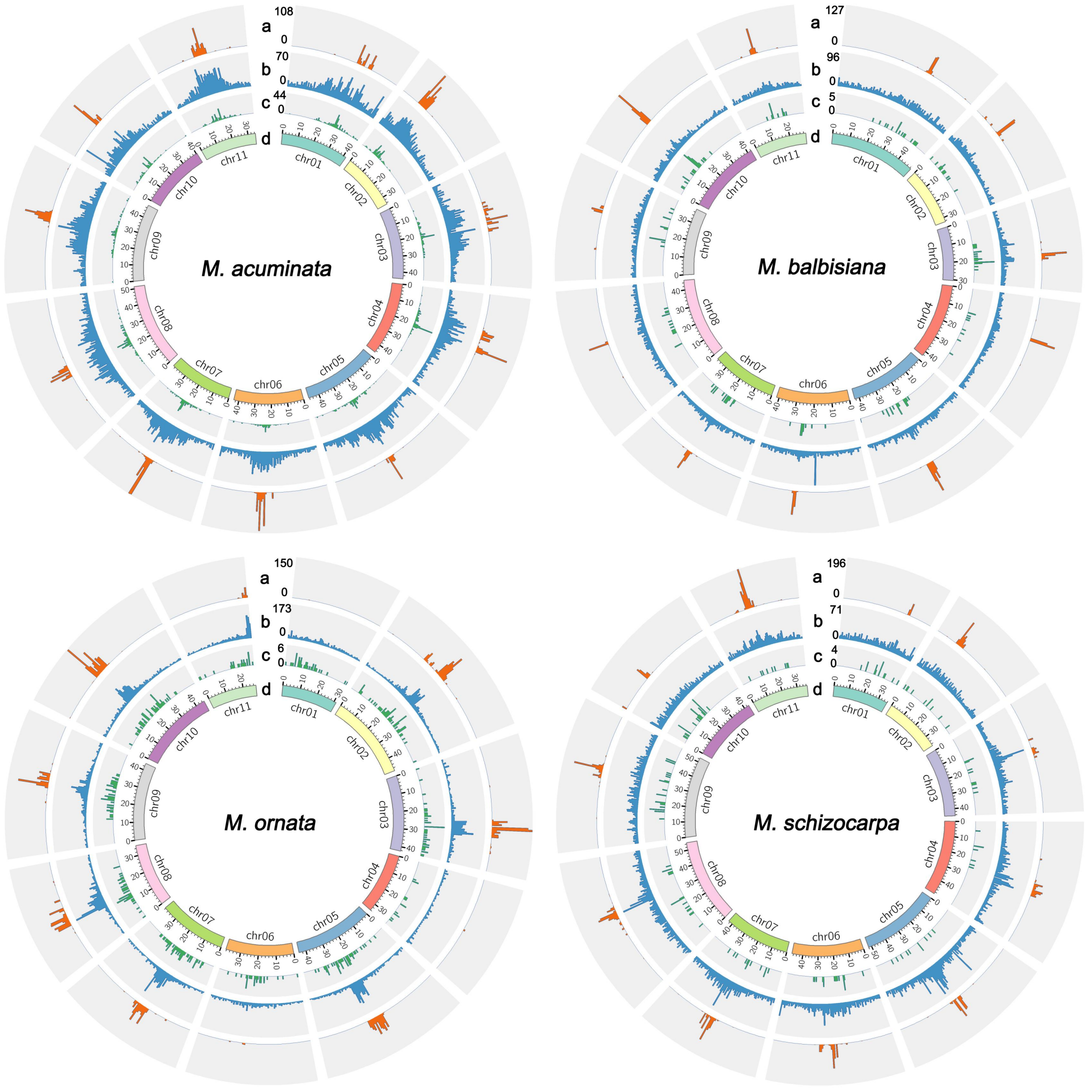


**Figure S1. The feature of centromeric regions of four sect. *Musa* species.** The circles from outer to inner represent the distribution of (a) *Nanica* sequences, (b) all tandem repeats with unit lengths of 100-200 bp, (c) *Mac-Cen183*, and (d) genomic chromosomes. 500 kb per window. Mac: *M. acuminata*.

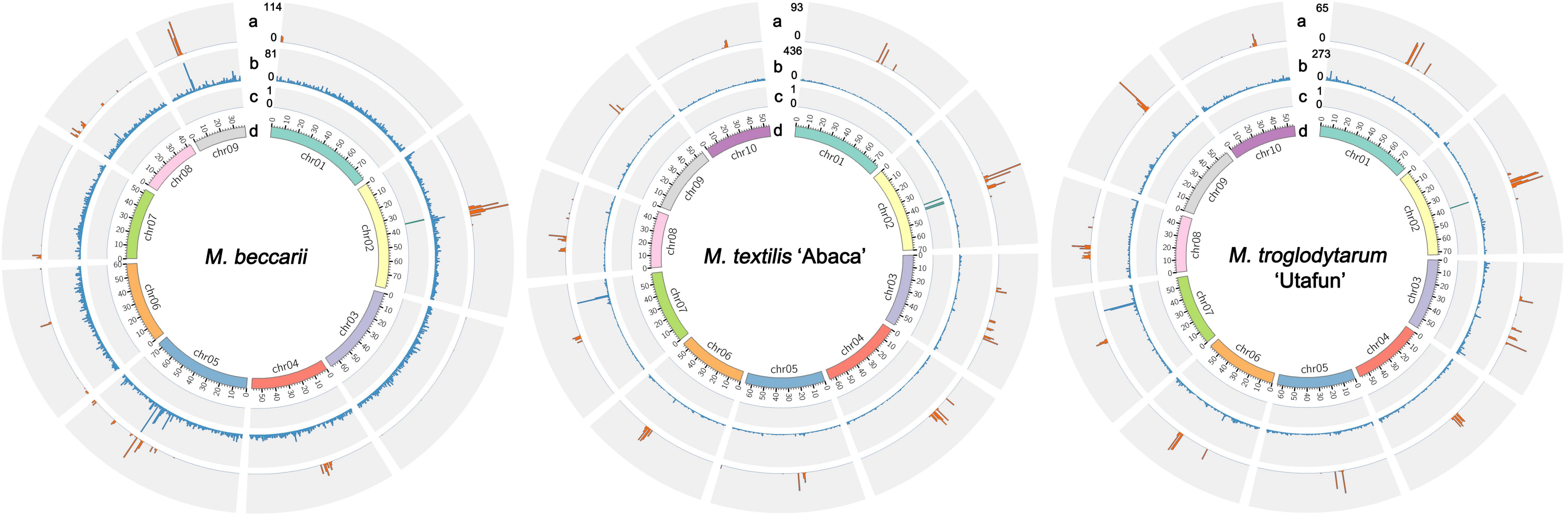


**Figure S2. The feature of centromeric regions of three sect. *Callimusa* species.** The circles from outer to inner represent the distribution of (a) *Nanica* sequences, (b) all tandem repeats with unit lengths of 100-200 bp, (c) *Mex-Cen186*, and (d) genomic chromosomes. 500 kb per window. Mex: *M. exotica*.

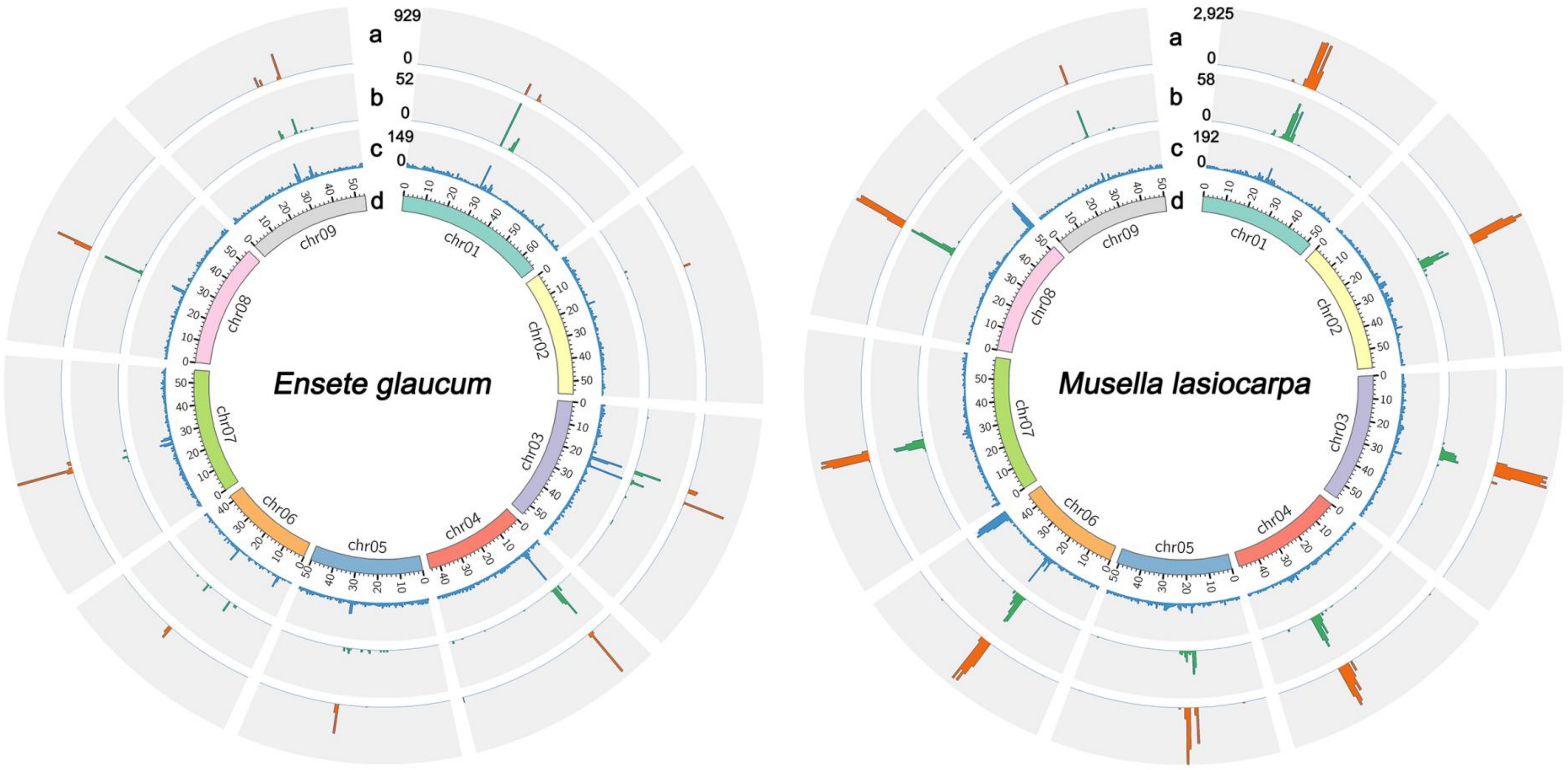


**Figure S3. The feature of centromeric regions of *E. glaucum* and *M. lasiocarpa*.** The outer to inner circles represent the distribution of (a) *Egcen*, (b) *Nanica*, (c) all tandem repeats with unit lengths of 100-200 bp, and (d) genomic chromosomes. 500 kb per window.

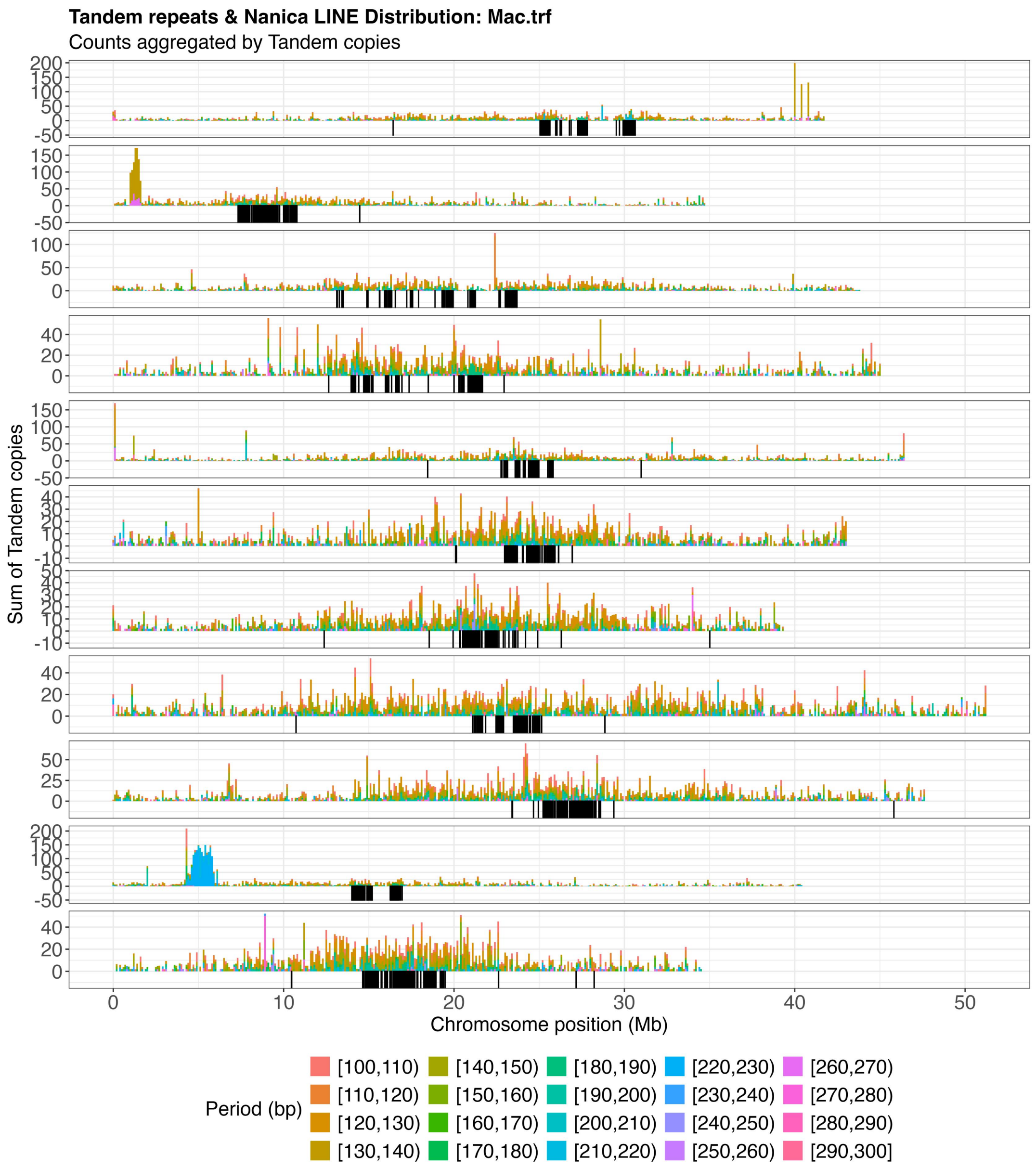


**Figure S4A. Distribution of *Nanica* LINE elements and tandem repeats (TRs) in sect. *Musa* species *M. acuminata*.** Genome-wide distributions of *Nanica* LINE elements and tandem repeats (TRs) were analyzed across chromosomes. TR copy numbers were calculated using 100 kb sliding windows. Tandem repeats with unit lengths <100 bp or >300 bp were excluded. The remaining TRs were further grouped into categories based on repeat unit length (10 bp intervals), and their copy numbers were summarized and visualized across chromosomes using distinct colors. The distribution of *Nanica* LINE elements is shown as a black line along each chromosome.

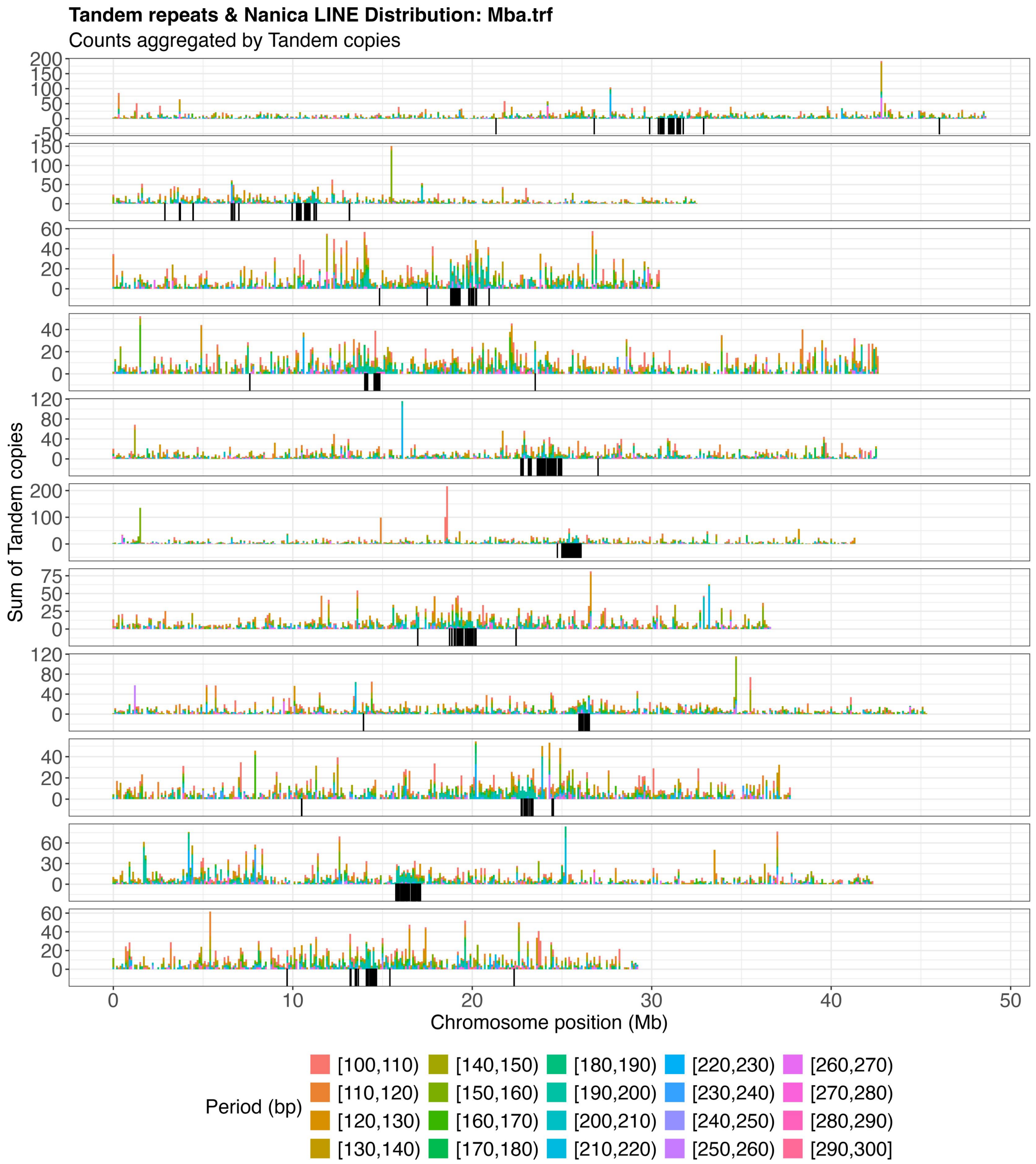


**Figure S4B. Distribution of *Nanica* LINE elements and tandem repeats (TRs) in sect. *Musa* species *M. balbisiana*.**

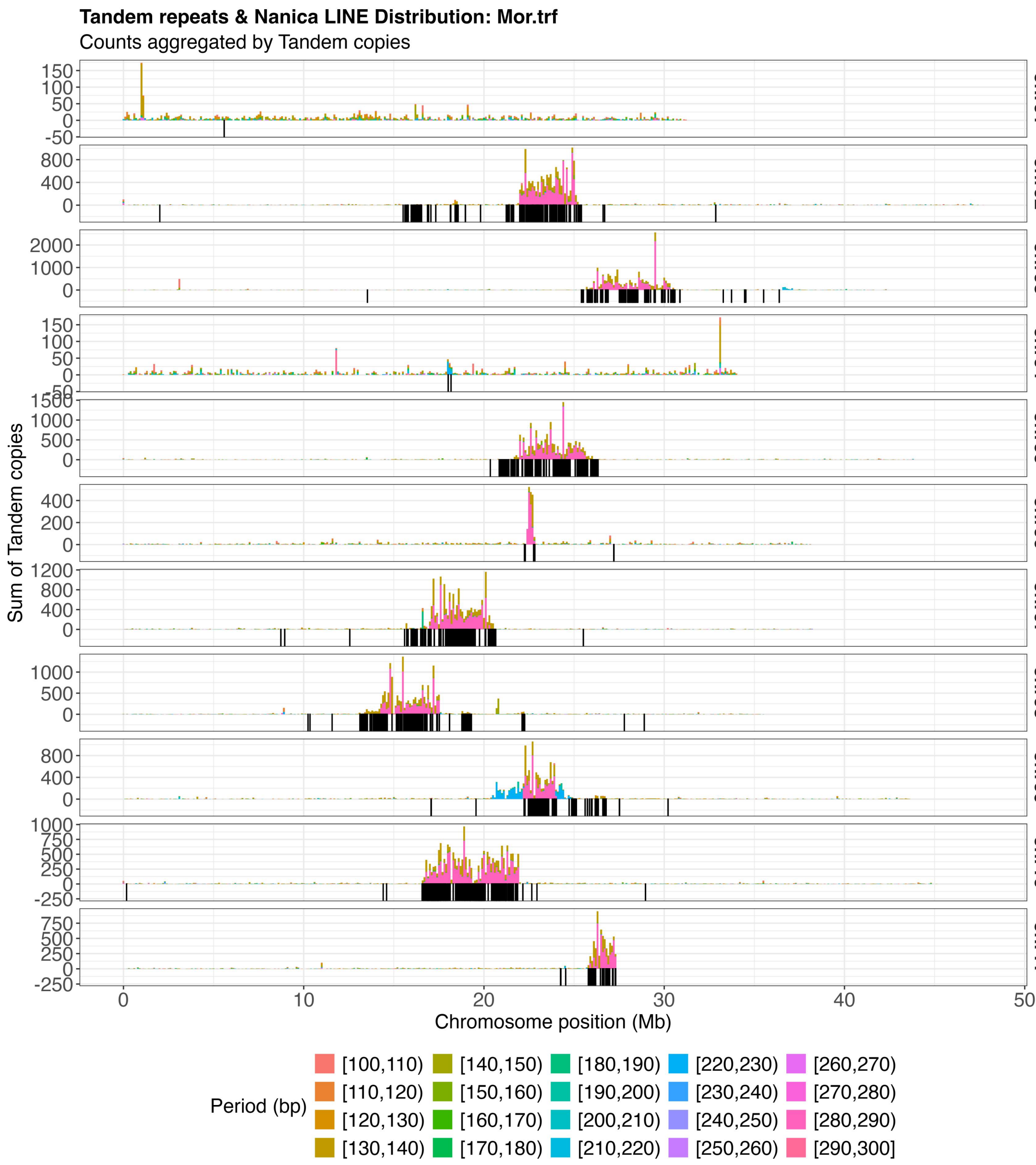


**Figure S4C. Distribution of *Nanica* LINE elements and tandem repeats (TRs) in sect. *Musa* species *M. ornata*.**

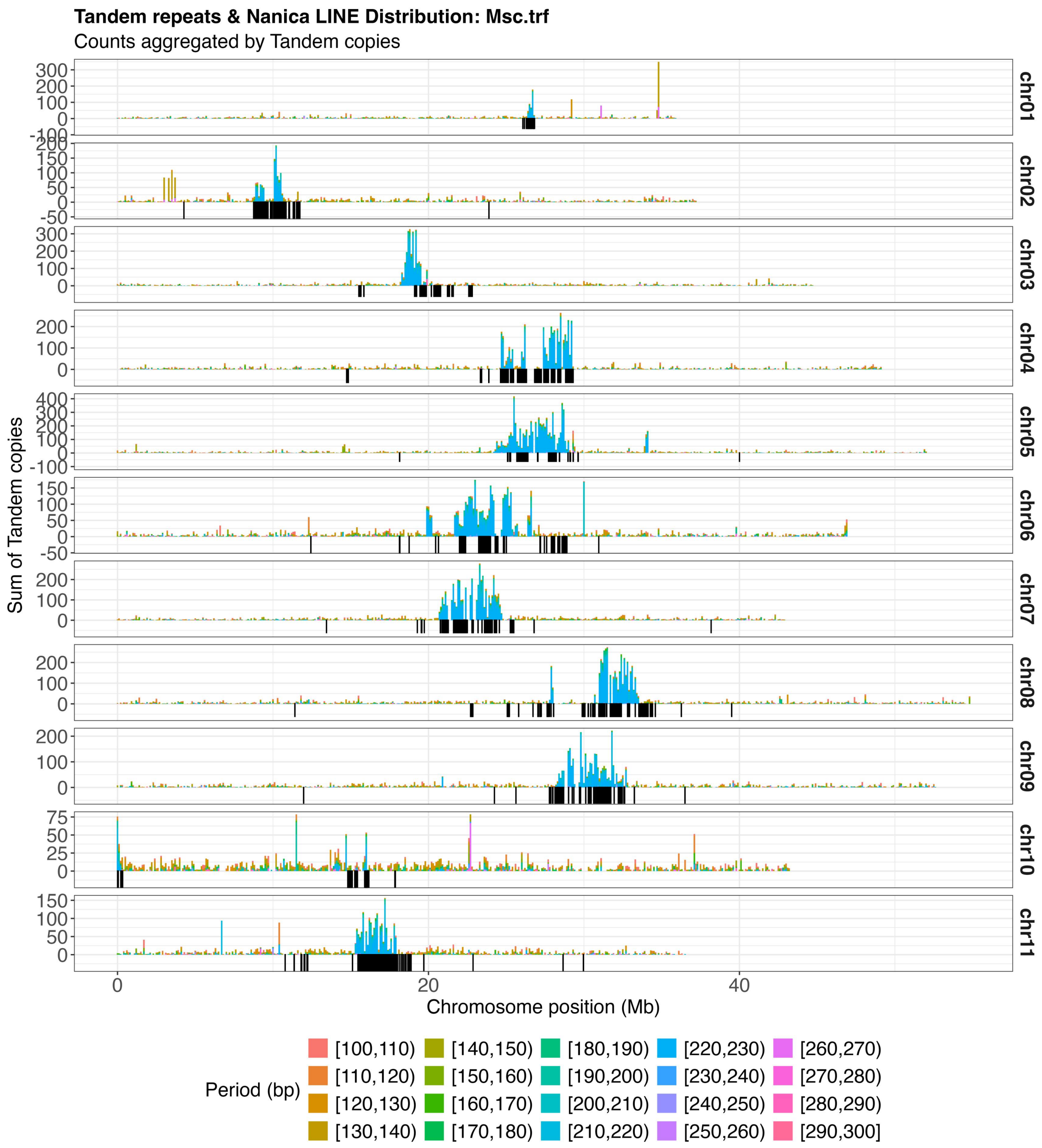


**Figure S4D. Distribution of *Nanica* LINE elements and tandem repeats (TRs) in sect. *Musa* species *M. schizocarpa*.**

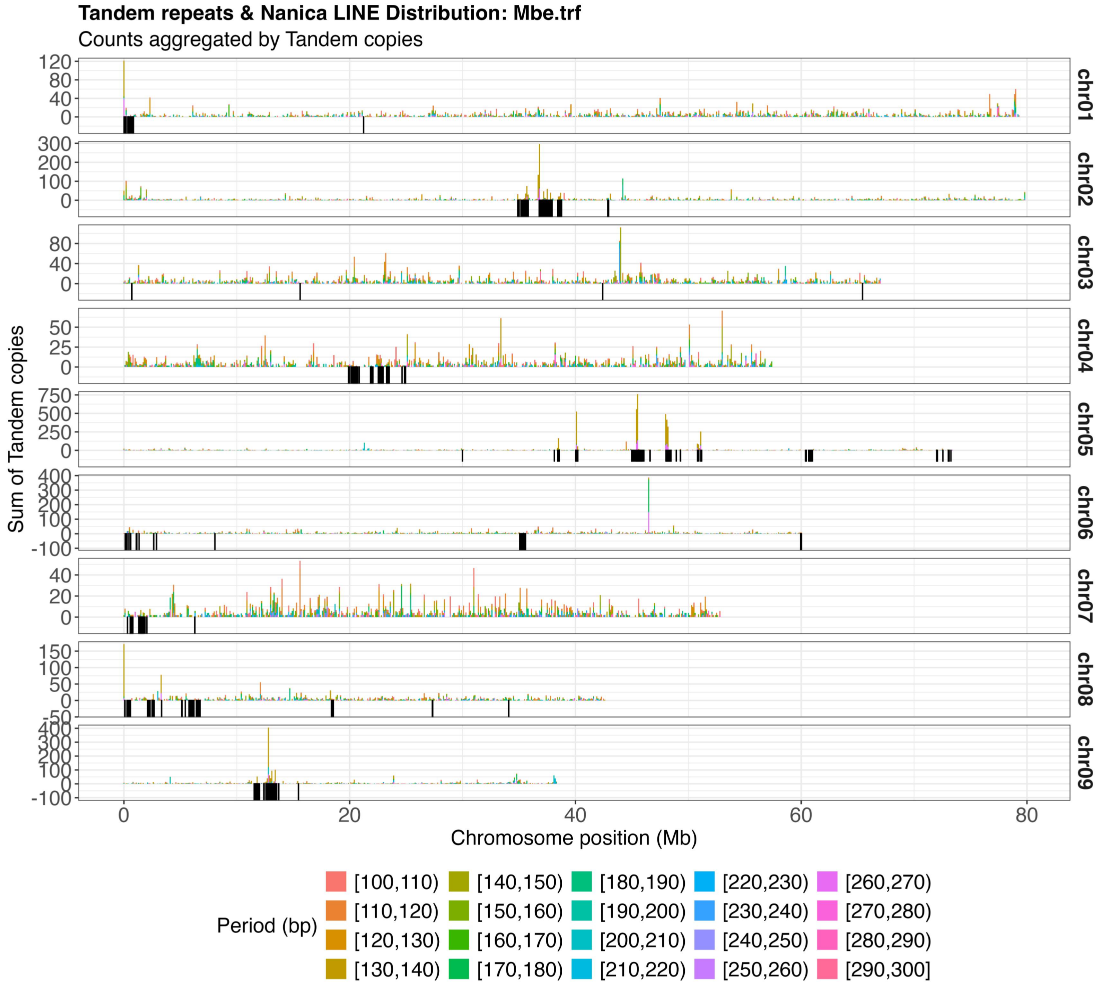


**Figure S4E. Distribution of *Nanica* LINE elements and tandem repeats (TRs) in sect. *Callimusa* species *M. beccarii*.**

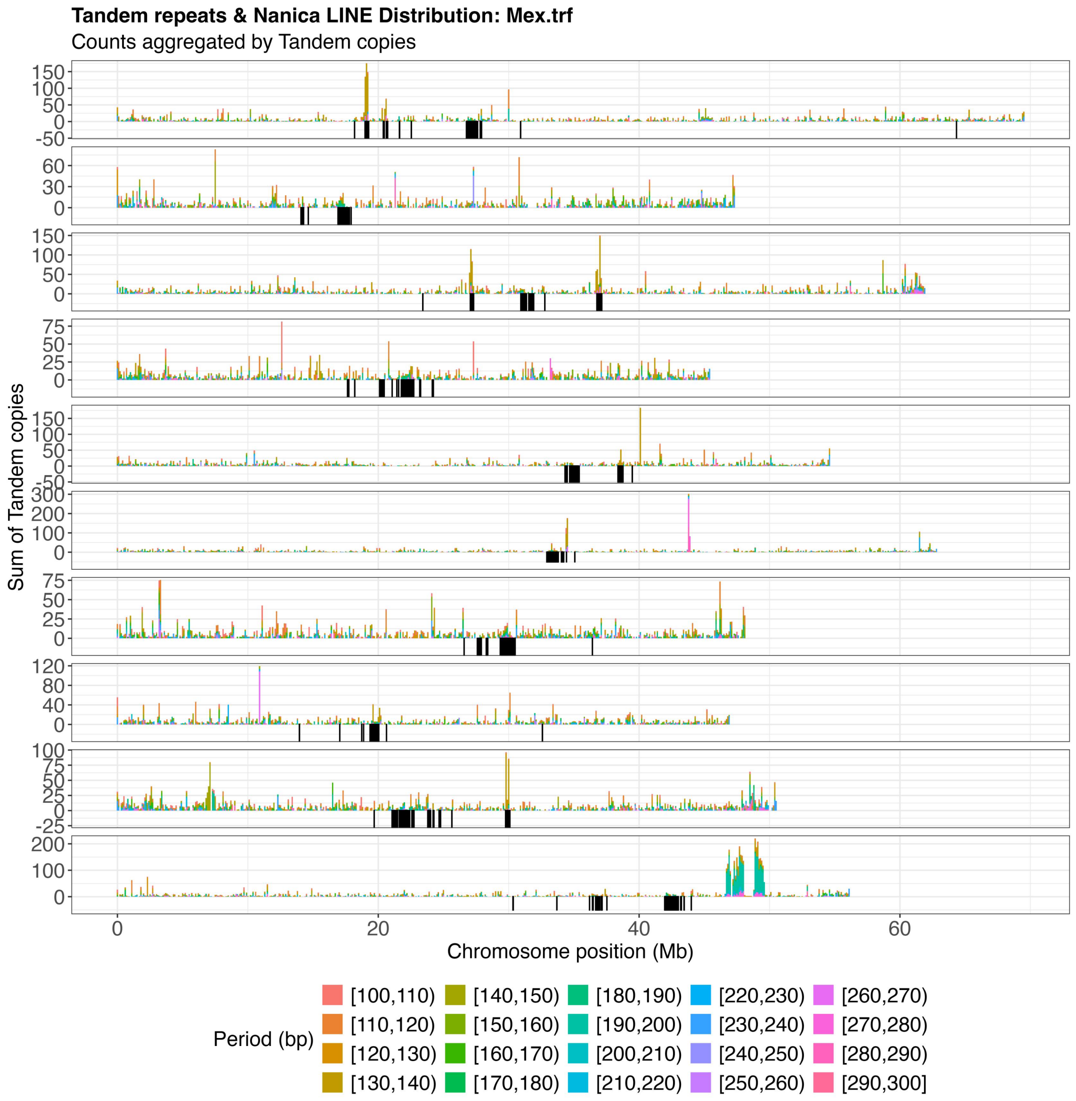


**Figure S4F. Distribution of *Nanica* LINE elements and tandem repeats (TRs) in sect. *Callimusa* species *M. exotica*.**

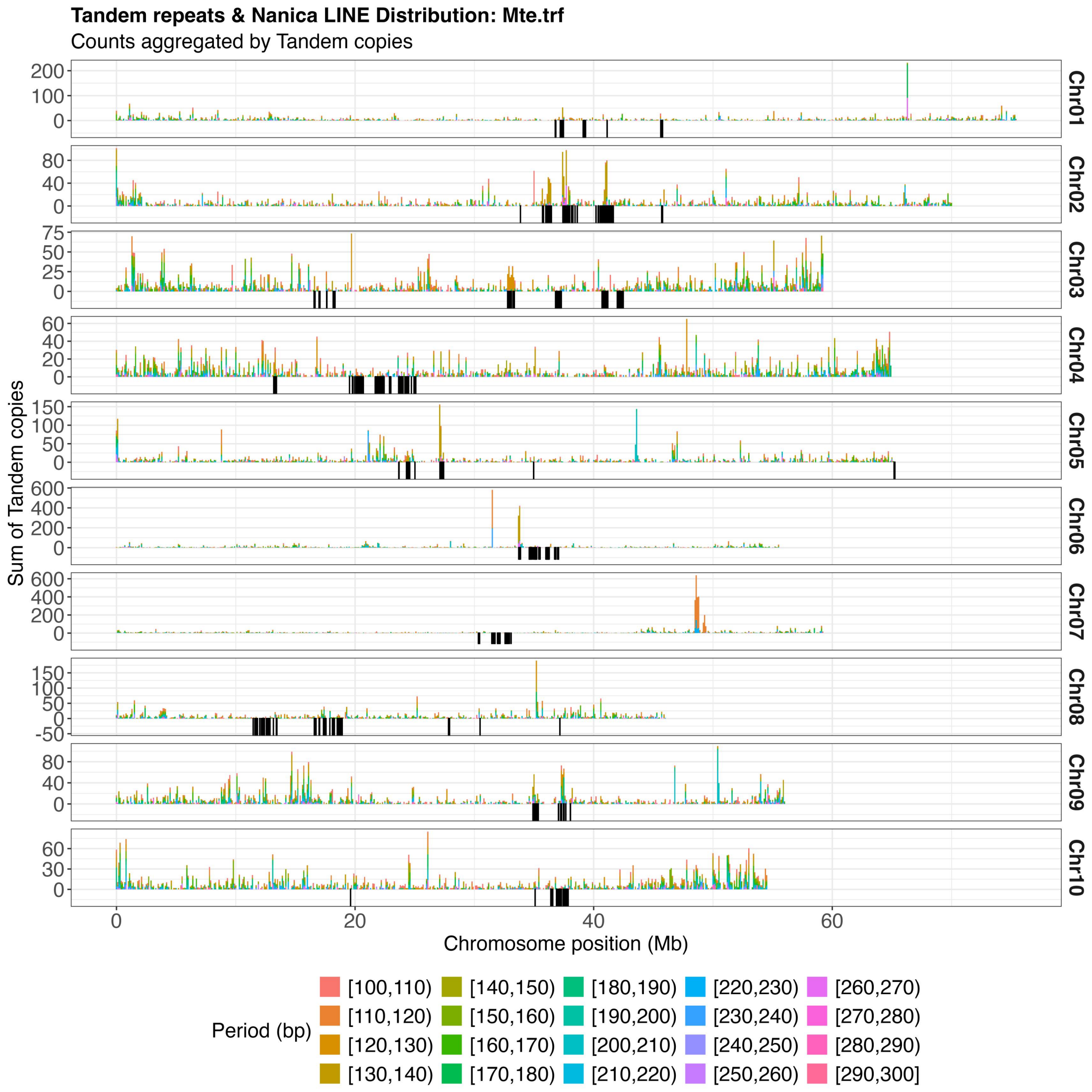


**Figure S4G. Distribution of *Nanica* LINE elements and tandem repeats (TRs) in sect. *Callimusa* species *M. textilis*.**

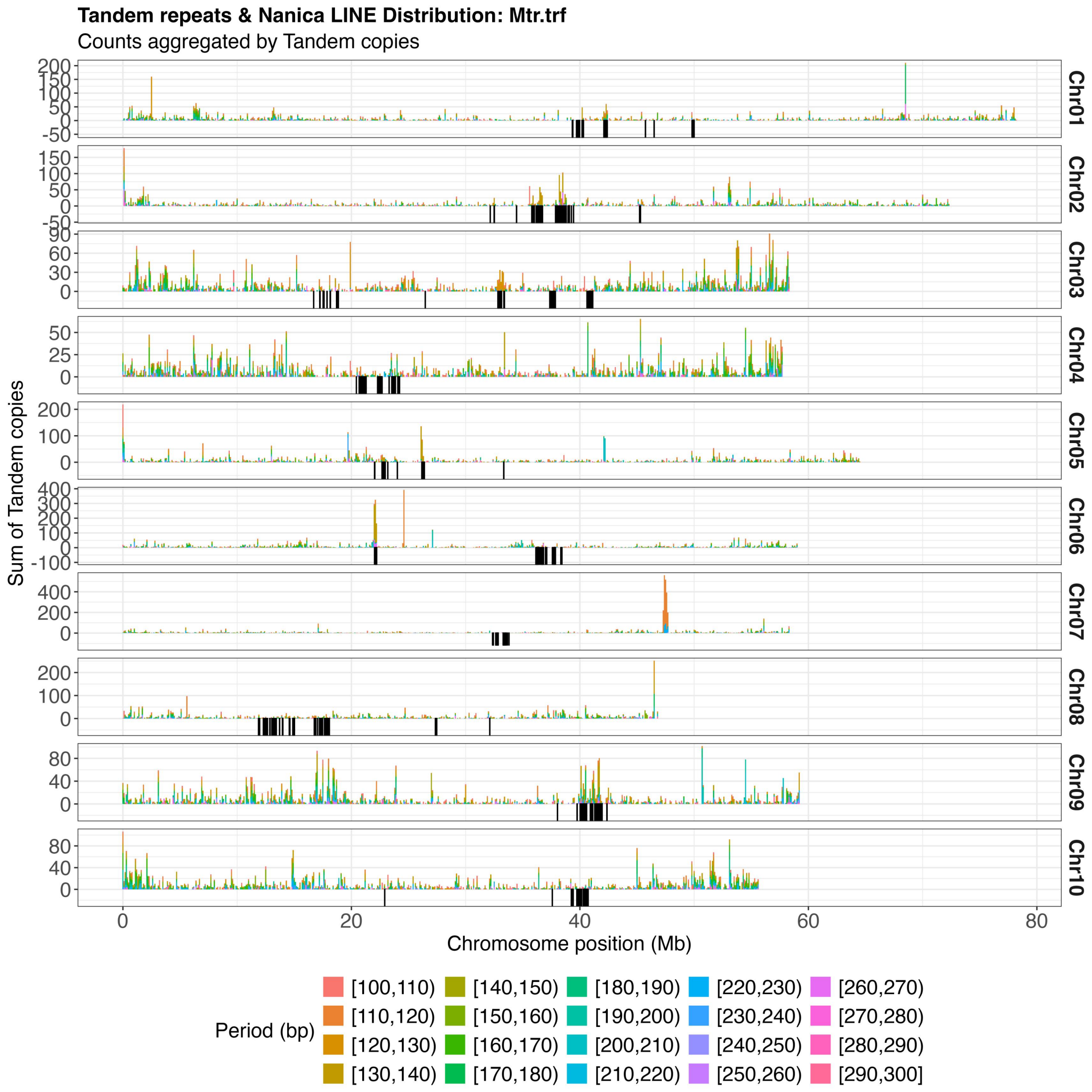


**Figure S4H. Distribution of *Nanica* LINE elements and tandem repeats (TRs) in sect. *Callimusa* species *M. troglodytarum*.**

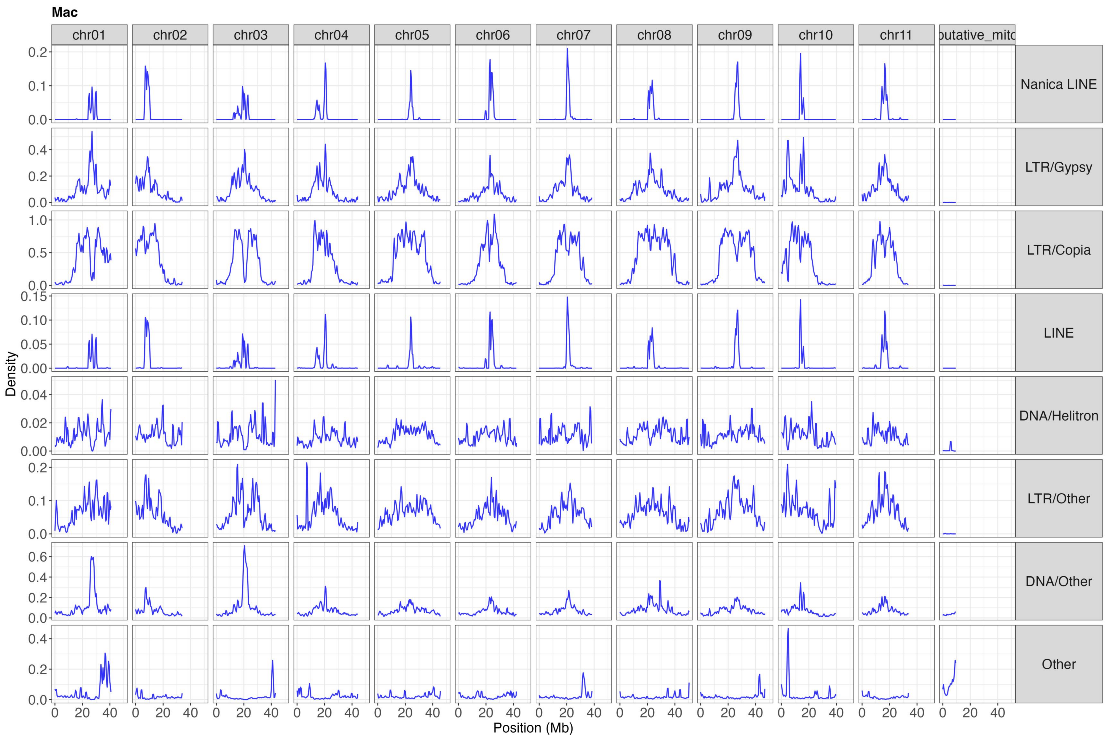


**Figure S5A. Transposable element landscape of sect. *Musa* species *M. acuminata*.** Genome-wide chromosomal distributions of transposable elements (TEs) are shown based on their density (coverage ratio). Putative centromeric regions are characterized by a pronounced enrichment of *Gypsy* elements and a concomitant depletion of *Copia* elements. Notably, these regions also show strong accumulation of *Nanica* LINE elements (a subclass of LINEs), supporting their potential role as indicators of centromeric regions in *Musa*.

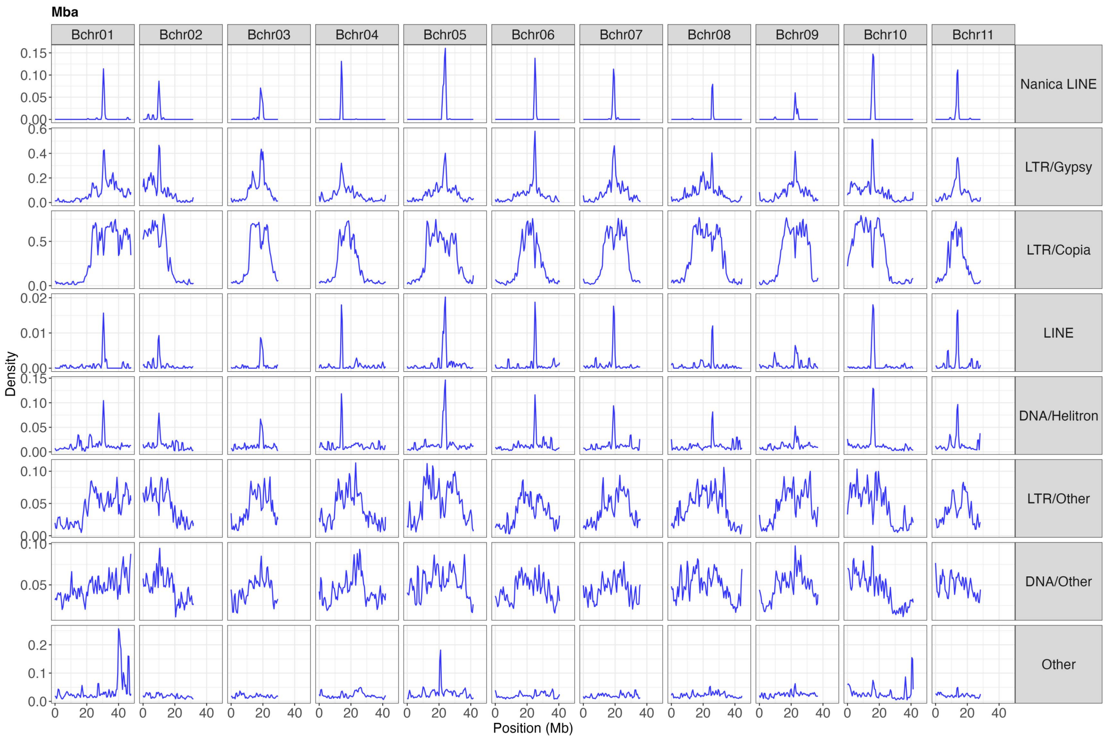


**Figure S5B. Transposable element landscape of sect. *Musa* species *M. balbisiana*.**

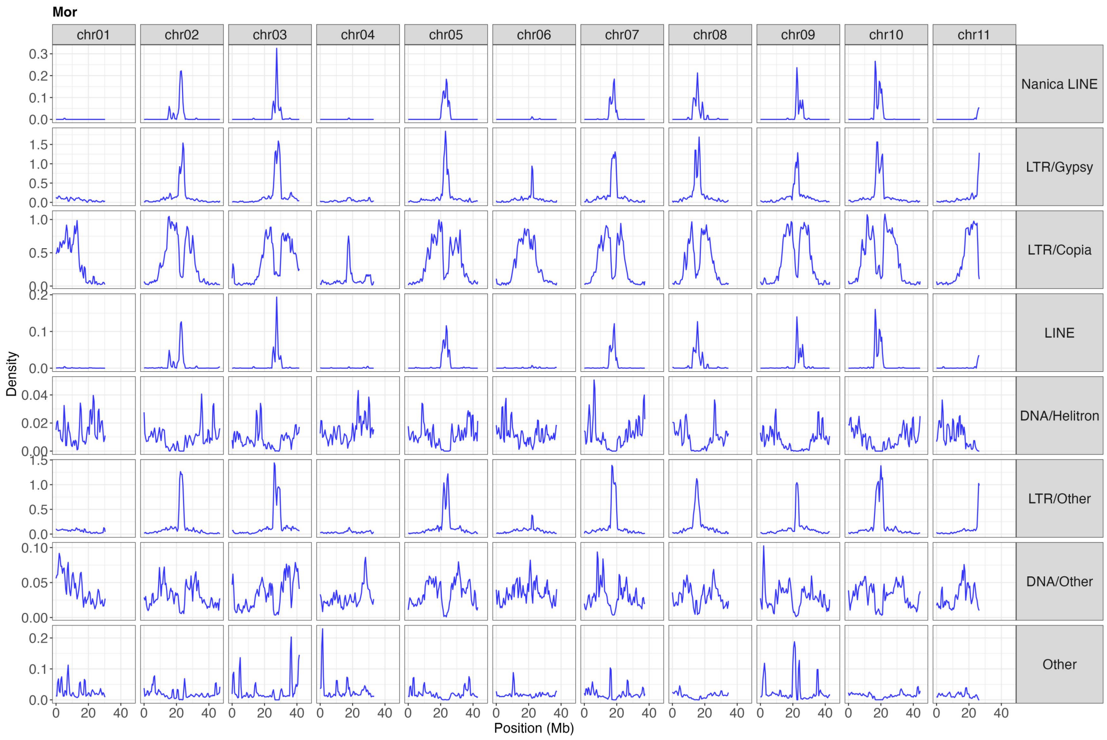


**Figure S5C. Transposable element landscape of sect. *Musa* species *M. ornata*.**

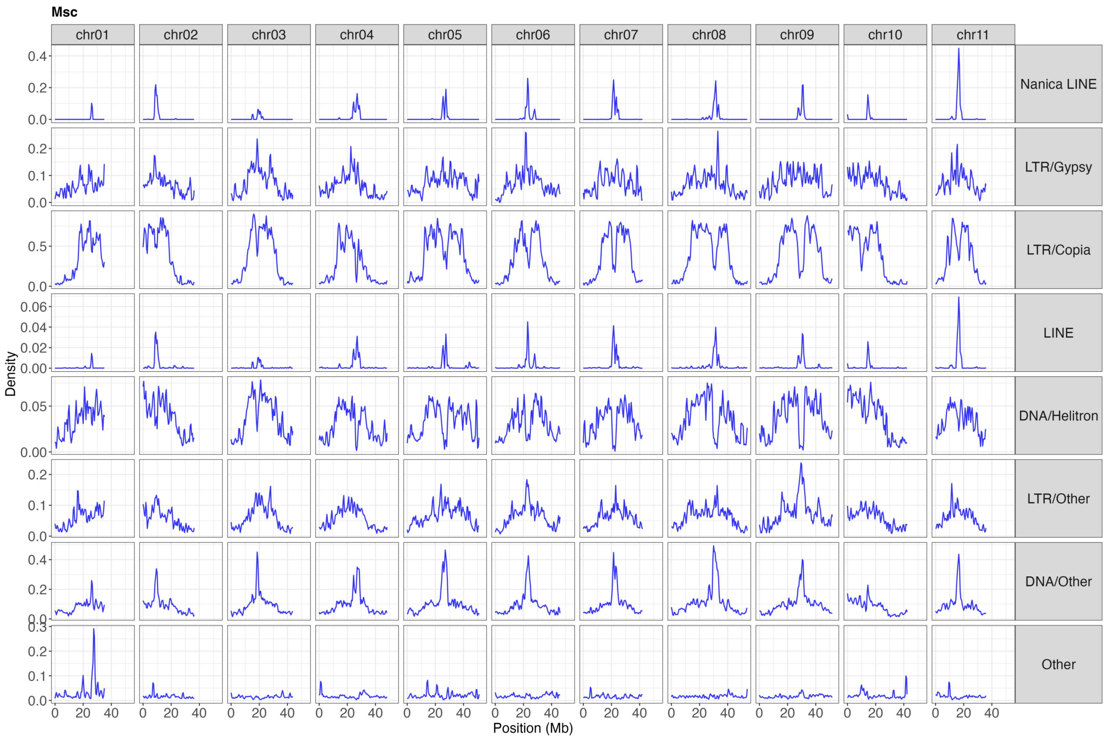


**Figure S5D. Transposable element landscape of sect. *Musa* species *M. schizocarpa*.**

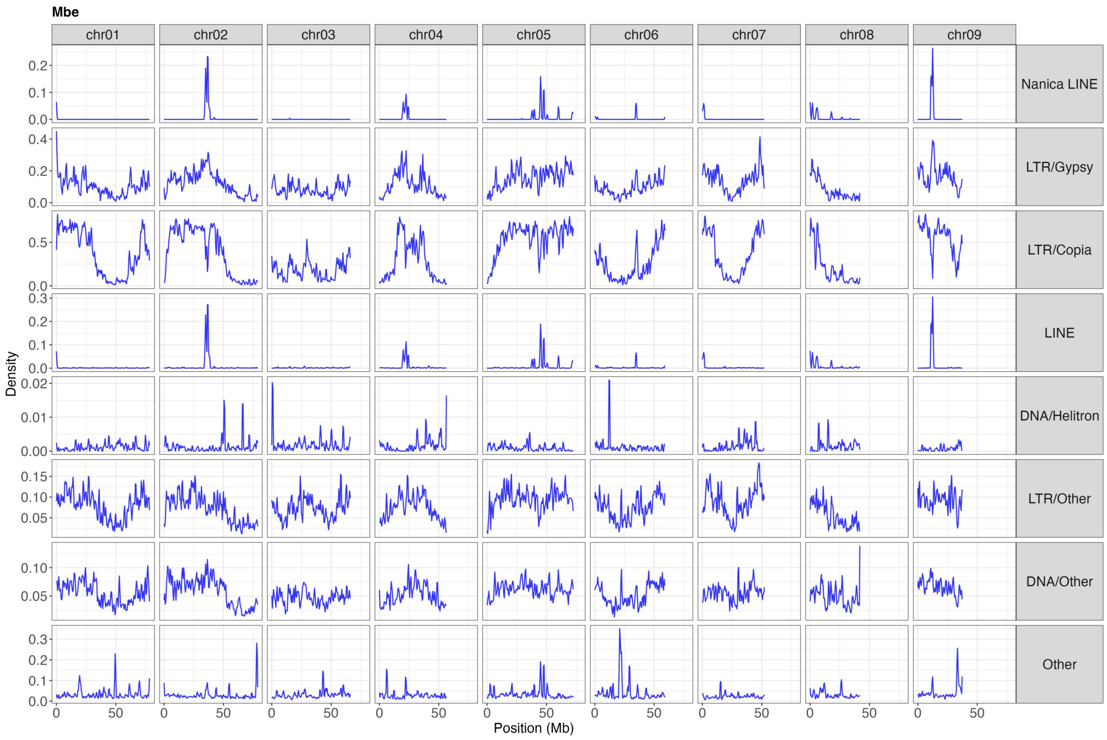


**Figure S5E. Transposable element landscape of sect. *Callimusa* species *M. beccarii*.**

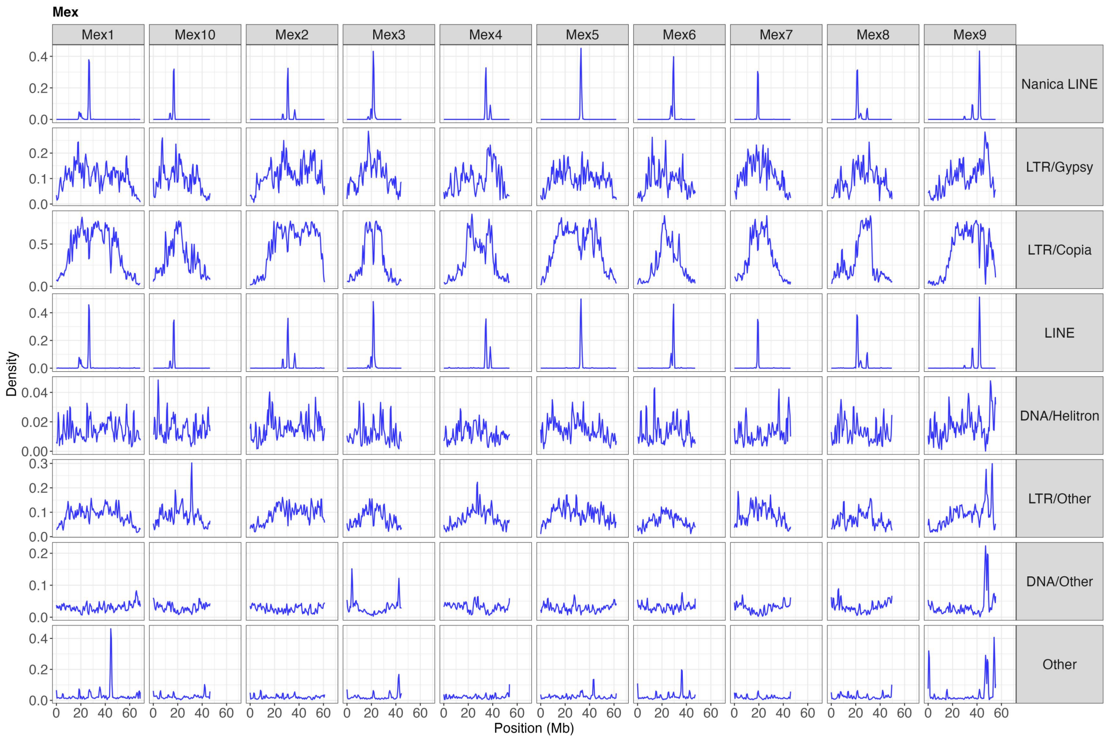


**Figure S5F. Transposable element landscape of sect. *Callimusa* species *M. exotica*.**

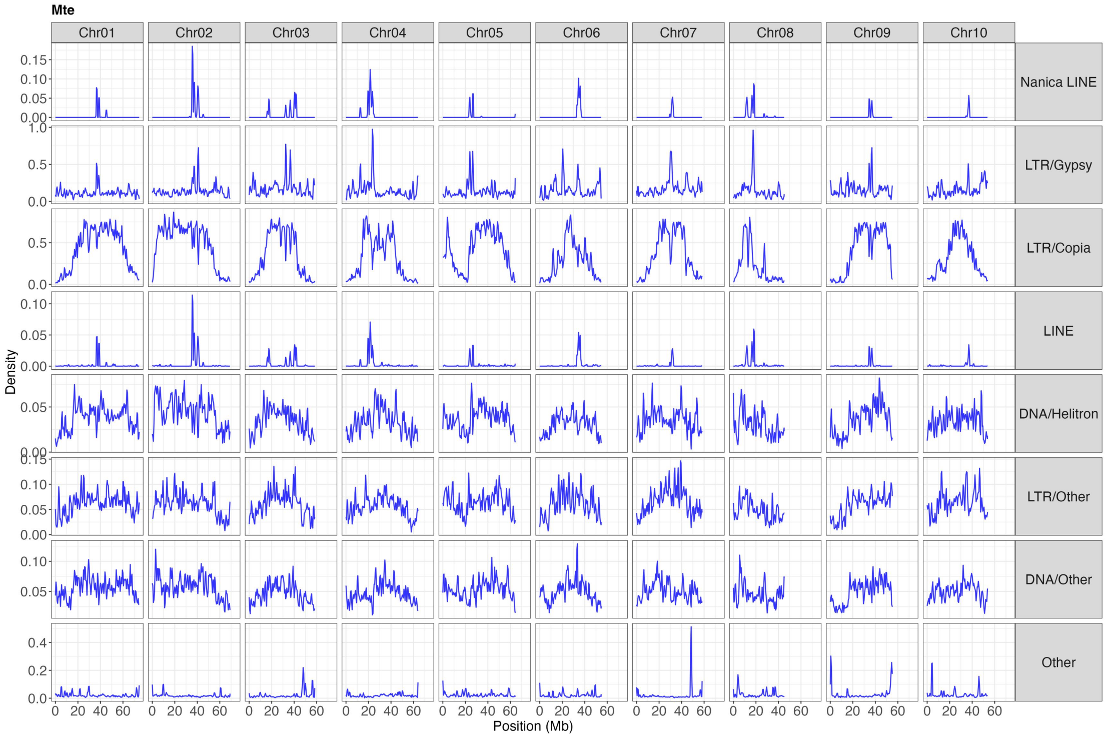


**Figure S5G. Transposable element landscape of sect. *Callimusa* species *M. textilis*.**

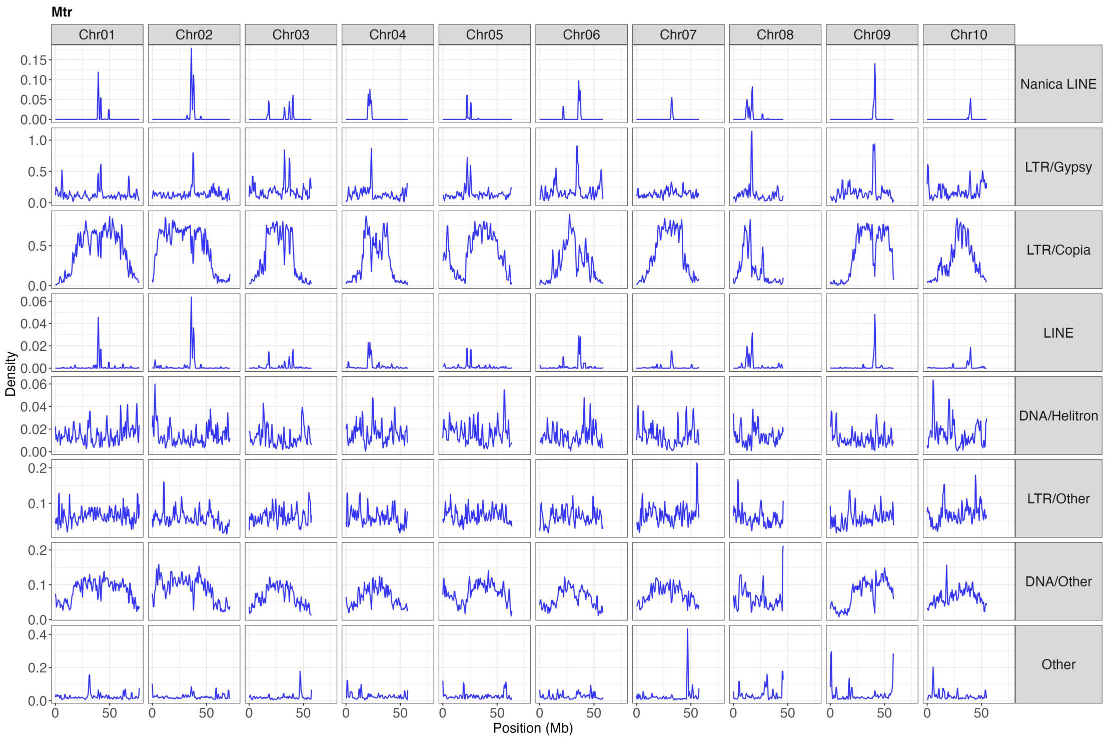


**Figure S5H. Transposable element landscape of sect. *Callimusa* species *M. troglodytarum*.**

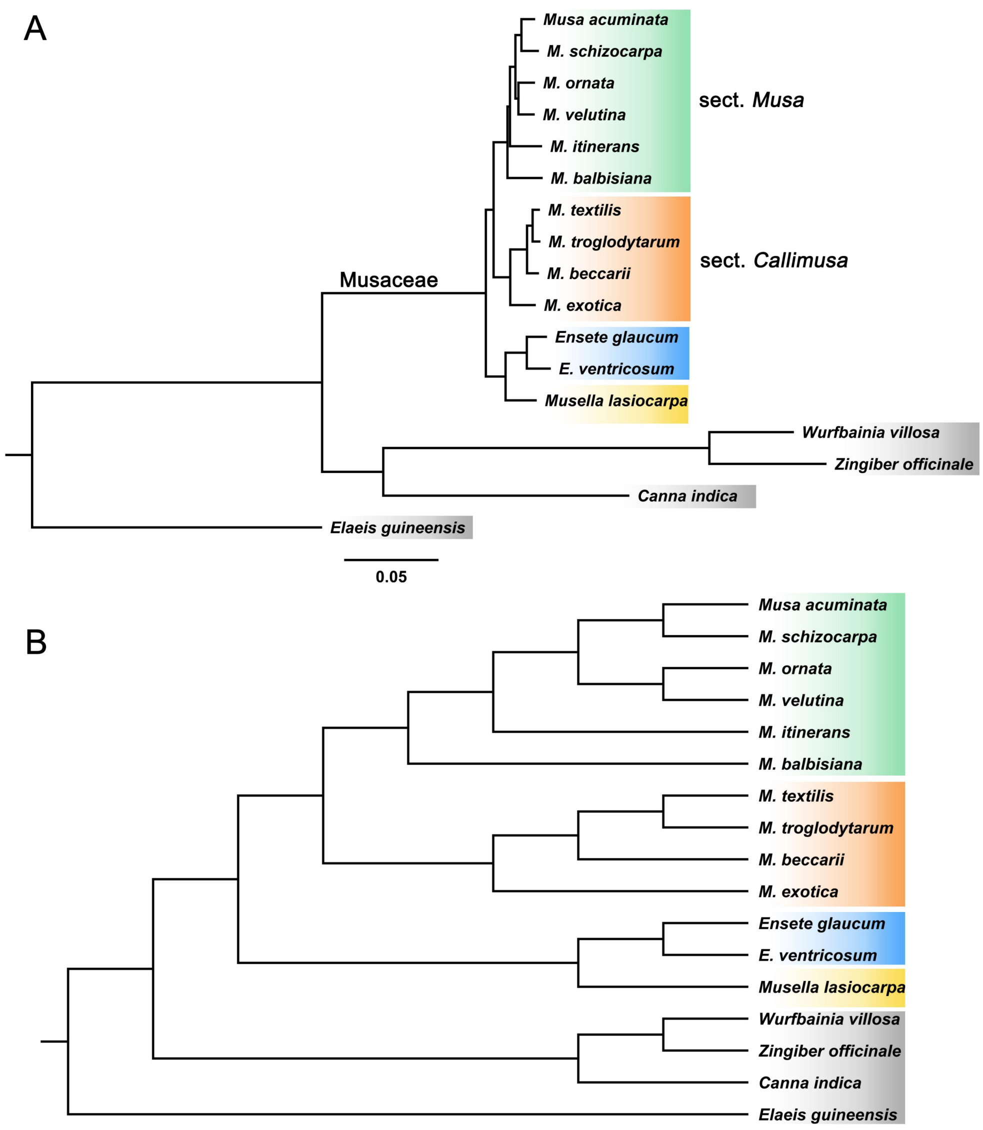


**Figure S6. Phylogenetic tree inferred from (A) concatenated method and (B) coalescent-based method. All branches are 100% supported.**

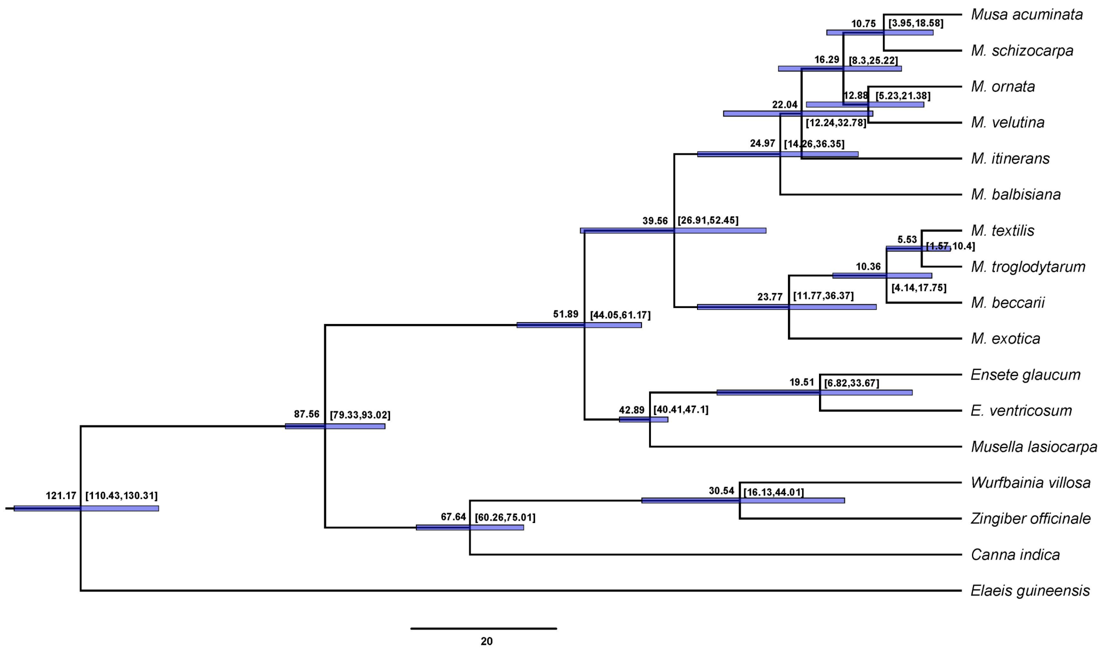


**Figure S7. The result of divergence time estimation.** The mean age (million years ago) was shown on the left side of the node and the 95% highest posterior density (HPD) on the right.

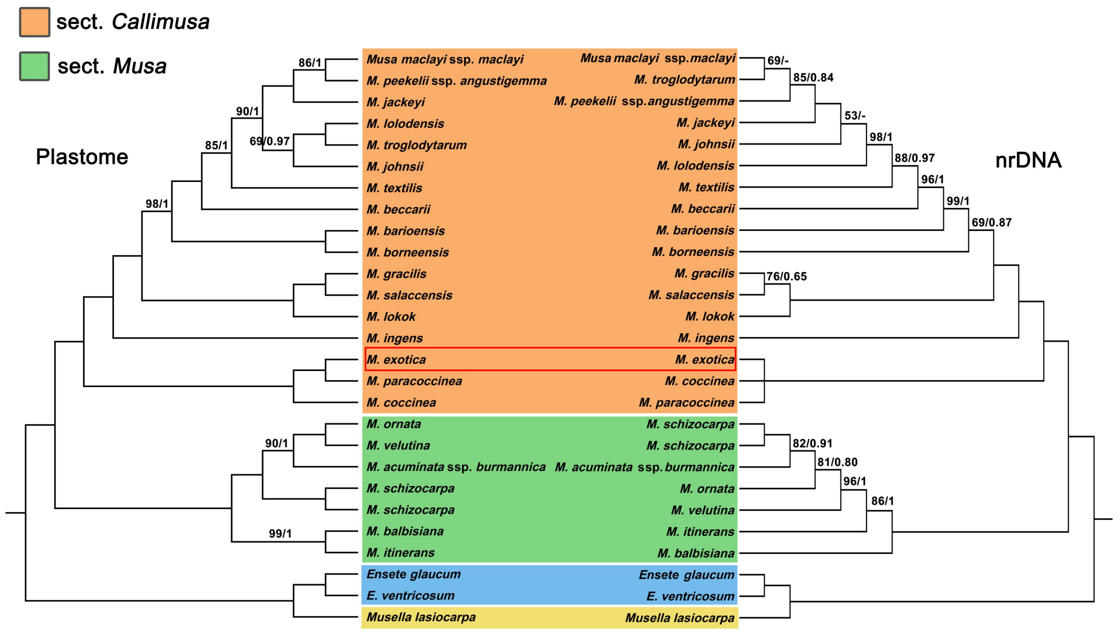


**Figure S8. *Musa* phylogenetic trees inferred from the plastome and nrDNA dataset.** Bootstrap (BS) values and posterior probabilities (PP) were shown on the branches except 100/1. Clade was set to polytomy when BS<50 and PP<0.5, '-' indicates PP under 0.5.

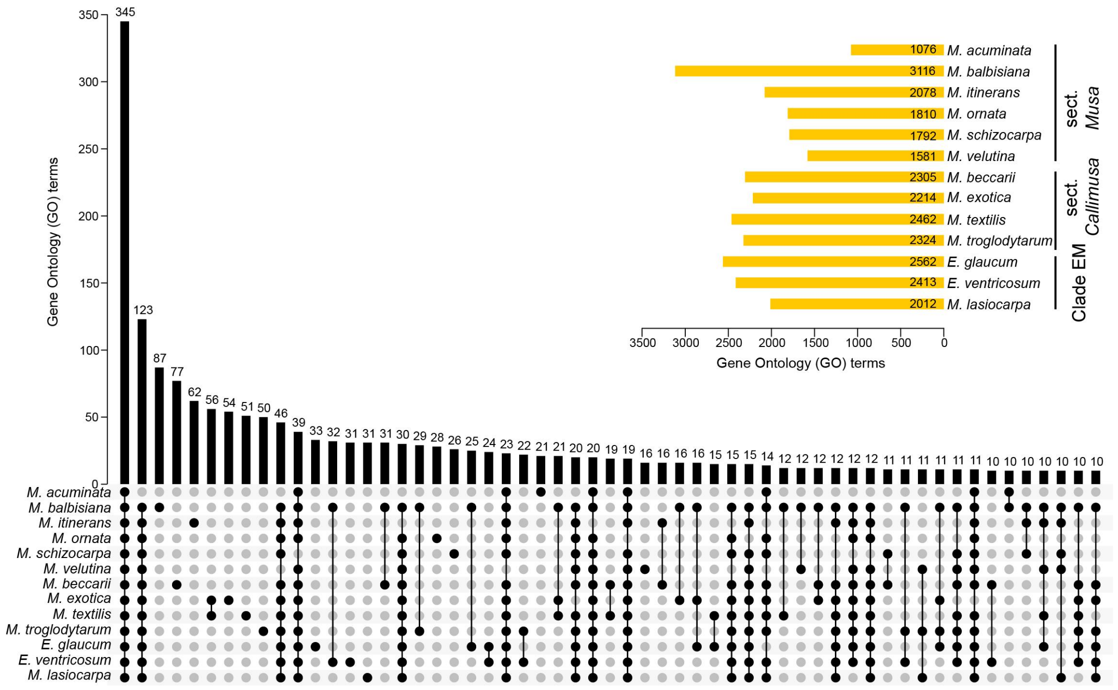


**Figure S9. Functional diversity of species-specific gene families across Musaceae revealed by GO enrichment analysis.** Upset plot showing the distribution and overlap of enriched Gene Ontology (GO) terms associated with species-specific gene families across Musaceae species. The bar plot (top) indicates the number of GO terms shared among different species combinations, while the connected dot matrix (bottom) represents the corresponding species intersections. The horizontal bar chart (upper right) summarizes the total number of enriched GO terms for each species, ranging from 1,076 to 3,116 categories, highlighting substantial functional diversity and pronounced lineage-specific enrichment patterns among species. Visualization was performed in TBtools-II.

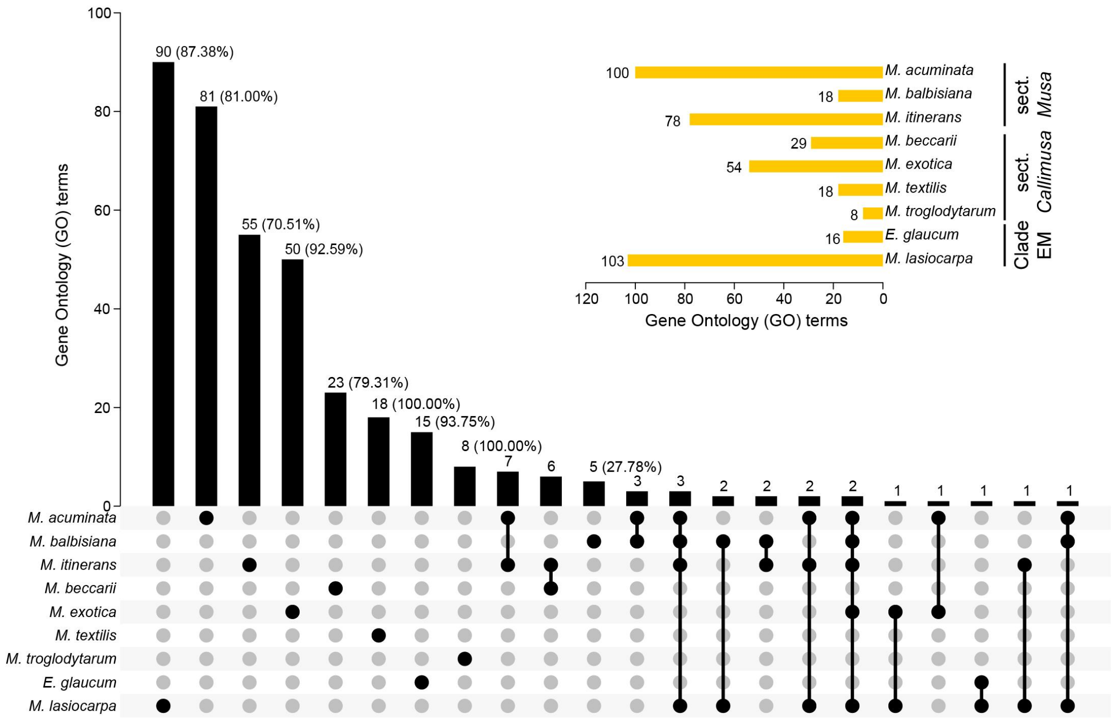


**Figure S10. Species-specific functional divergence in Musaceae revealed by GO enrichment analysis.** Upset plot showing the distribution and overlap of significantly enriched Gene Ontology (GO) terms associated with species-specific gene families across Musaceae species. The bar plot (top) indicates the number of GO terms shared among different species combinations, while the connected dot matrix (bottom) represents the corresponding species intersections. The horizontal bar chart (upper right) summarizes the total number of significantly enriched GO terms for each species, with ~70.51%–100.00% being unique to individual species, collectively highlighting strong species-specific functional divergence among lineages. Visualization was performed in TBtools-II.

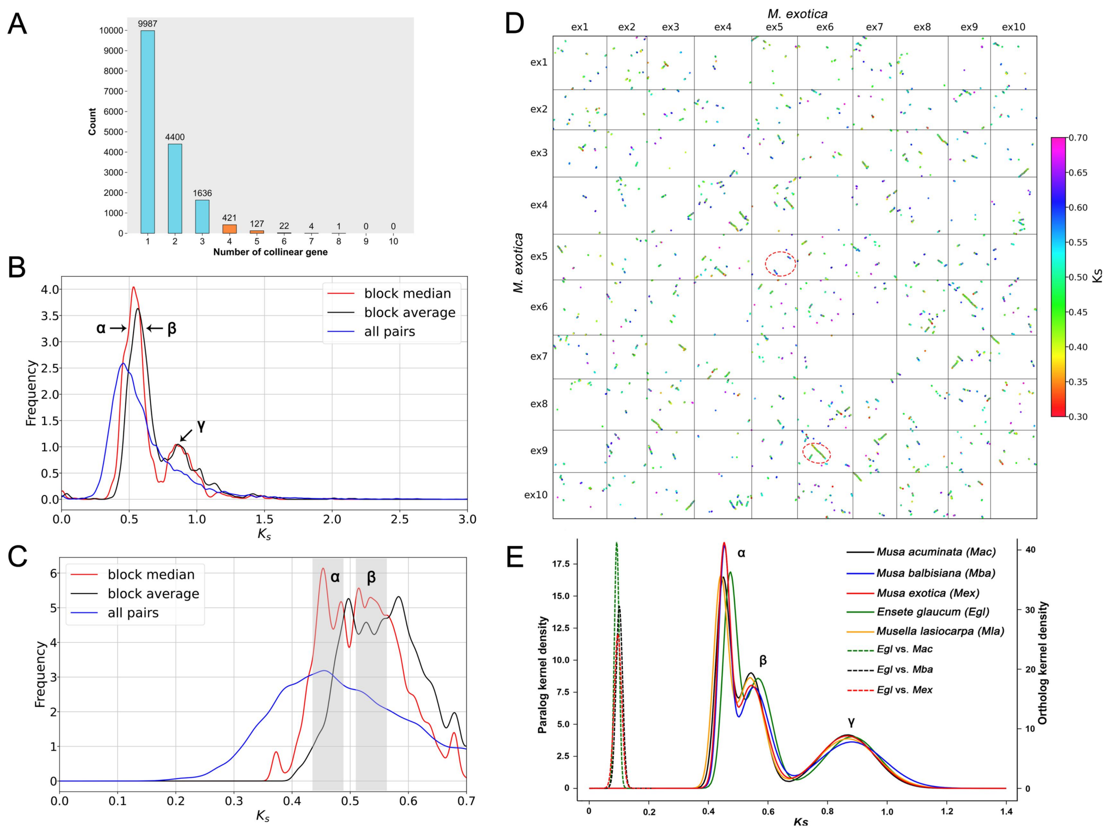


**Figure S11. Whole-genome duplication (WGD) analyses in Musaceae. (A)** Distribution of collinear gene counts per syntenic block in Musa exotica. **(B)** Ks (synonymous substitution rate) frequency distributions of collinear gene pairs and syntenic blocks across a broad range (Ks = 0–3), showing major peaks corresponding to ancient duplication events. **(C)** Zoomed-in Ks distribution (Ks = 0–0.7) in *M. exotica*, highlighting more recent duplication signals and resolving overlapping peaks. **(D)** Intraspecific synteny dot plot of *M. exotica*, with collinear gene pairs colored by Ks values (0.3–0.7). Two representative clusters of syntenic blocks (highlighted by dashed circles) illustrate duplicated regions derived from distinct WGD events. **(E)** Ks distribution of paralogous (solid lines) and orthologous (dashed lines) gene pairs among representative Musaceae species, revealing three shared ancient WGD events (α, β, and γ) and supporting their occurrence prior to species divergence.

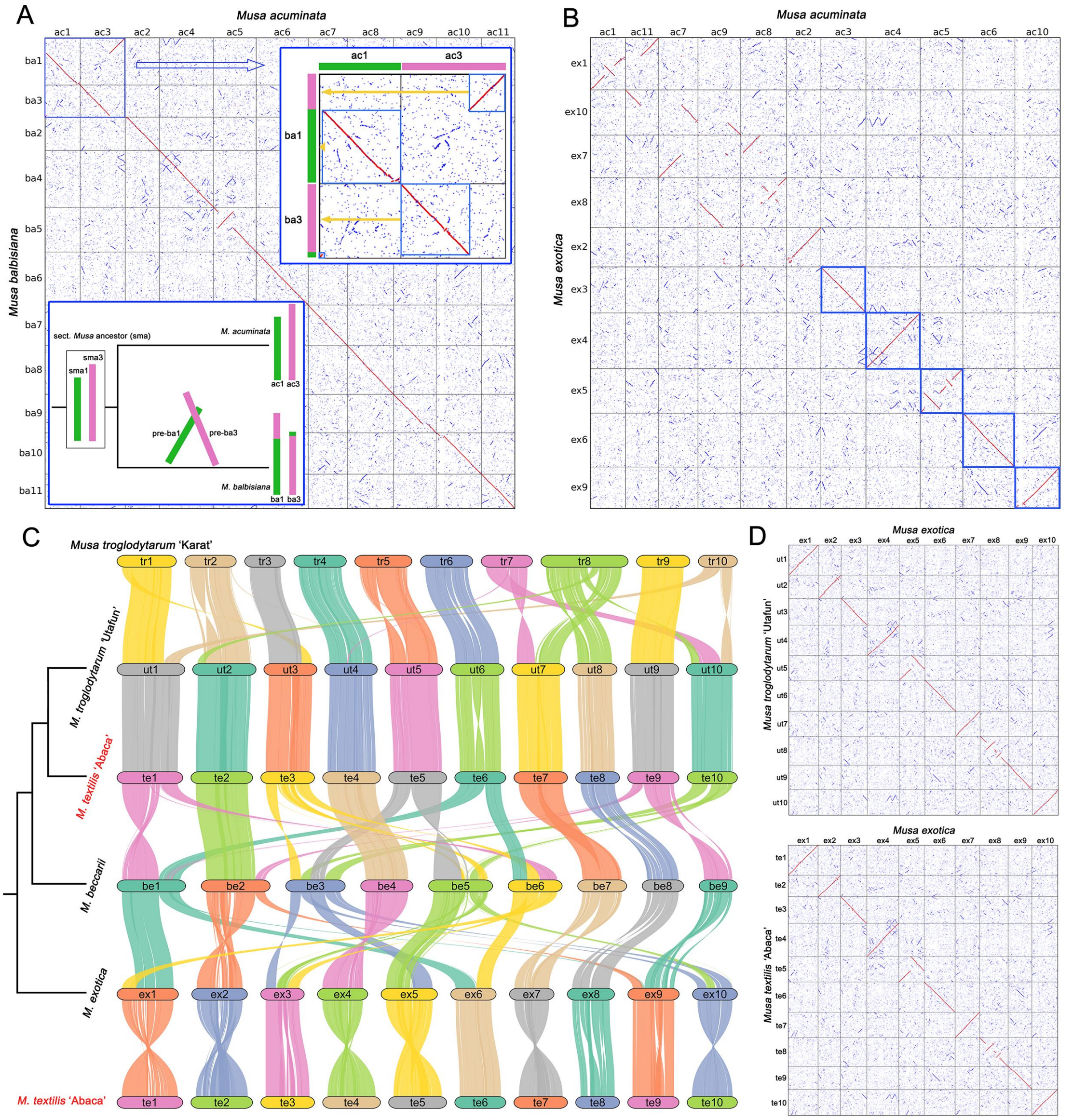


**Figure S12. Genomic collinearity of *Musa* species.** (A) The genomic collinearity dot plot and karyotype variation between *M. acuminata* (Mac) and *M. balbisiana* (Mba). (B) The genomic collinearity dot plot shows the five protochromosomes (blue box) of *Musa*. (C) Genomic collinearity of sect. *Callimusa* species. (D) The genomic collinearity between *M. exotica* (Mex) and *M. textilis* and *M. troglodytarum*. The complete collinearity between Mex's Chr3 and Mac's Chr3 (B) suggests that the reciprocal chromosome translocation occurred independently in Mba. *M. troglodytarum* 'Utafun' and *M. textilis* 'Abaca' both show one-to-one chromosome-level collinearity with Mex (C, D), indicating that the ancestral *Callimusa* karyotype is $n$ = 10. *M. beccarii* is a unique species with $n$ = 9 in *Callimusa*, however, we cannot parse its trajectory of karyotype evolution and believe that its assembled genome need correction.

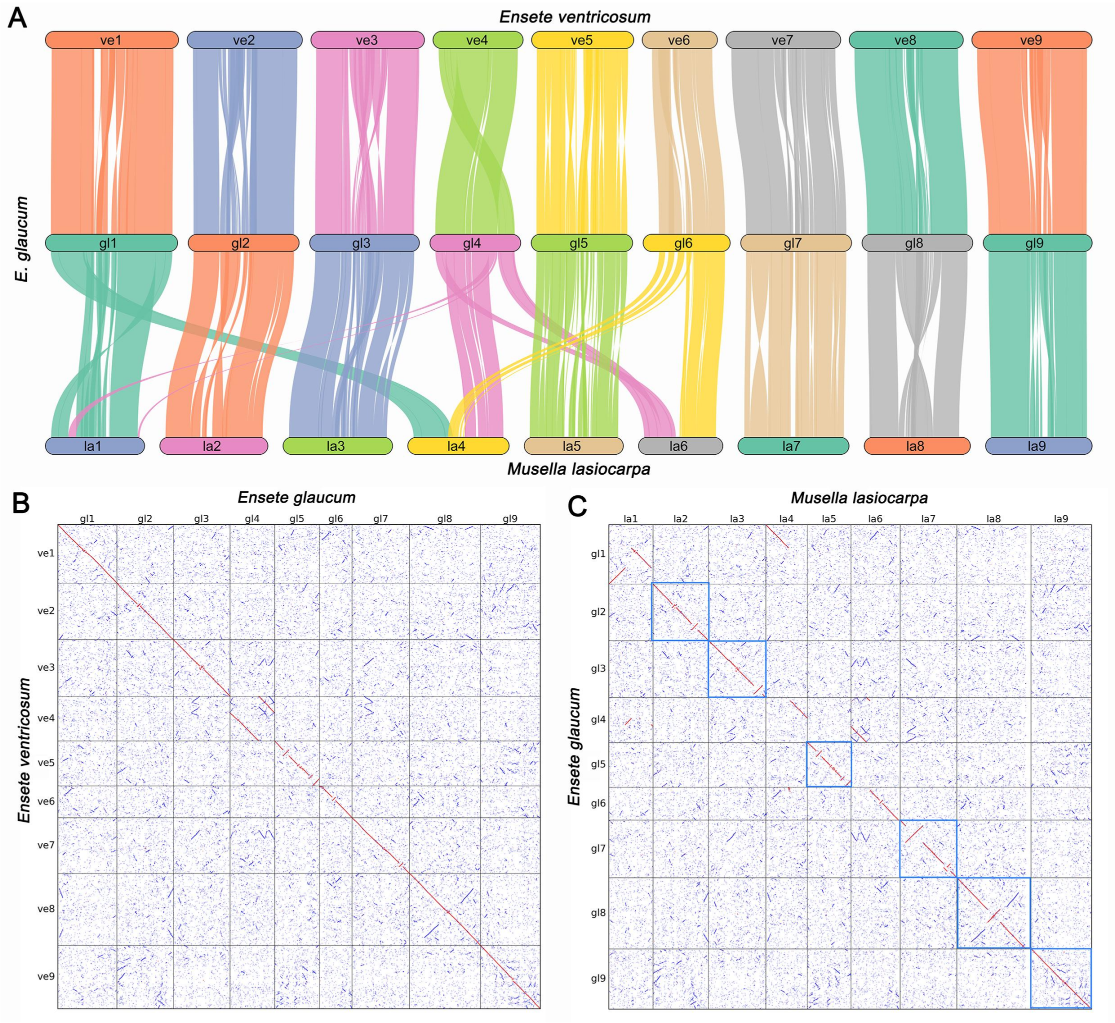


**Figure S13. Genomic collinearity between *Ensete* and *Musella* species.** (A) Genomic collinearity of *Ensete*-*Musella* lineage. (B) Genomic collinearity dot plot between two *Ensete* species. (C) Genomic collinearity dot plot between *Musella lasiocarpa* and *E. glaucum* show the six protochromosomes (blue box) of *Ensete*-*Musella* lineage.

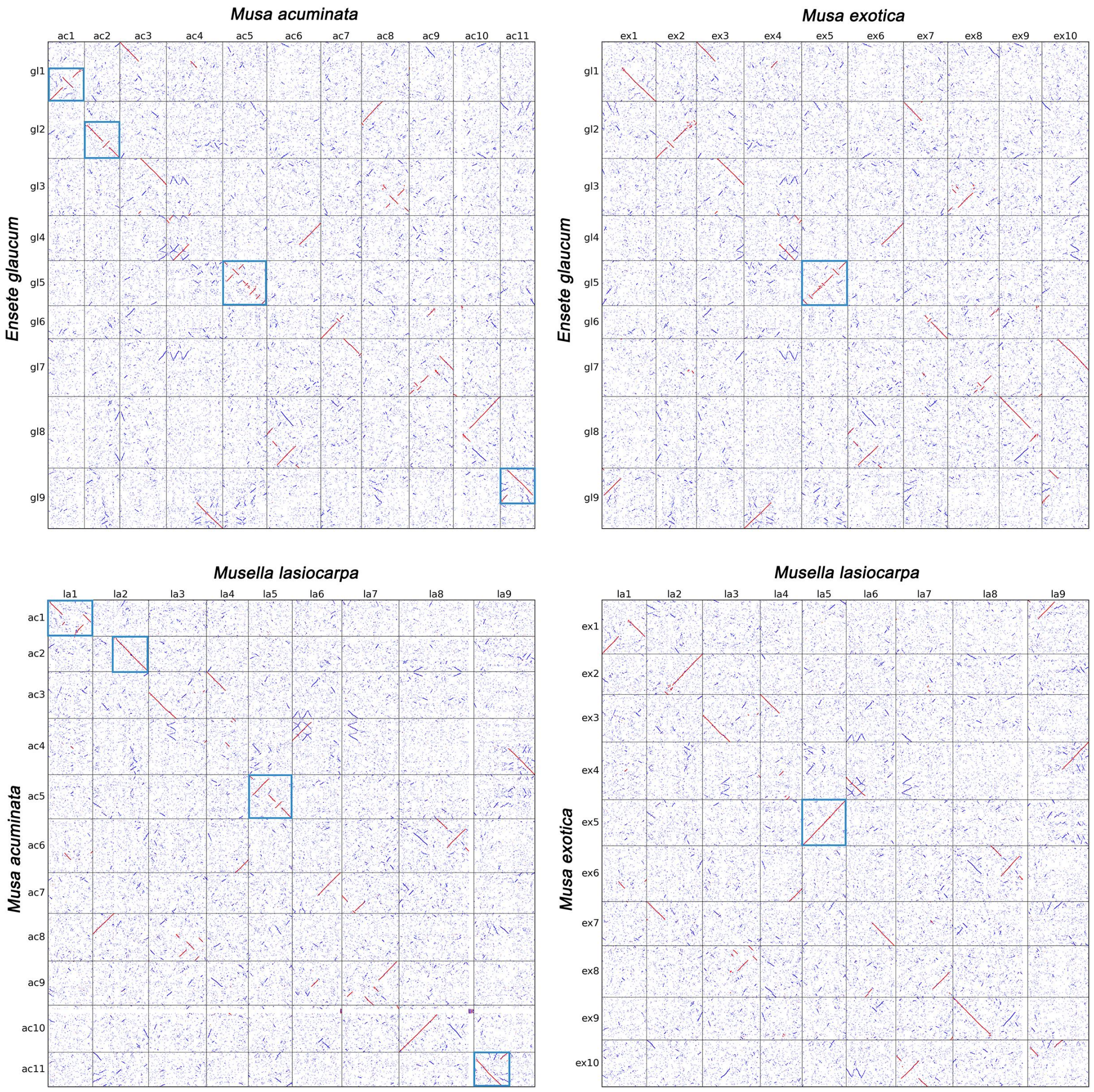


**Figure S14. Genomic collinearity dot plots between different Musaceae species show the protochromosomes (blue box) of Musaceae family.**

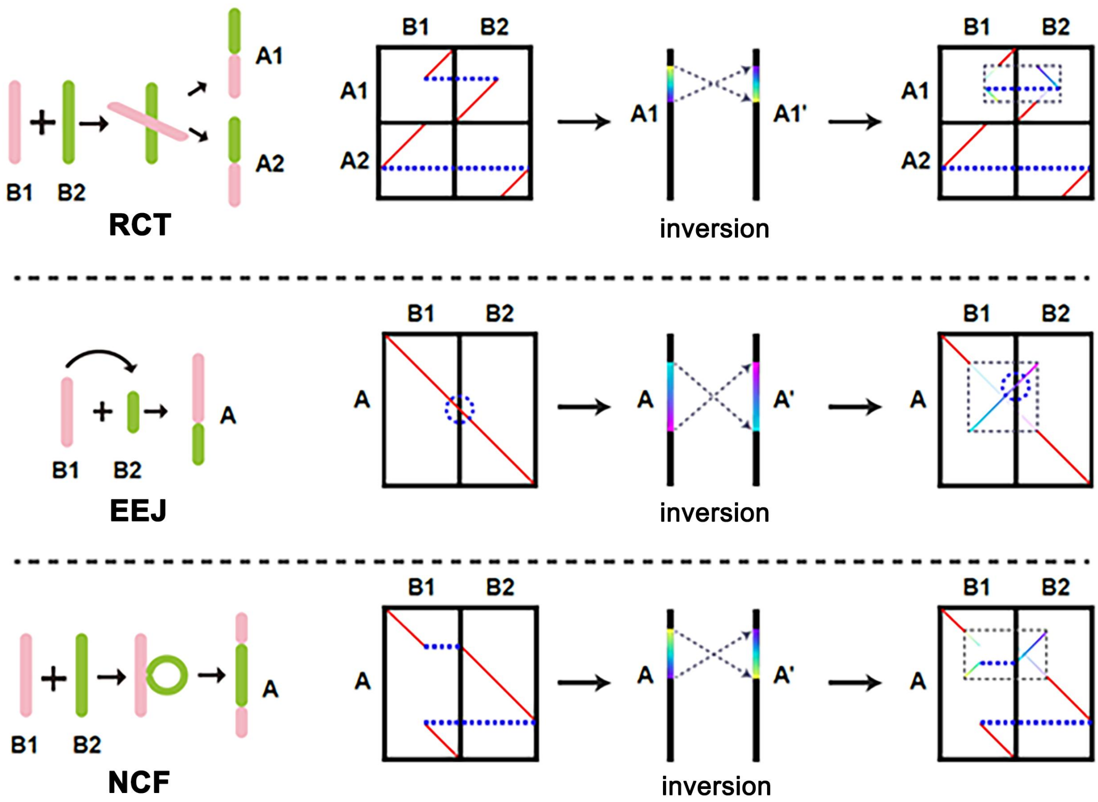


**Figure S15. Three basic types of inter-chromosome rearrangement.** RCT, reciprocal chromosome translocation; EEJ, end-to-end joining; NCF, nested chromosome fusion (https://github.com/SunPengChuan/wgdi-example/blob/main/Karyo-type_Evolution.md).

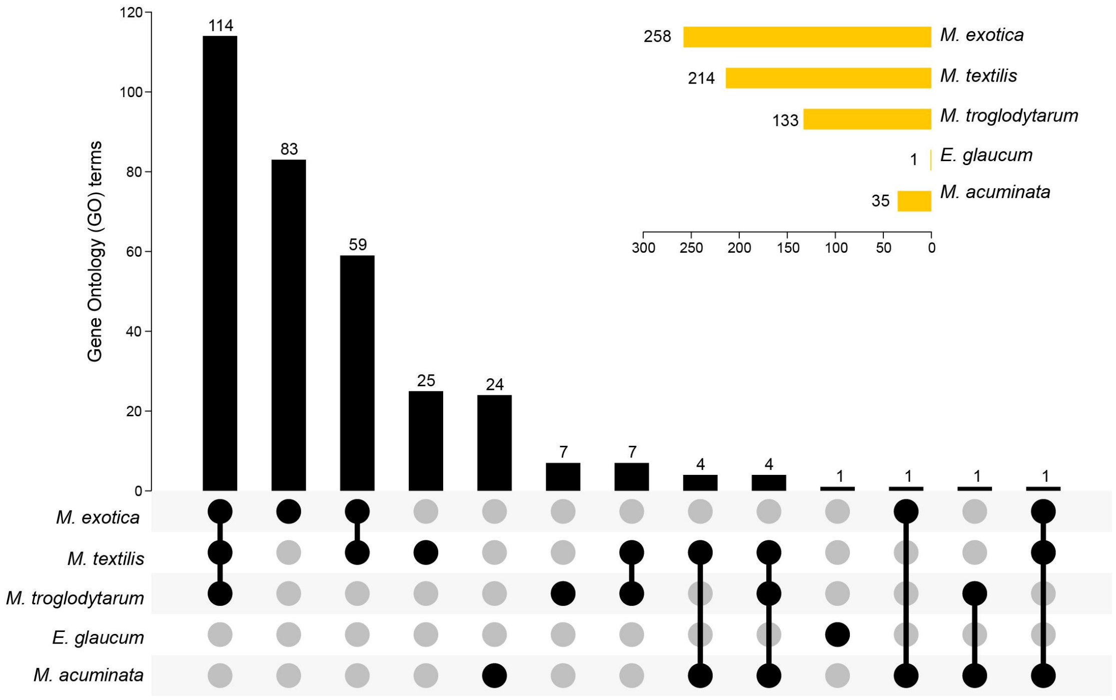


**Figure S16. Functional divergence of genes in chromosomal breakpoint regions across Musaceae revealed by GO enrichment analysis.** Upset plot showing the distribution and overlap of significantly enriched Gene Ontology (GO) terms associated with genes located within and adjacent to chromosomal structural variation breakpoints (±50 flanking genes) across Musaceae species. The bar plot (top) indicates the number of GO terms shared among different species combinations, while the connected dot matrix (bottom) represents the corresponding species intersections. The horizontal bar chart (upper right) summarizes the total number of significantly enriched GO terms per species. Visualization was performed in TBtools-II.

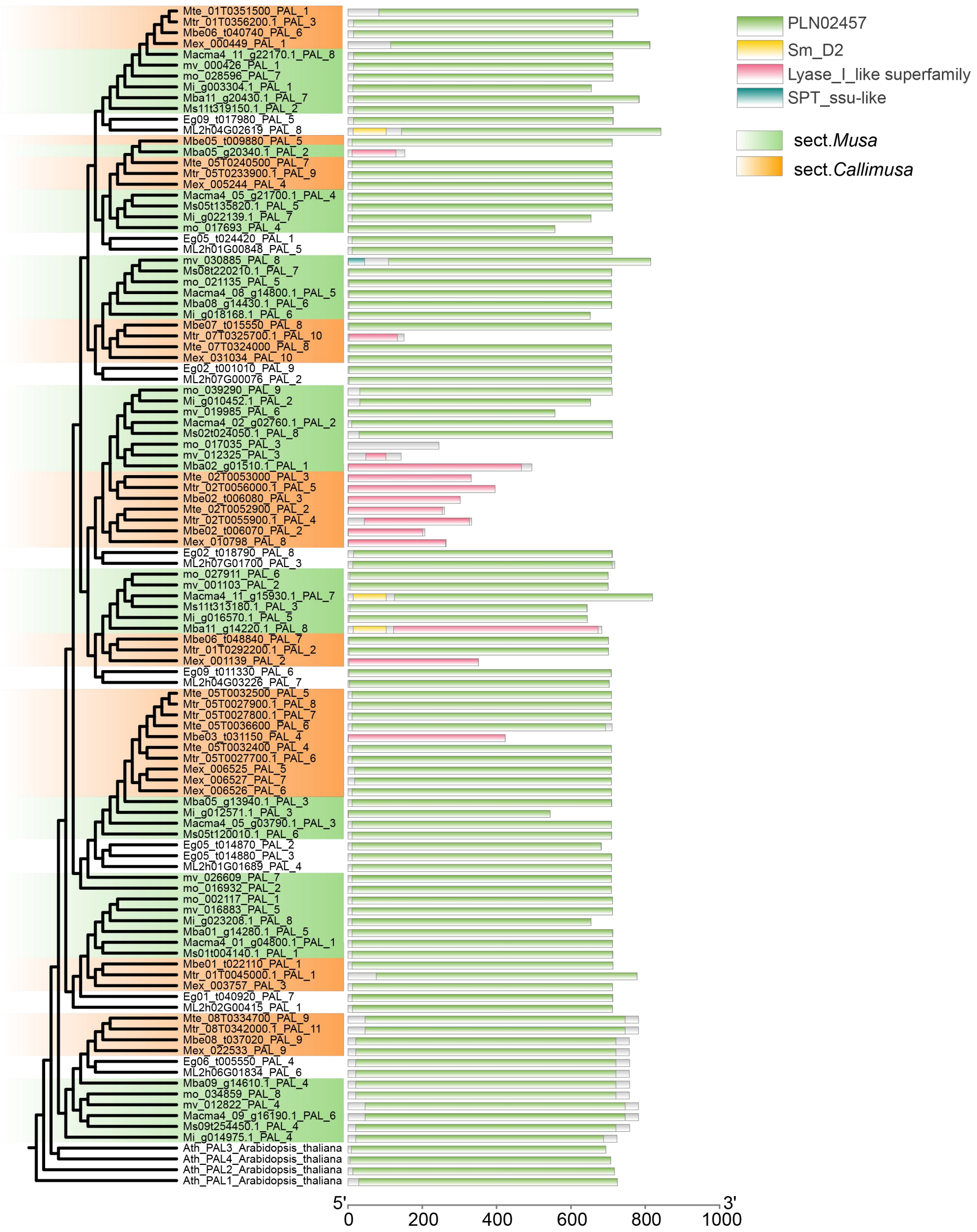


**Figure S17A. Maximum likelihood phylogenetic tree of *PAL/PTAL* gene copies involved in the anthocyanin biosynthesis pathway across Musaceae species and representative outgroups.** Clades are shaded by sections, with green indicating sect. *Musa* and orange indicating sect. *Callimusa*. Conserved domain architectures, annotated using the NCBI Conserved Domain Database (CDD; https://www.ncbi.nlm.nih.gov/Structure/bwrpsb/bwrpsb.cgi), are displayed as bar plots to the right of the tree. Visualization was performed in TBtools-II.

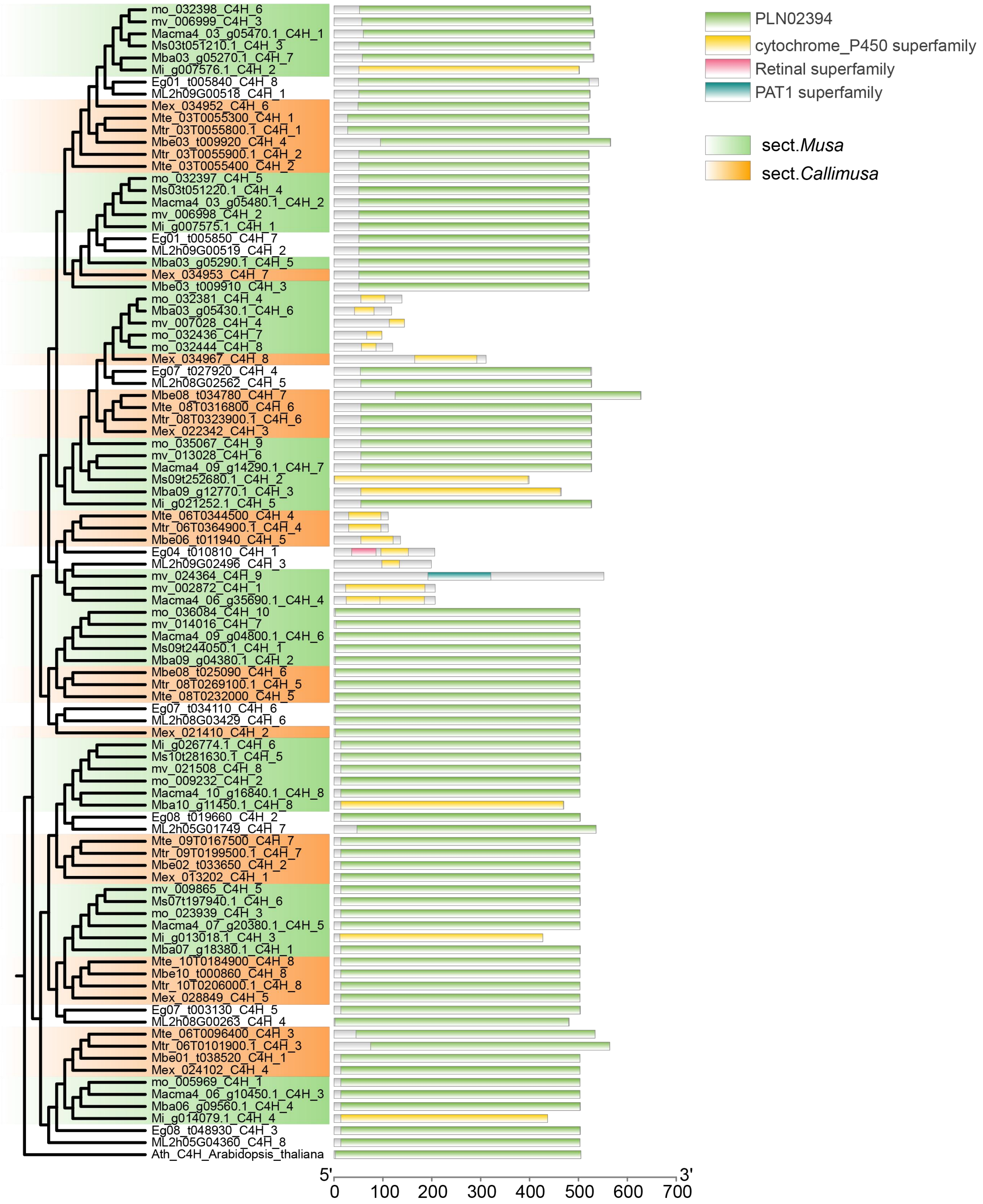


**Figure S17B. Maximum likelihood phylogenetic tree of *C4H* gene copies involved in the anthocyanin biosynthesis pathway across Musaceae species and representative outgroups.**

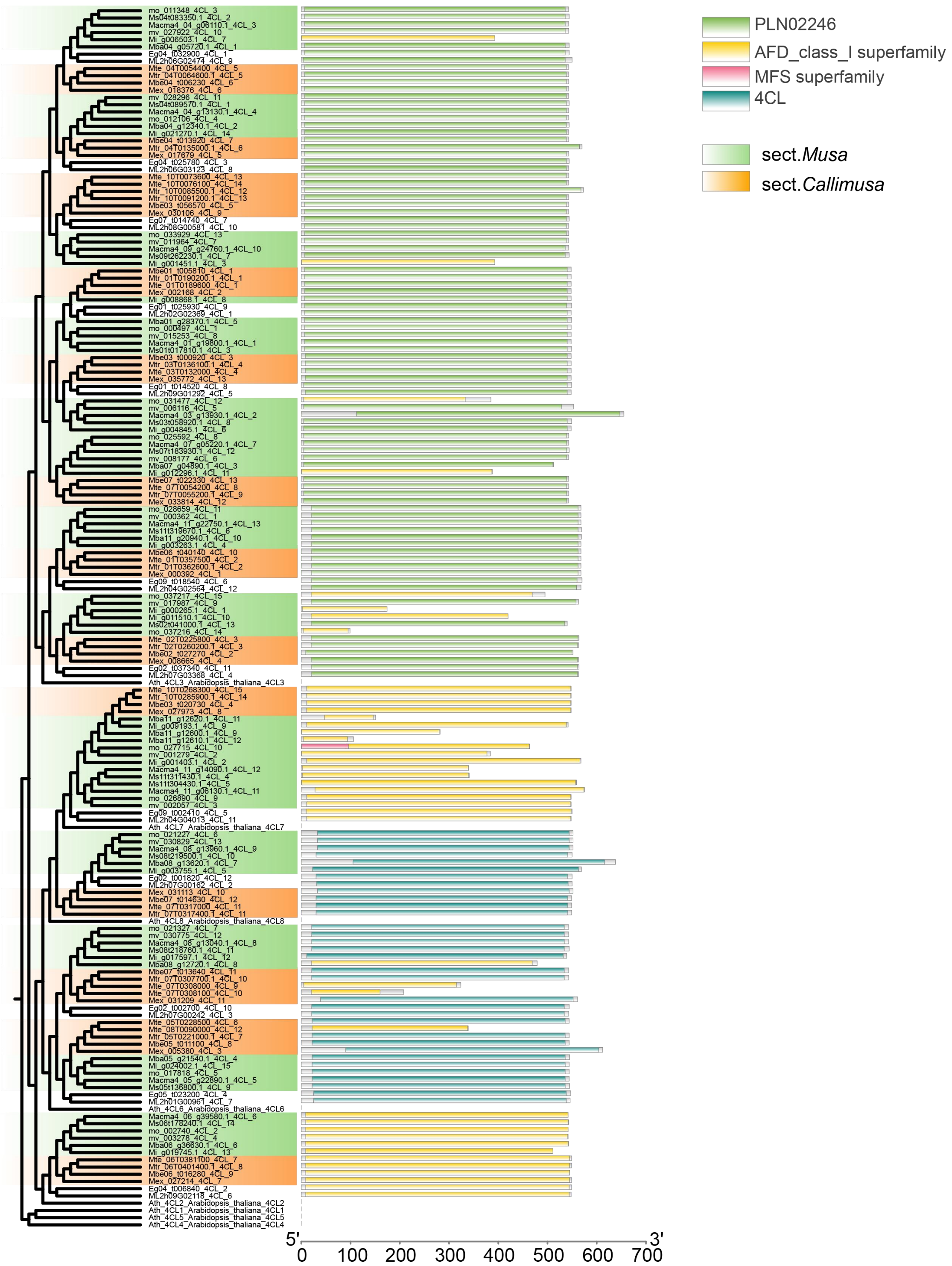


**Figure S17C. Maximum likelihood phylogenetic tree of *4CL* gene copies involved in the anthocyanin biosynthesis pathway across Musaceae species and representative outgroups.**

### Part I

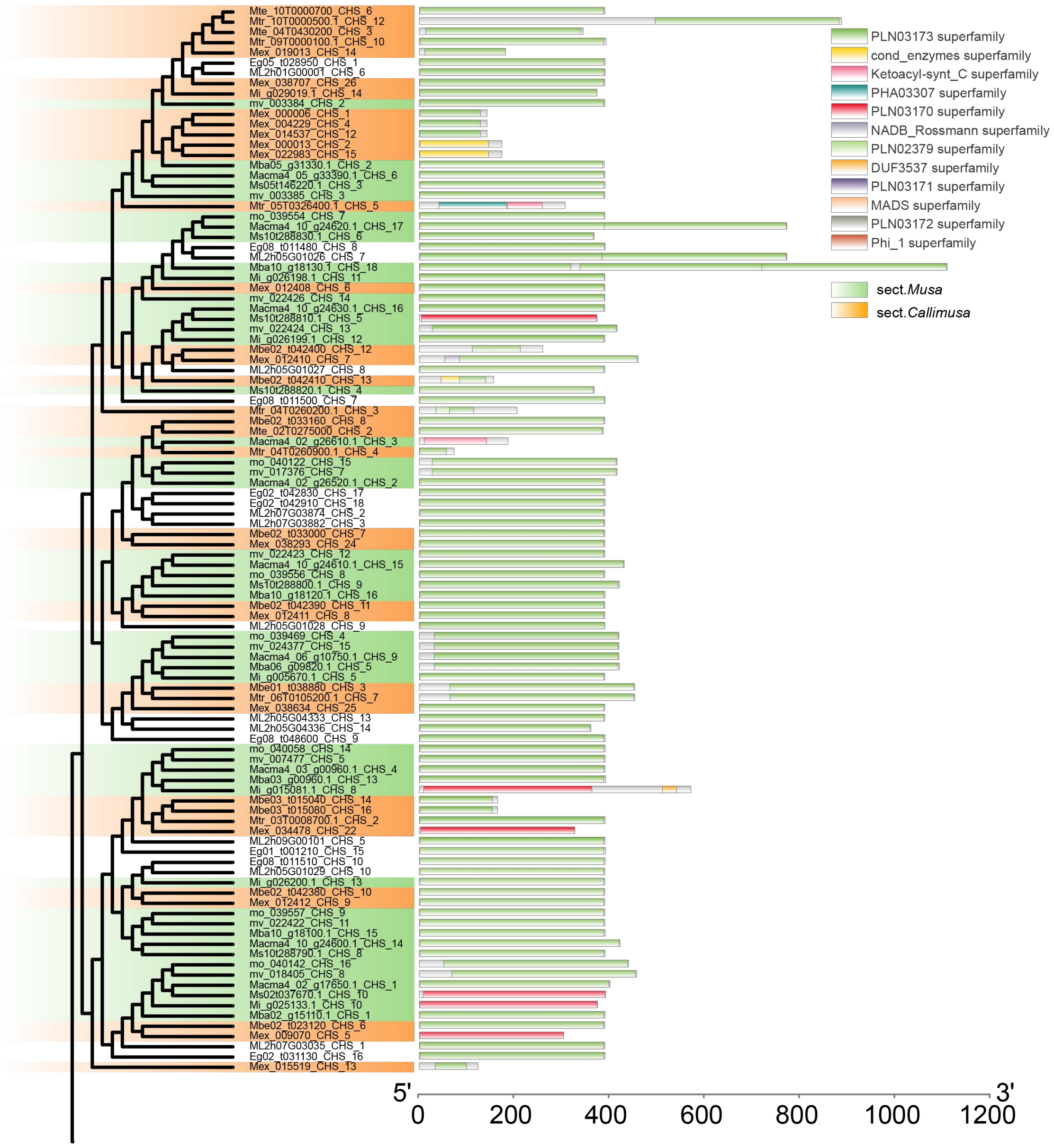


**Figure S17D. Maximum likelihood phylogenetic tree of *CHS* (PART I) gene copies involved in the anthocyanin biosynthesis pathway across Musaceae species and representative outgroups.**

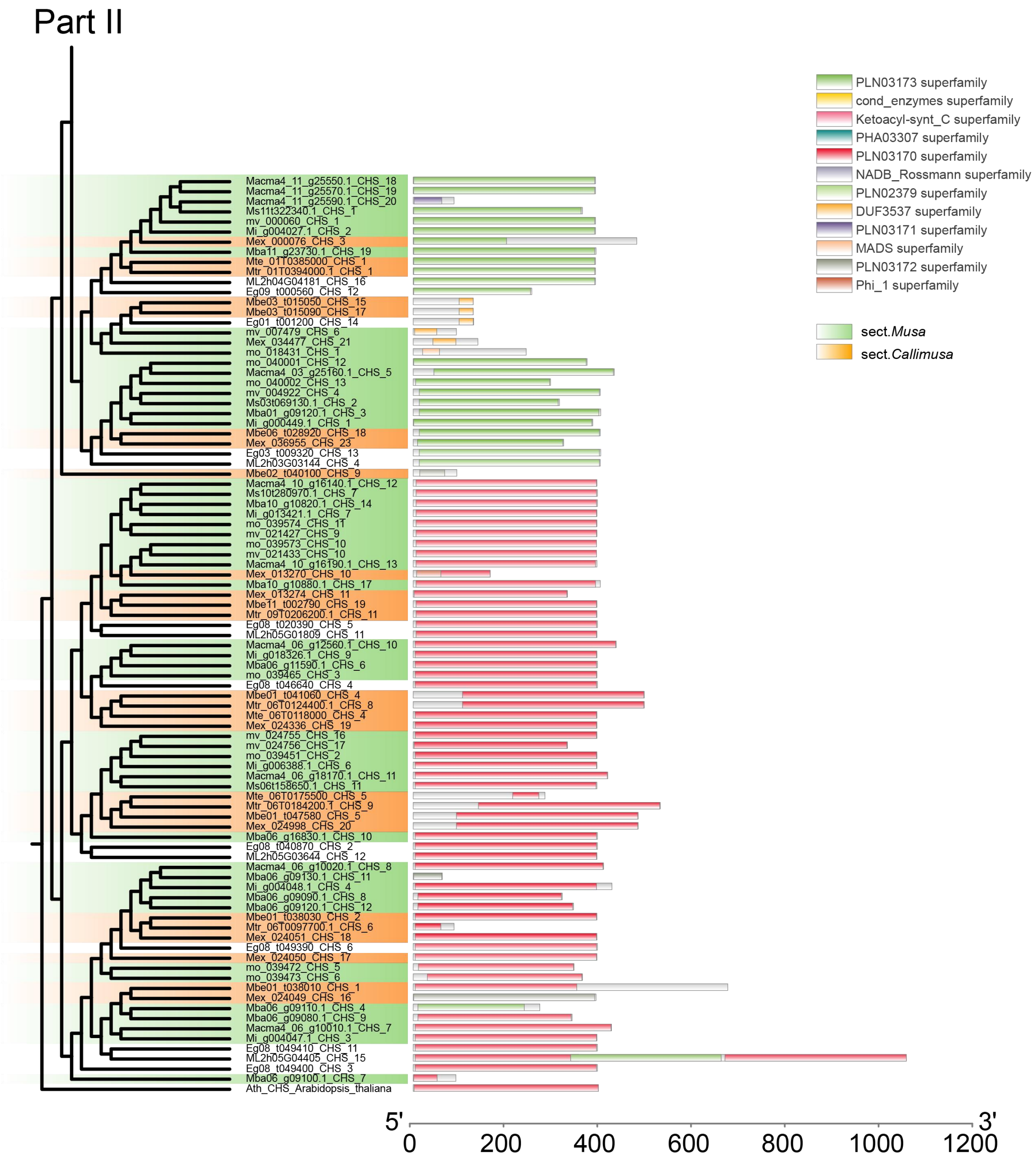


**Figure S17E. Maximum likelihood phylogenetic tree of *CHS* (PART II) gene copies involved in the anthocyanin biosynthesis pathway across Musaceae species and representative outgroups.**

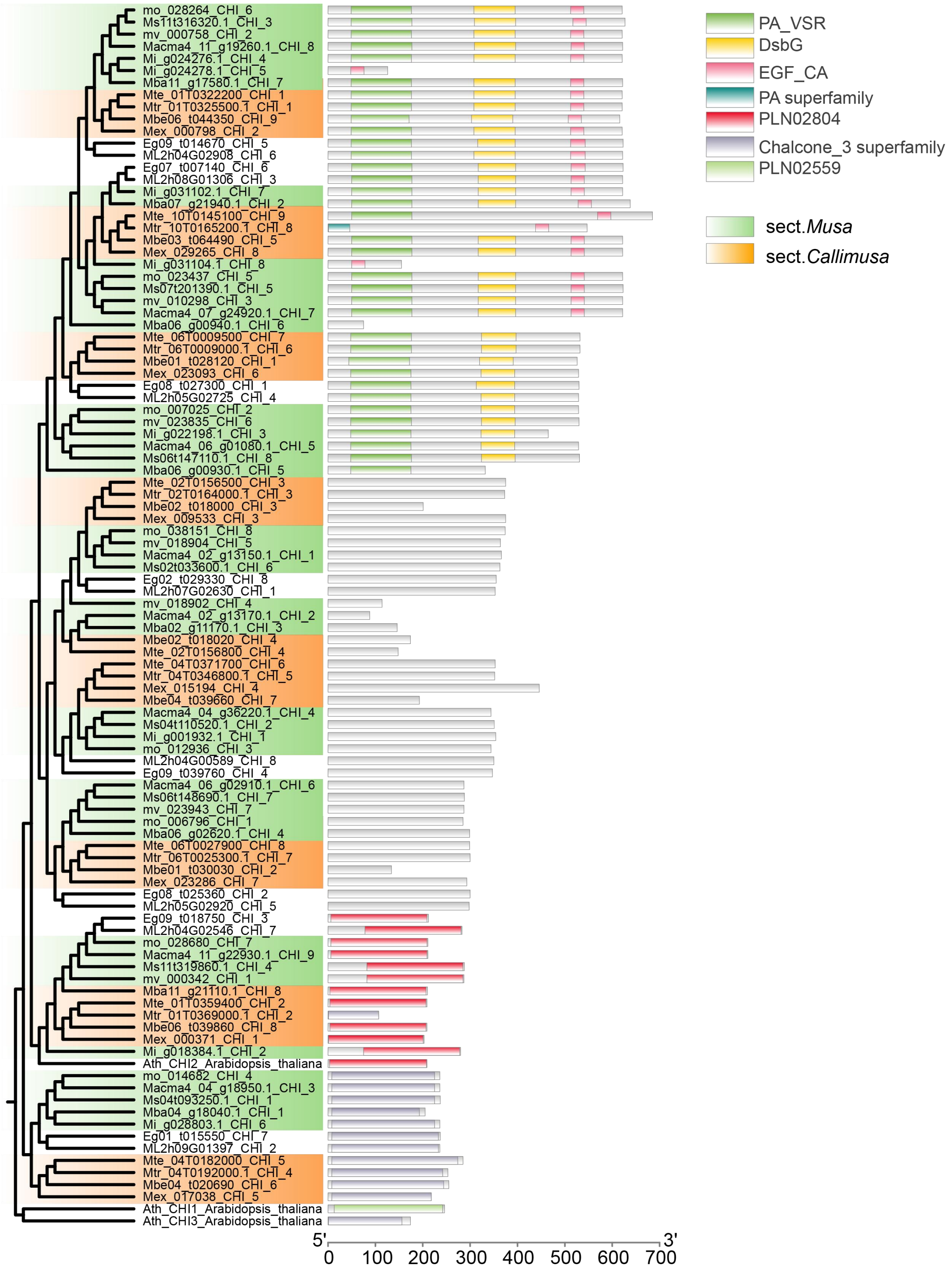


**Figure S17F. Maximum likelihood phylogenetic tree of *CHI* gene copies involved in the anthocyanin biosynthesis pathway across Musaceae species and representative outgroups.**

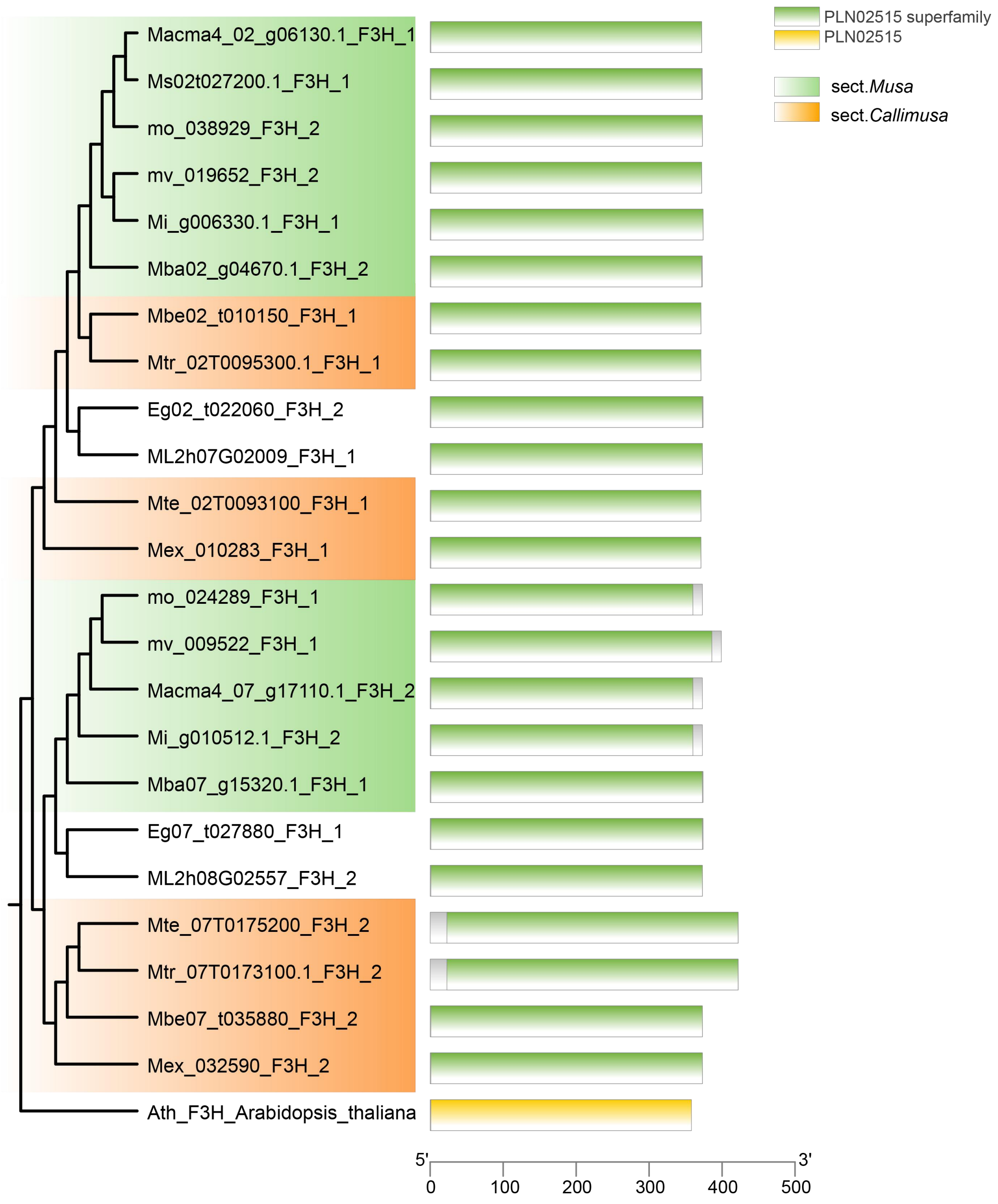


**Figure S17G. Maximum likelihood phylogenetic tree of *F3H* gene copies involved in the anthocyanin biosynthesis pathway across Musaceae species and representative outgroups.**

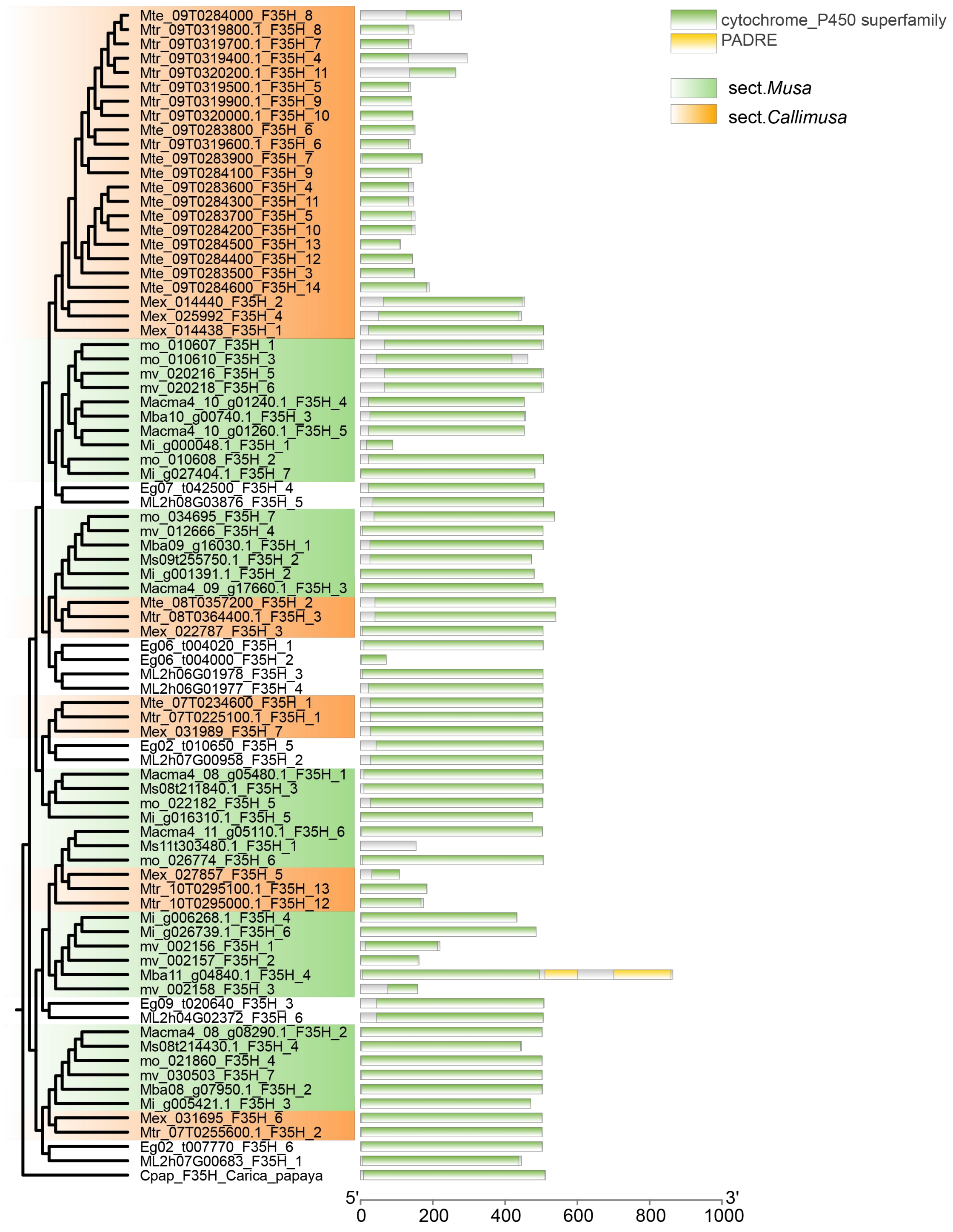


**Figure S17H. Maximum likelihood phylogenetic tree of *F3′5′H* gene copies involved in the anthocyanin biosynthesis pathway across Musaceae species and representative outgroups.**

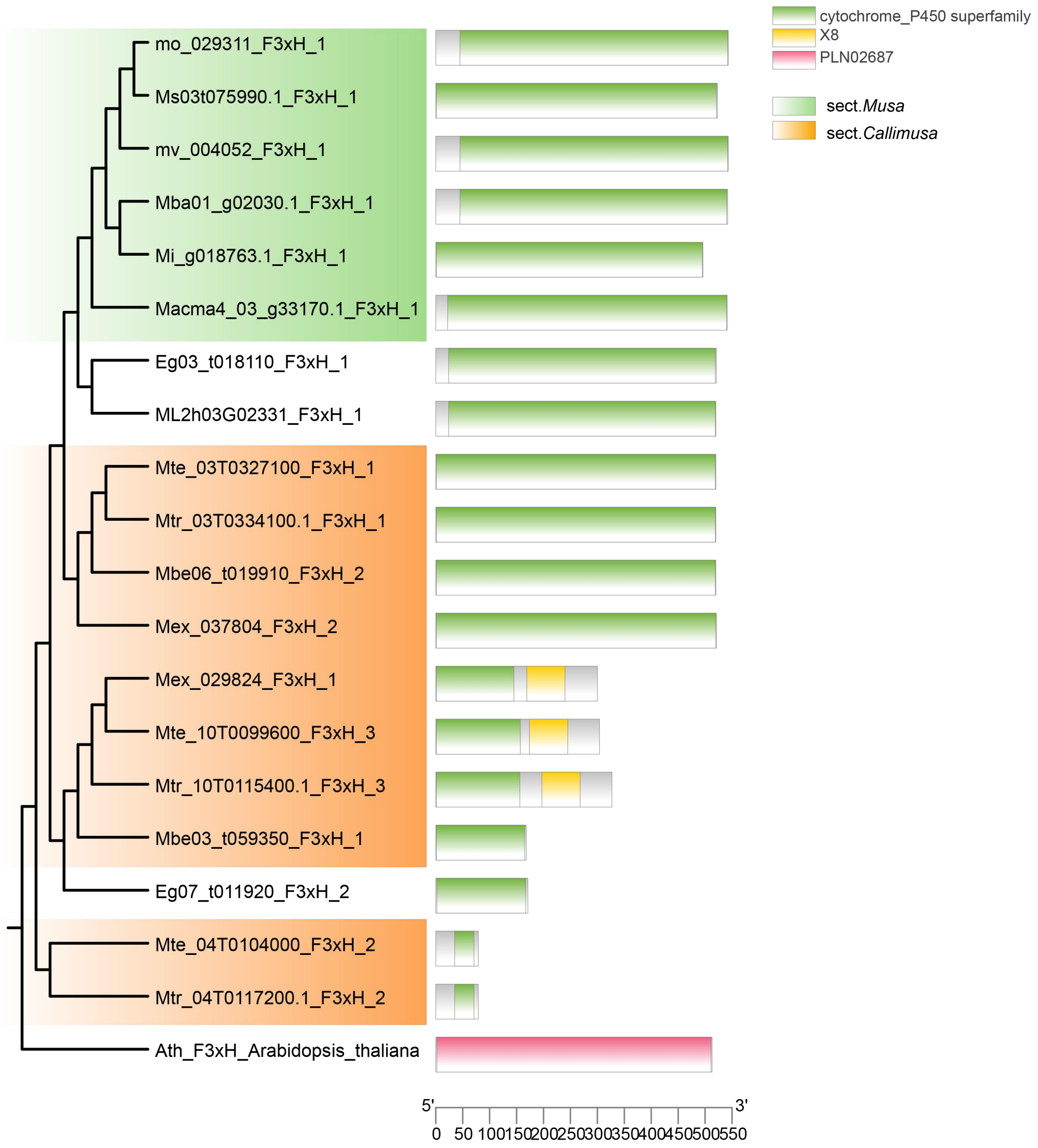


**Figure S17I. Maximum likelihood phylogenetic tree of *F3'H* gene copies involved in the anthocyanin biosynthesis pathway across Musaceae species and representative outgroups.**

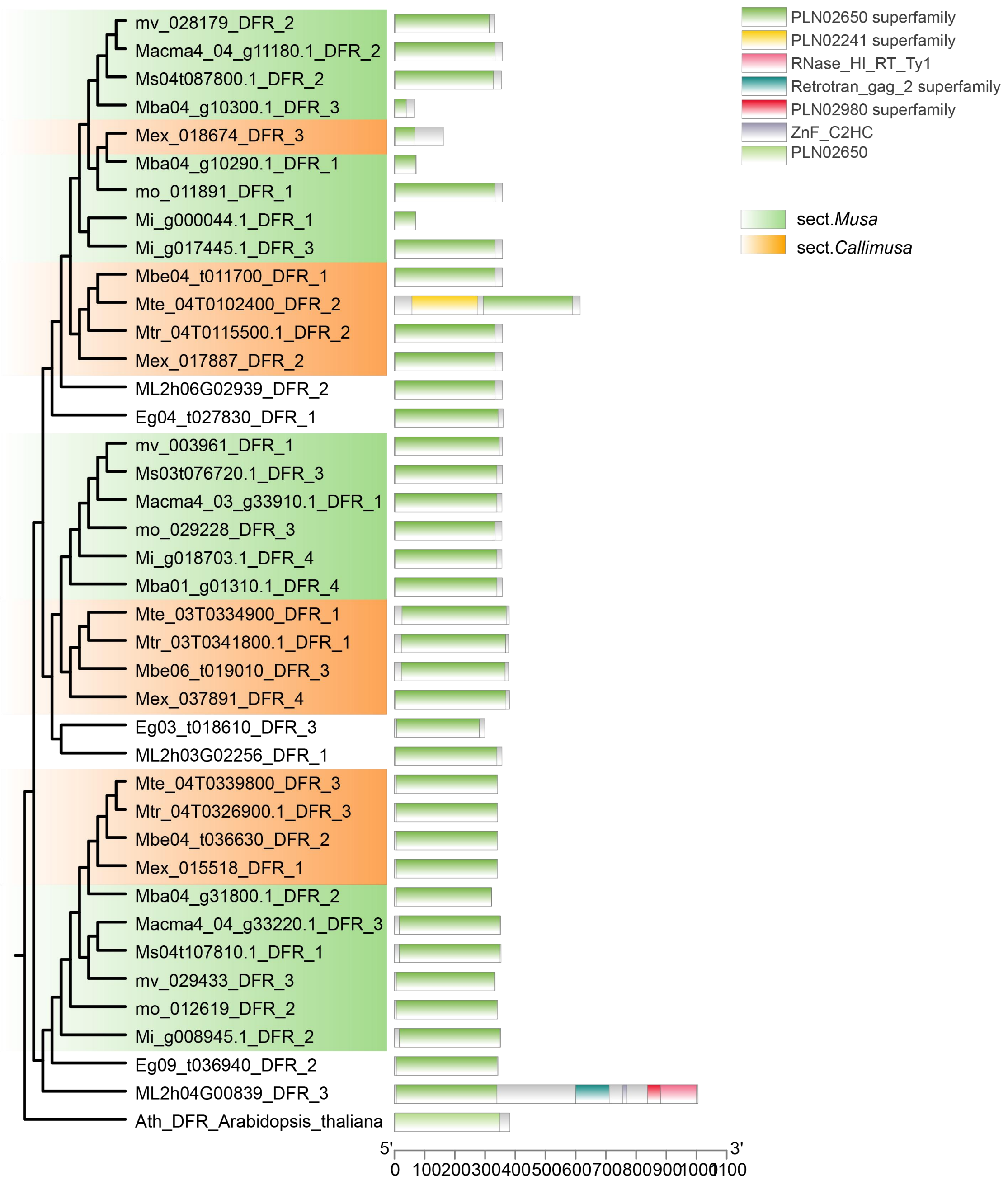


**Figure S17J. Maximum likelihood phylogenetic tree of *DFR* gene copies involved in the anthocyanin biosynthesis pathway across Musaceae species and representative outgroups.**

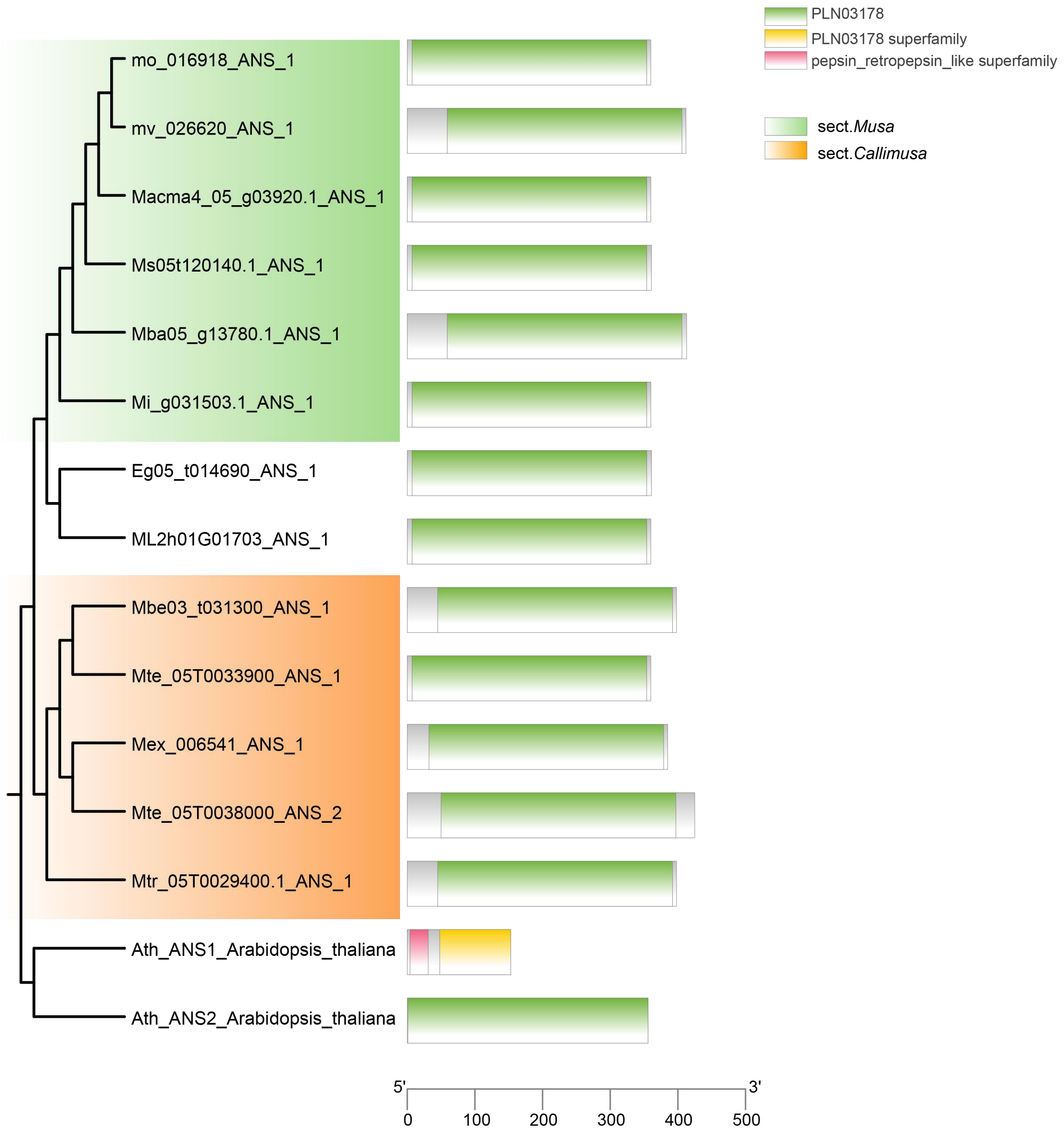


**Figure S17K. Maximum likelihood phylogenetic tree of *ANS* gene copies involved in the anthocyanin biosynthesis pathway across Musaceae species and representative outgroups.**

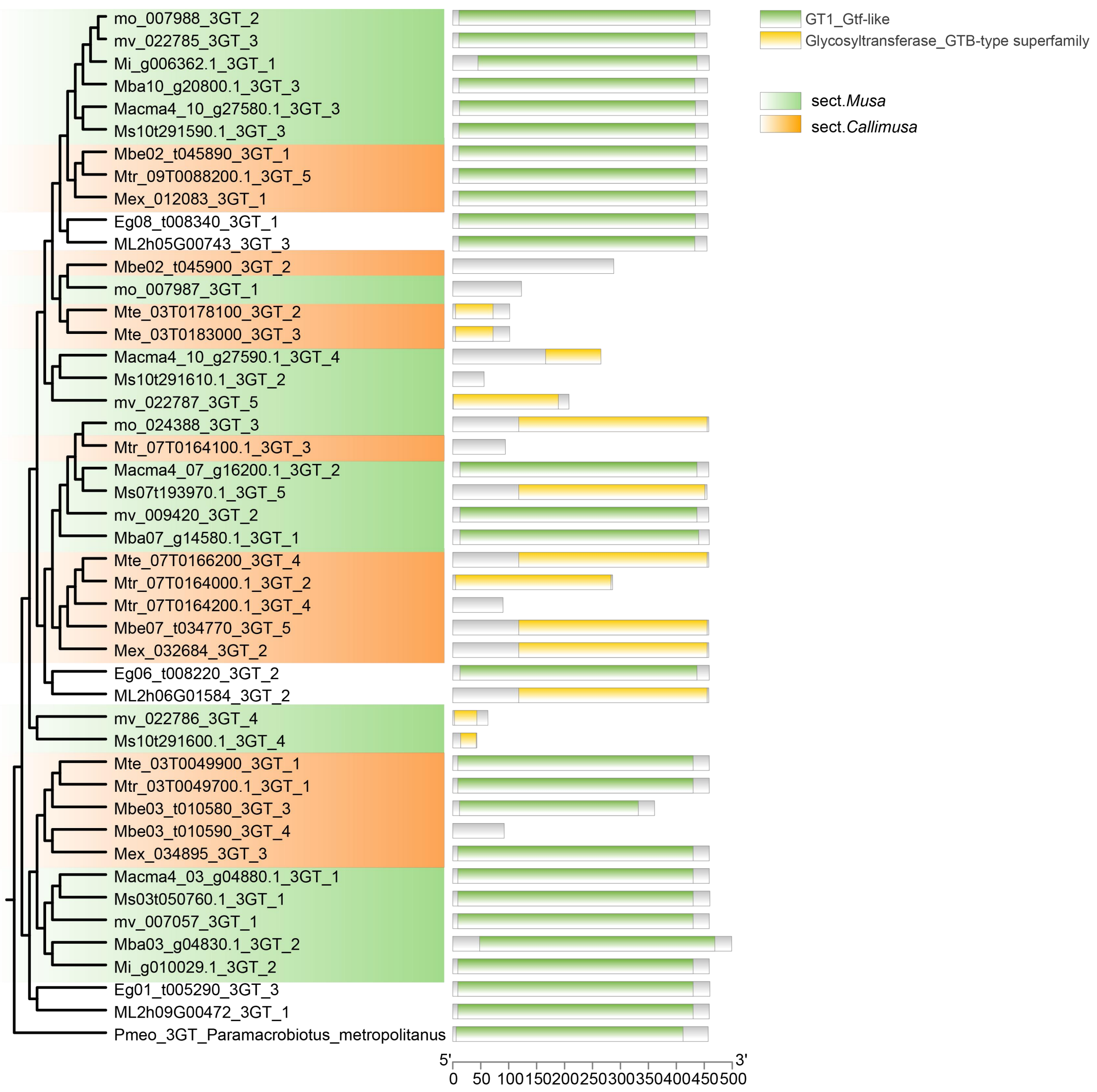


**Figure S17L. Maximum likelihood phylogenetic tree of *3-GT* gene copies involved in the anthocyanin biosynthesis pathway across Musaceae species and representative outgroups.**

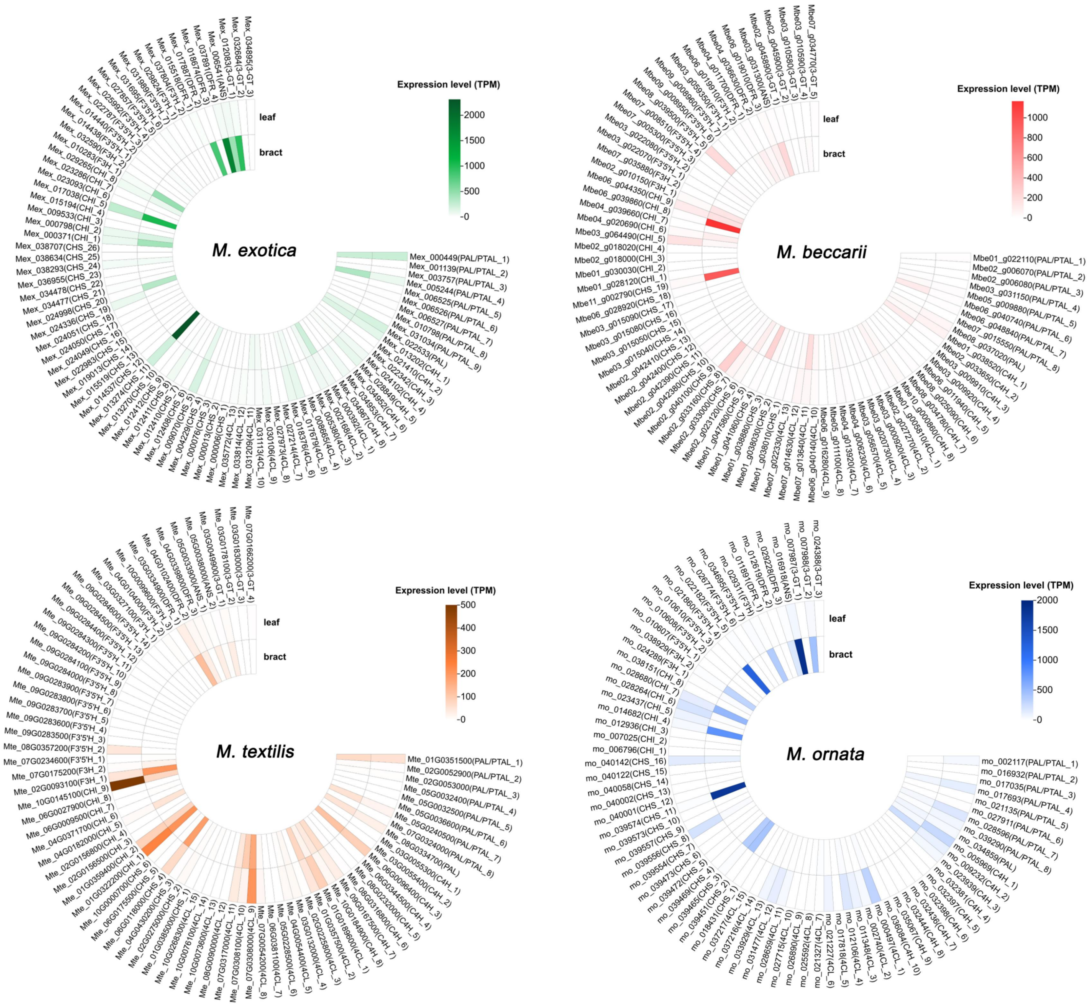


**Figure S18. Expression profiles of enzyme-coding genes in bract and leaf tissues of sect. *Callimusa* species.** Gene expression levels were quantified in transcripts per million (TPM) and standardized across samples to enable comparison between tissues.

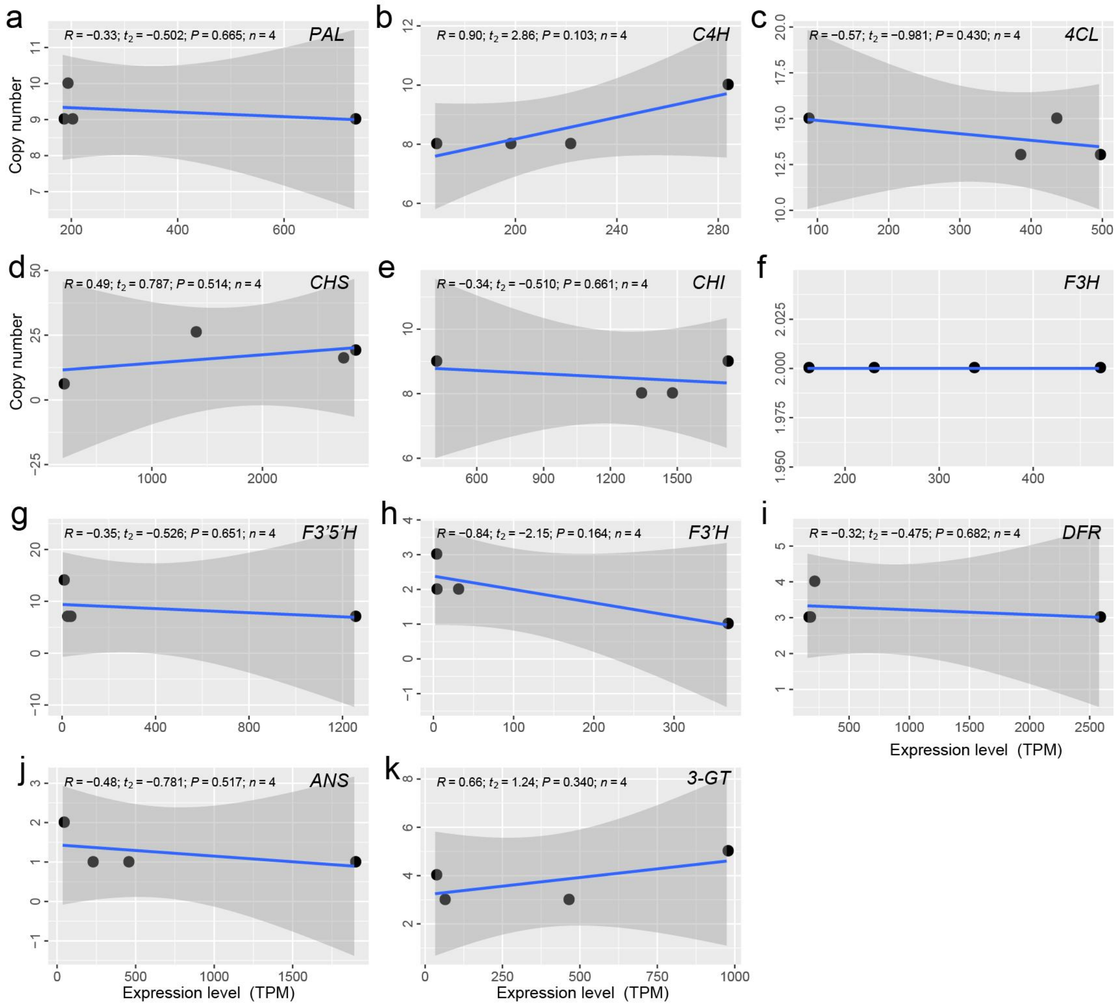


**Figure S19. Correlation between gene copy number and expression levels of key anthocyanin biosynthesis genes.** Linear regression lines were fitted using an ordinary least squares (OLS) model, with 95% confidence intervals indicated by shaded regions. Pearson's correlation coefficients (*R*) were used to assess statistical significance. The *R*, *t*, *P* values, and sample sizes (*n*) are provided in each panel, indicating no significant correlation between gene copy number and expression level.
