## Supplementary Data for "Ancestral Musaceae karyotype reconstruction provides insights into chromosome evolution and bract coloration"

### Detailed Procedures for Ancestral Karyotype Reconstruction in Musaceae

The ancestral karyotype can be inferred using either a bottom-up approach (progressively merging from closely related lineages) or a top-down approach (starting from hypotheses about deeper ancestors). Given that chromosome numbers vary little across the major lineages of Musaceae and structural rearrangements within lineages are relatively limited, we adopt the bottom-up workflow here.

#### Step 1: Reconstruct the Ancestral Karyotype for Each Clade Node

##### sect. *Callimusa*

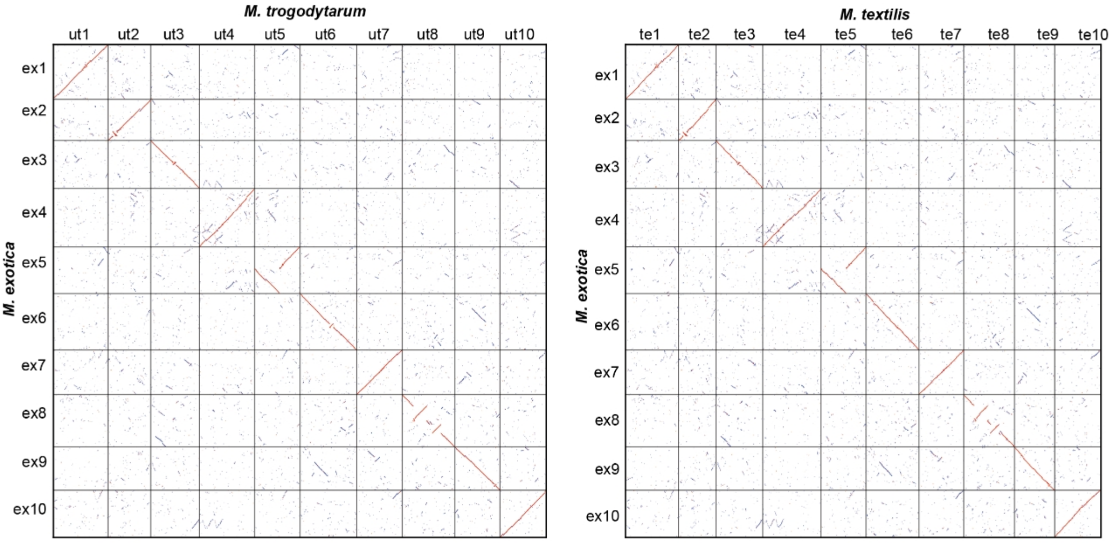

The three genomes show a 1:1 chromosomal correspondence, indicating that all three have largely retained the ancestral karyotype of this clade. Any of these species could therefore serve as a representative of the nodal ancestor; here we select *M. exotica* (Mex).

#### sect. *Musa*

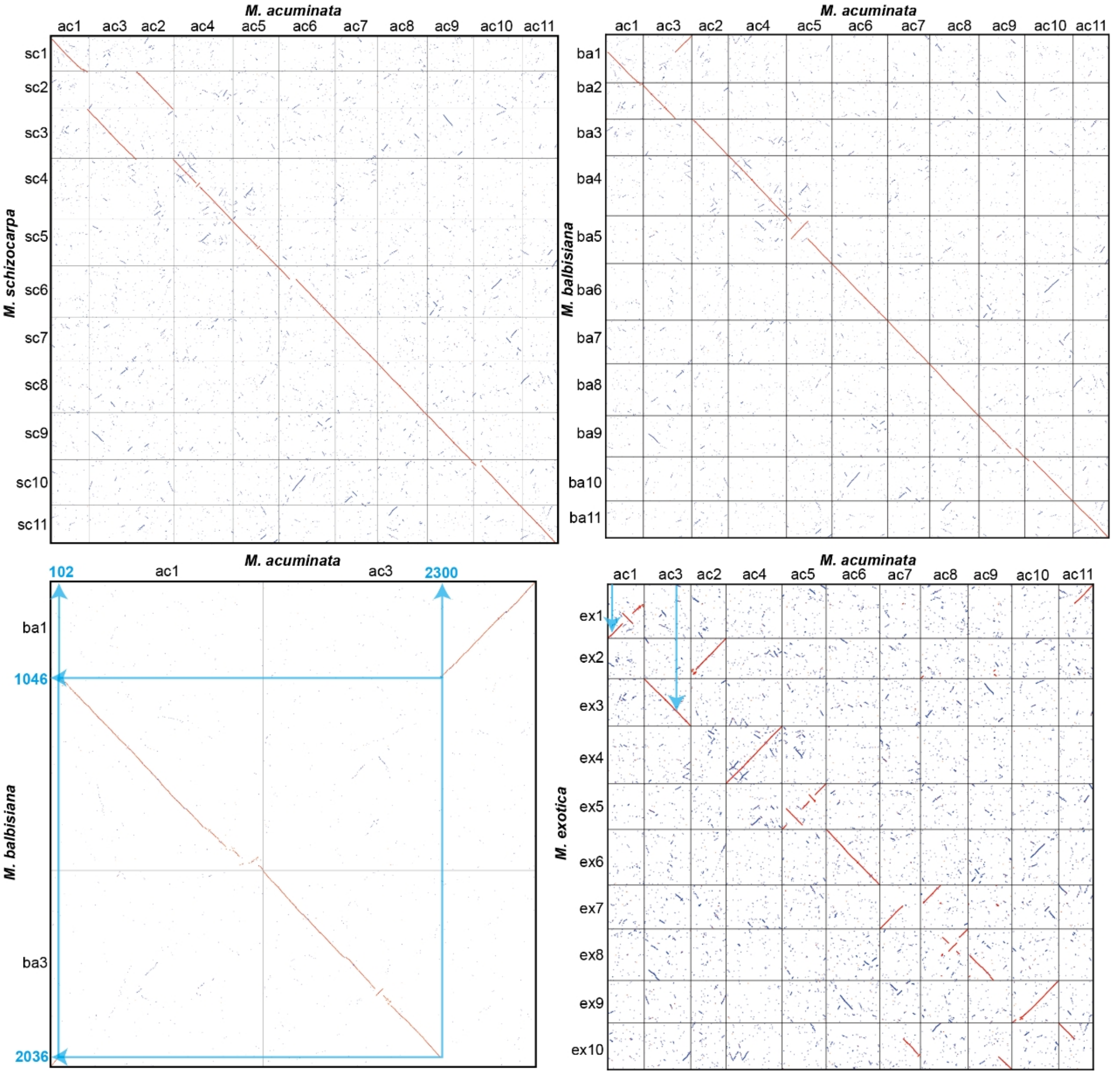

*M. acuminata* (Mac) and *M. schizocarpa* (Msc) share a 1:1 chromosomal relationship, while a reciprocal translocation exists between Mac and *M. balbisiana*. Verification using Mex from the outgroup clade sect. *Callimusa* and the outgroup *Canna indica* consistently shows that Mex and Mac share the same chromosomal structure, indicating that the reciprocal translocation most likely occurred specifically in the *M. balbisiana* lineage. The ancestral karyotype of this clade node is therefore represented by Mac.

*Ensete*

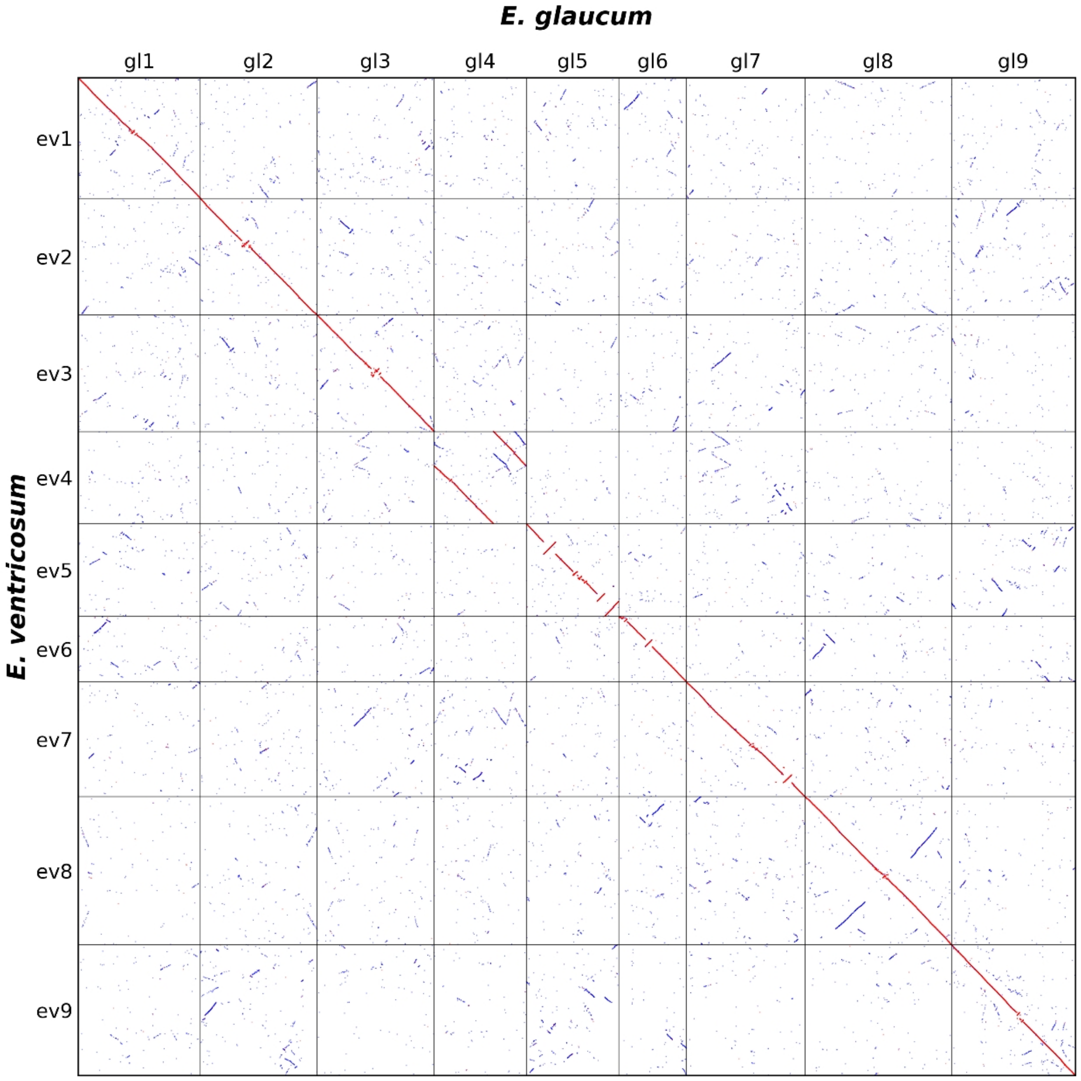

*E. ventricosum* (Eve) and *E. glaucum* (Egl) exhibit a complete 1:1 correspondence with clear collinearity, so Eve is used as the representative of the ancestral karyotype for this clade node.

***Musella***

Only one genome is currently available, *M. lasiocarpa* (Mla), which is therefore used directly to represent the ancestral karyotype of this clade node.

**Step 2: Cross-Clade Comparisons to Identify Protochromosomes**

**sect. *Callimusa* vs. sect. *Musa* (Mex vs. Mac)**

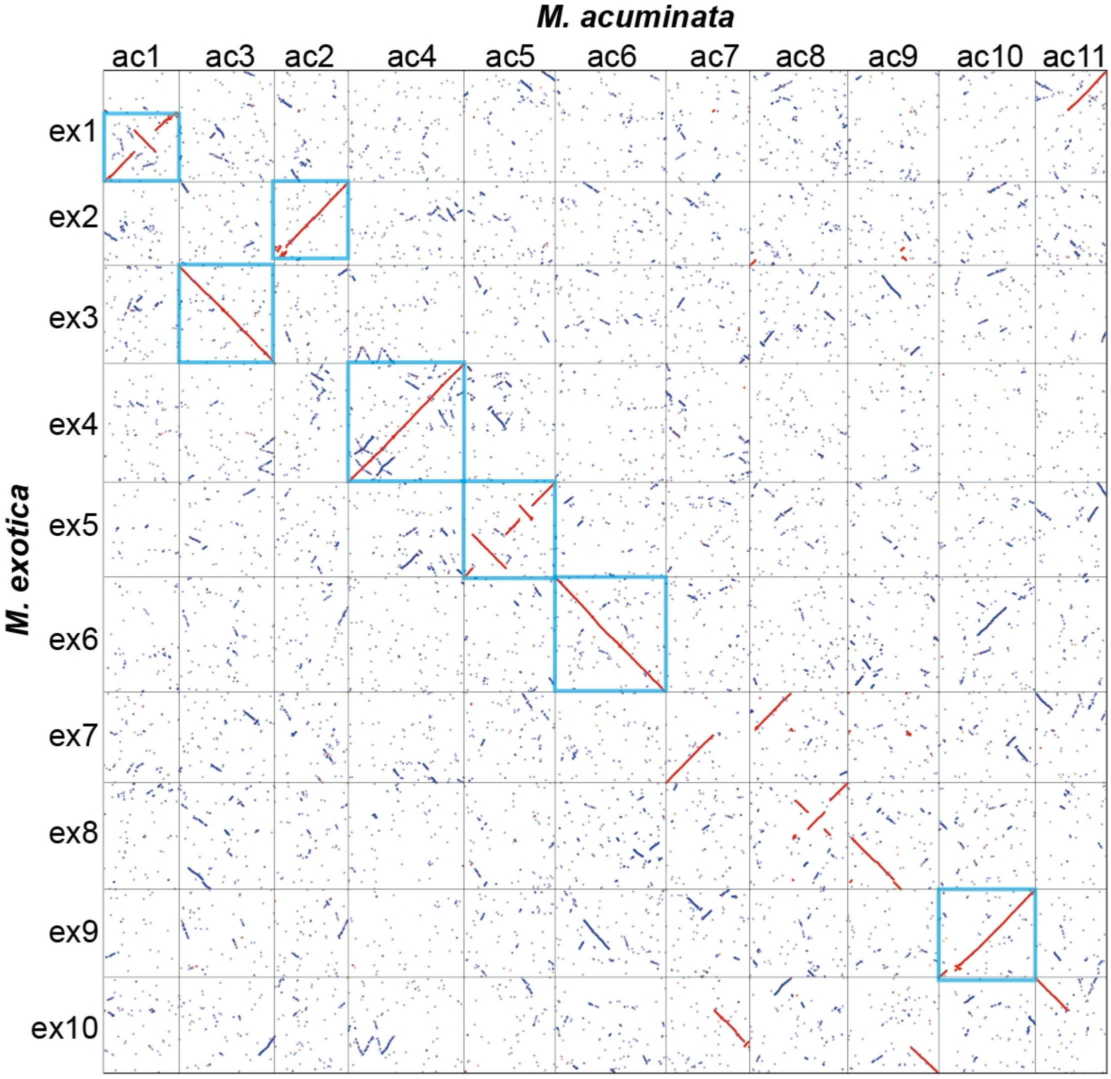

Since the nodal ancestor of sect. *Callimusa* can be substituted by Mex, and that of sect. *Musa* by Mac, a direct comparison between Mex and Mac is sufficient. By identifying Shared or Nested patterns in the dot plot / synteny results, **7 protochromosomes** can be defined (blue boxes).

***Musella* vs. *Ensete* (Mla vs. Ev)**

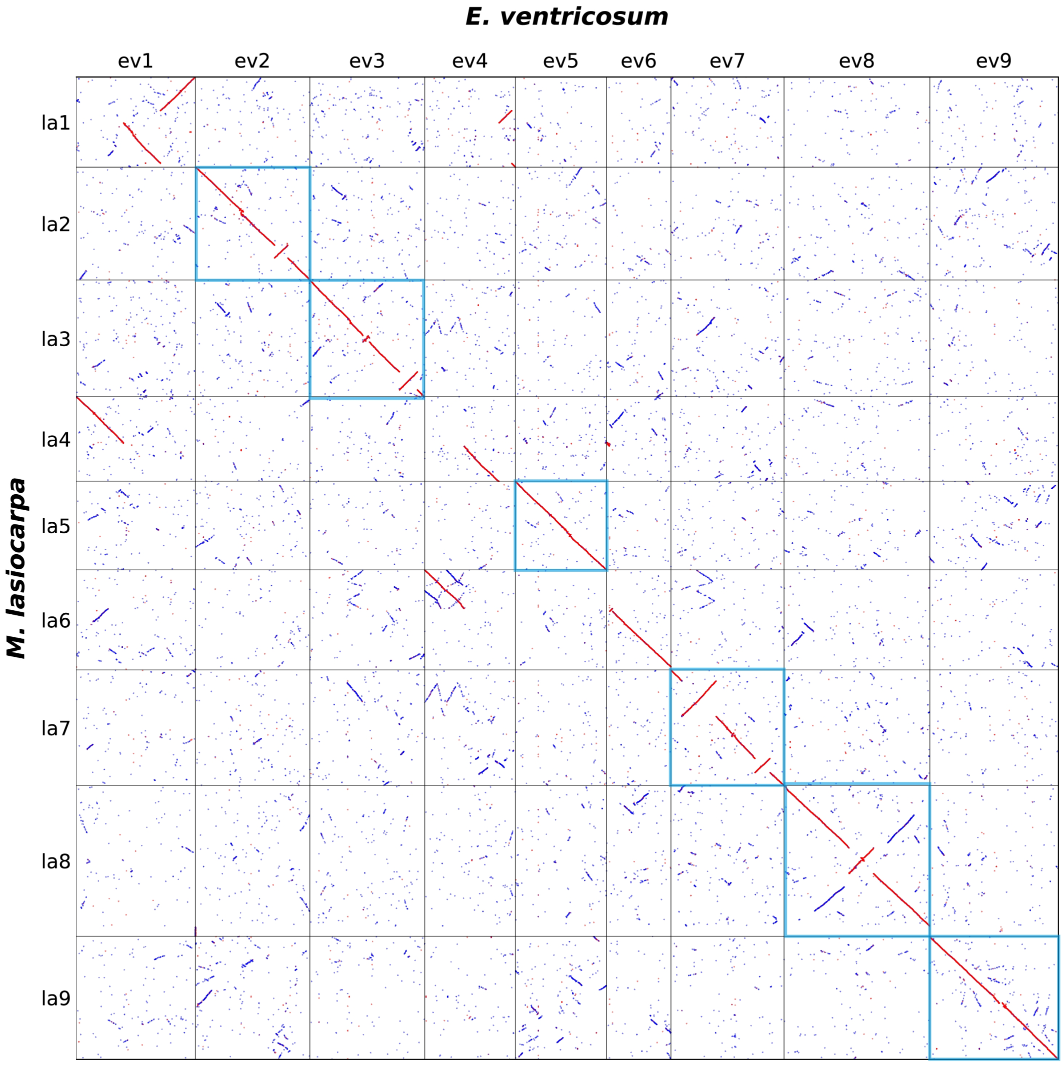

Likewise, by identifying Shared or Nested patterns, **6 protochromosomes** can be defined (blue boxes).

### Inferring the Ancestral Musaceae Karyotype (AMK)

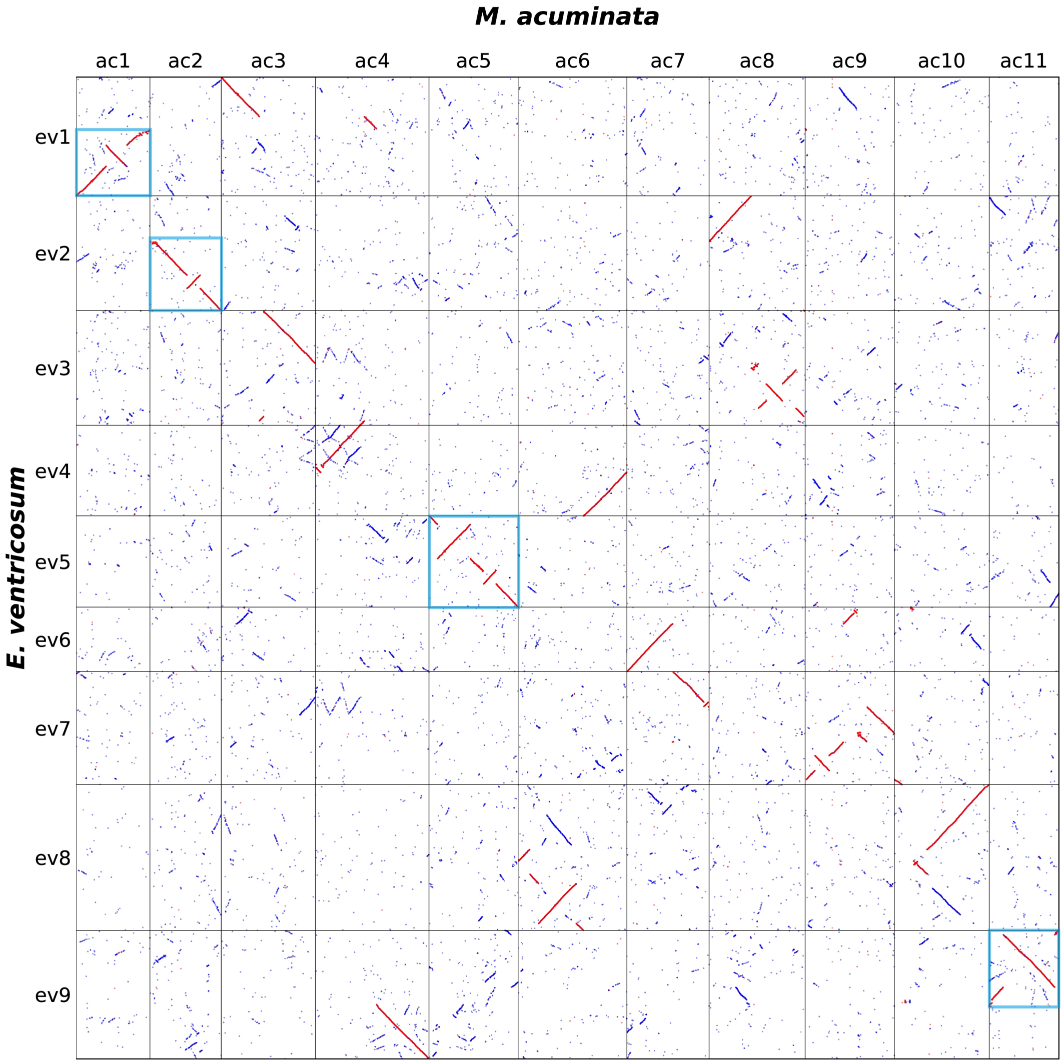

Comparing Mac with Eve reveals additional Shared or Nested patterns at a deeper phylogenetic level, allowing the identification of **4 protochromosomes** (blue boxes) for subsequent AMK reconstruction.

**Step 3: Reconstruct the Ancestral Karyotype Based on Fusion Points (RCT / EEJ)**

The core of this step is to reconstruct the composition of ancestral chromosomes from the protochromosomes identified above, by incorporating fusion points observed in modern chromosomes and expressing them in RCT or EEJ format. Special attention should be paid to cases where clade nodes correspond to ancestral chromosomes; a single representative species should be used to display all identified ancestral chromosomal regions to avoid omissions. Here we use *M. acuminata* (Mac) as the display species.

1. Assign distinct colors to each identified ancestral chromosomal segment (protochromosome / ancestral block).
2. Map these colored ancestral segments back onto the chromosomes of modern species to locate candidate fusion points.
3. Validate candidate fusion points using the outgroup *Canna indica* to determine which lineage most likely harbors each fusion or rearrangement event.

Using Mac (ac1, ac2, …) and Mex (ex1, ex2, …) as an example: chromosomes that display continuous collinearity with no apparent breakpoints (e.g., ac1, ac2) can be treated as relatively conserved ancestral segments. For chromosomes assembled from multiple segments, fusion points must be cross-referenced with Eve collinearity to distinguish "ancestral states" from "lineage-specific events."

In the Mex vs. Mac dot plot, a reciprocal translocation is inferred between ac7 and ac11: since ac1 and ac11 have already been confirmed as ancestral chromosomes, the second half of ac7 constitutes an ancestral chromosome of *Musa*; because this block is also intact in Ev, it is elevated to an AMK ancestral chromosome. By the same reasoning, the first half of ac7 is likewise an ancestral chromosome of *Musa*, and is similarly elevated to an AMK ancestral chromosome because it remains intact in Ev.

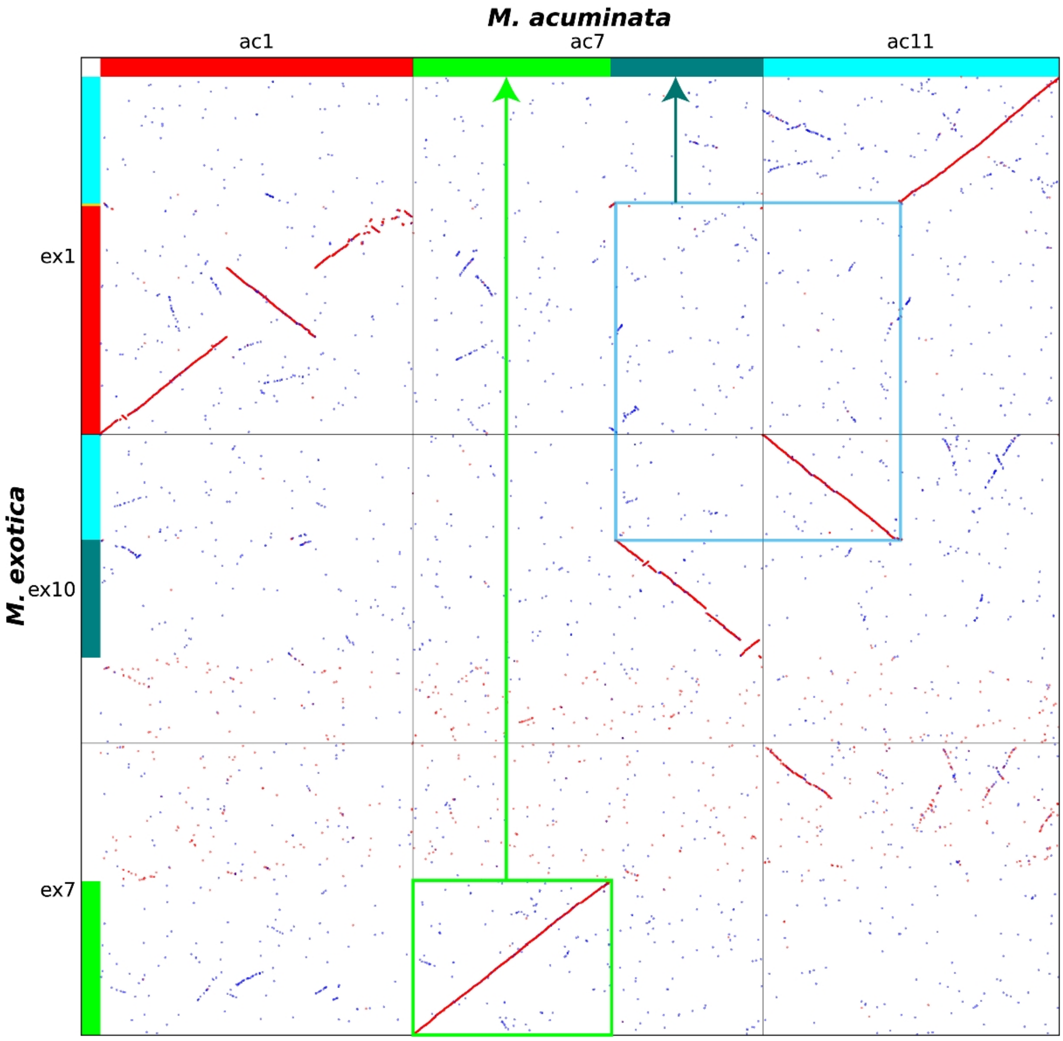

In the Mac vs. Eve comparison, the portion of ev9 remaining after removing the identified protochromosome constitutes an intact block within ac4, and ac4 is already an ancestral chromosome of *Musa*. For Ev, the formation of ev9 conforms to the EEJ pattern; the residual block can therefore be added as a new AMK ancestral chromosome, while the remaining region of ac4 can be incorporated as an additional protochromosome specific to the *Musa* lineage.

Following the same RCT / EEJ logic, multiple ancestral chromosomes can be progressively identified:

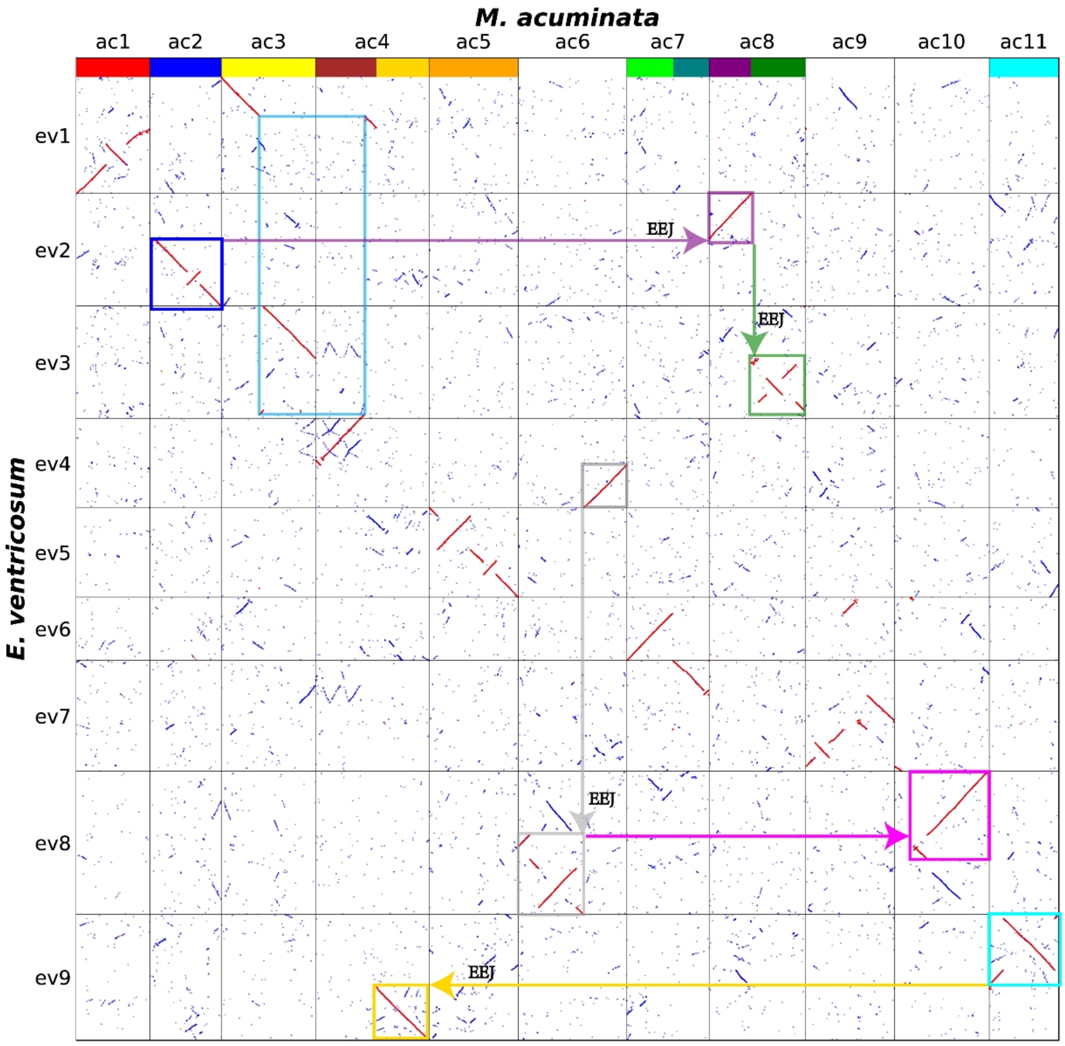

It should be noted that the upper half of ev4 represents a subset of a complete ancestral chromosome. We split the ancestral chromosome in the anterior portion of ac4 into two blocks at the fusion point and validated this with the outgroup *Canna indica*:

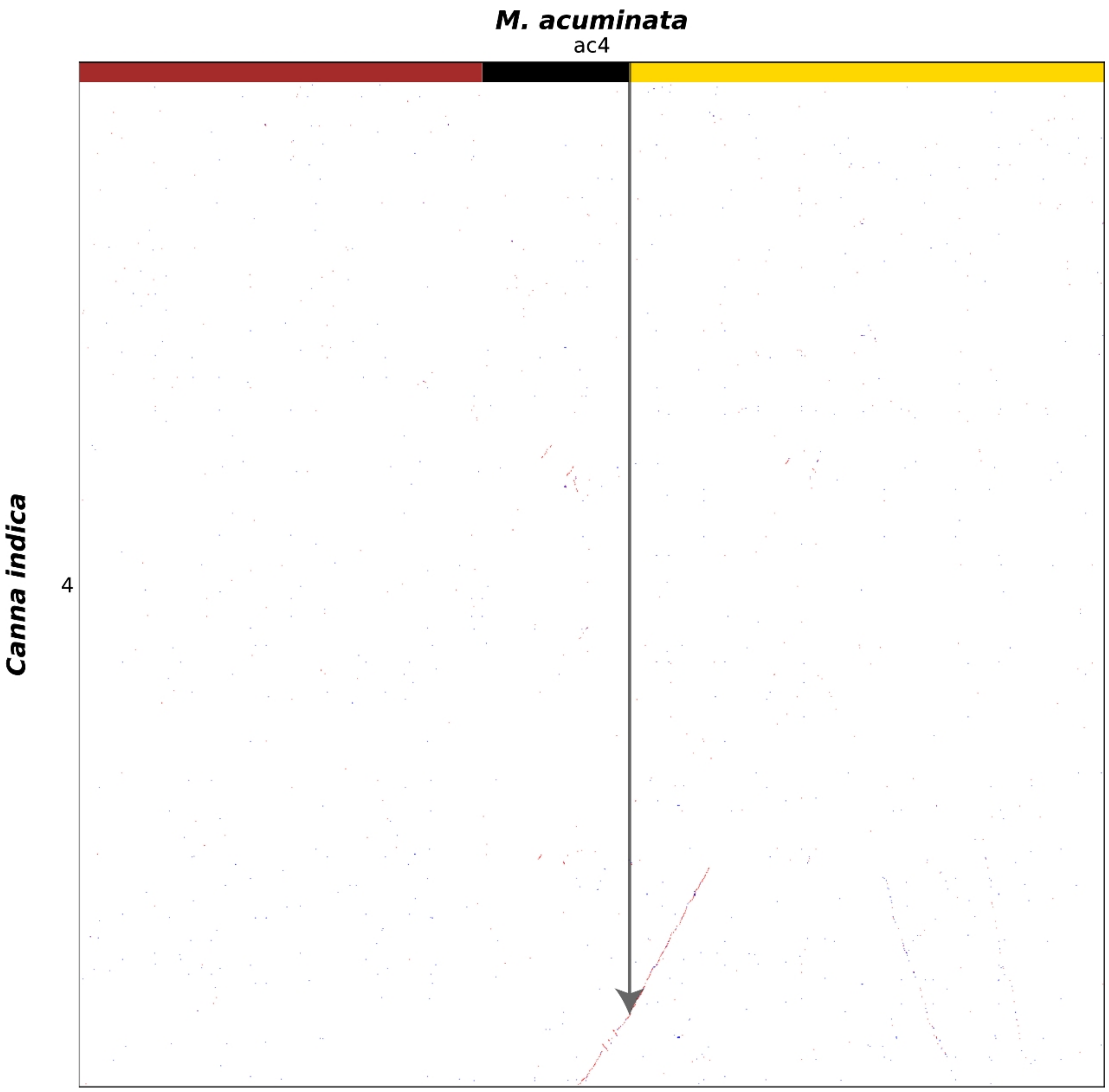

The validation shows that the two blocks are contiguous in *Canna indica*, suggesting that the formation of ev9 likely involved an RCT event in which almost no sequence from the ancestral chromosome ac11 was incorporated. Subsequently, ac6 is further divided into two ancestral chromosomes, and one additional ancestral chromosome is extracted from ac10.

**Step 4: Determine Ancestral Chromosomes from Remaining Segments Using Fusion Points**

Using outgroups as references, the combinatorial relationships among remaining segments are resolved to finalize the ancestral chromosomes.

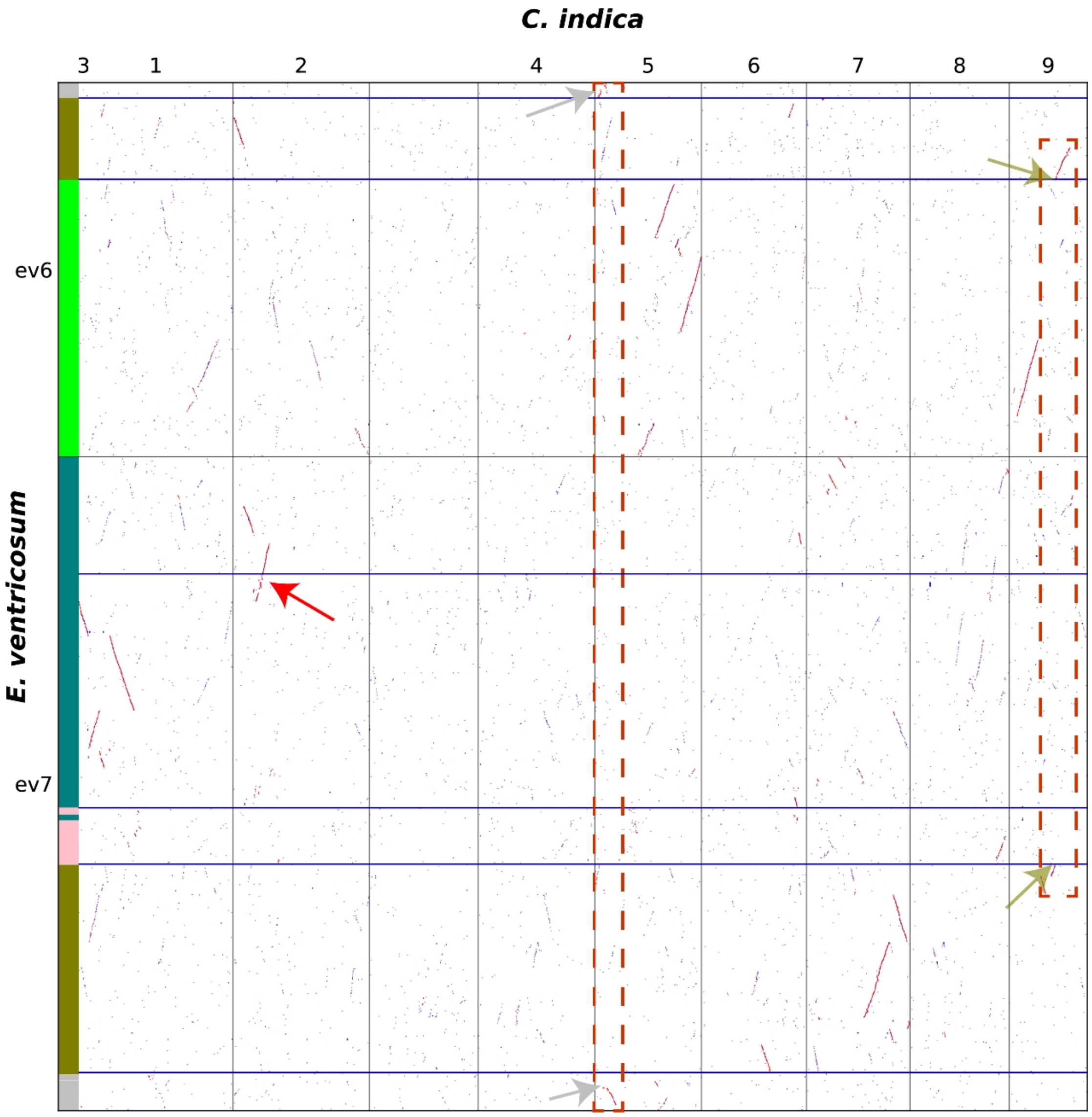

Through comparison of Eve with the outgroup *Canna indica*, the ancestral chromosomes corresponding to ev7 and ac9 can be confirmed to be in a joined state; the two foremost blocks of ev6 are confirmed to be in a separated state. Finally, the remaining regions are partitioned into multiple colored blocks based on breakpoint positions, and new ancestral chromosomes are assembled by combining blocks according to their correspondence relationships (indicated by dashed boxes and arrows), yielding the final AMK karyotype.

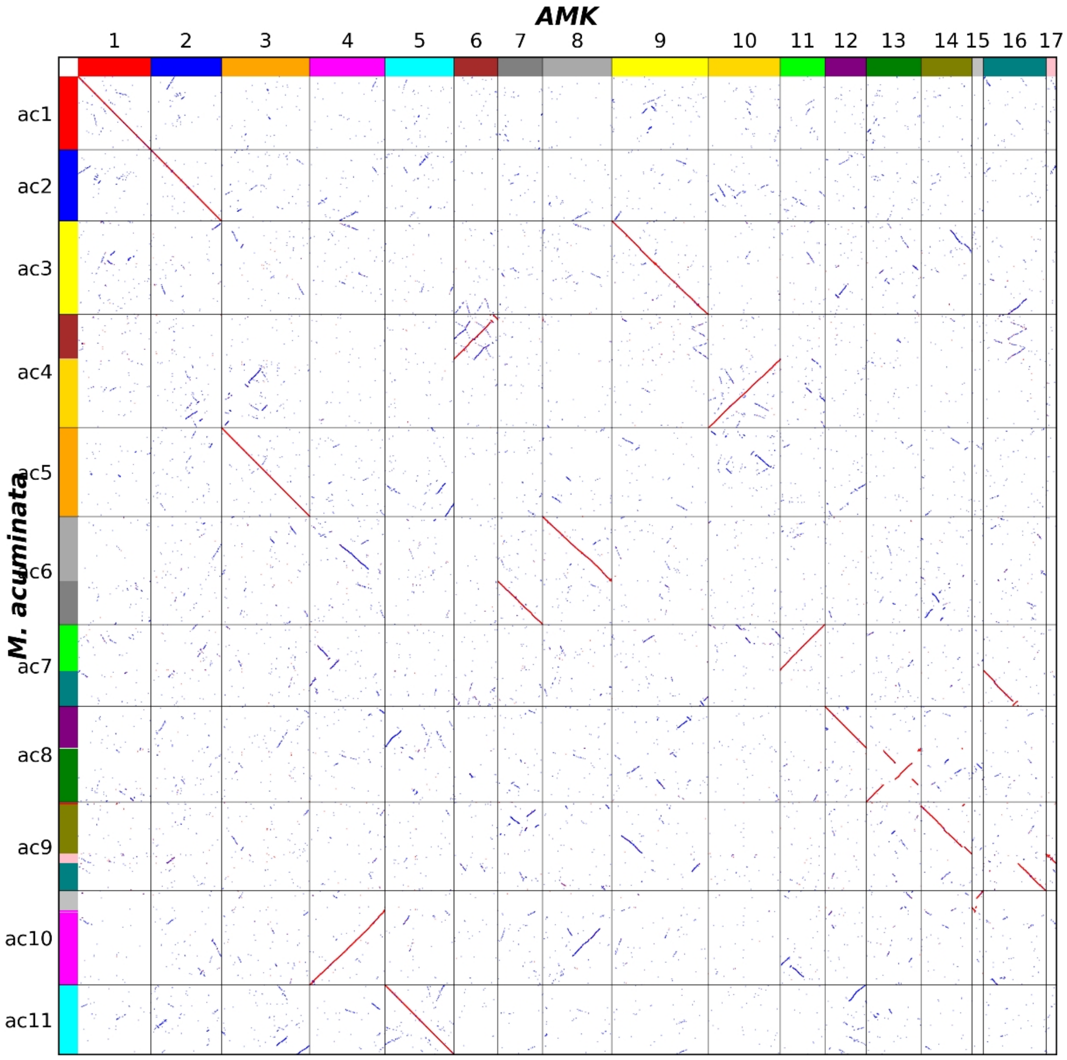

AMK
M. acuminata
ac1
ac2
ac3
ac4
ac5
ac6
ac7
ac8
ac9
ac10
ac11
